# Recurring daytime and nighttime modes of VOC emissions in a cool-temperate oak forest

**DOI:** 10.64898/2026.07.20.739470

**Authors:** Kanako Sekimoto, Yoshihisa Suyama, Yukiumi Kita, Daisuke Fukuyama, Abigail Koss, Asami Matsukami, Sota Tomihira, Hiroki Yamagishi, Kaori Shiojiri, Takuya Saito, Kazufumi Yazaki

## Abstract

Plants emit substantial amounts of biogenic volatile organic compounds (VOCs) that link plant physiological activity to ecological interactions and atmospheric chemistry. However, the processes regulating VOC emissions within the forest air and at the interface directly above the canopy remain poorly characterized. In this study, we investigated forest-scale VOC dynamics in a cool-temperate deciduous forest dominated by *Quercus crispula* using high-time- and high-mass-resolution proton-transfer-reaction time-of-flight mass spectrometry (PTR-ToF-MS) integrated with positive matrix factorization (PMF). This non-targeted, process-oriented framework was applied to forest-interior and canopy-top atmospheres during rain-free summer days in 2024 and 2025 to extract dominant modes of VOC emissions. The PMF consistently resolved two recurring modes characterized by the daytime and nighttime enhancement patterns. The daytime mode was dominated by isoprene and its oxidation products and demonstrated strong light- and temperature-dependence, whereas the nighttime mode was enriched in mono- and sesquiterpenes. The daytime contribution exhibited a pronounced morning–afternoon asymmetry, indicating non-linear physiological and canopy-scale controls. These patterns were reproducible across the years. This study demonstrates that combining PTR-ToF-MS with PMF enables robust, top-down identification of recurring modes of forest VOC variability within and above forest canopies, linking leaf-level physiology and ecosystem-scale atmospheric exchange.

**Highlight:** Forest-scale VOC emissions were resolved into recurring daytime and nighttime modes using a non-targeted PTR-ToF-MS and PMF framework.

## 1. Introduction

Plants emit a vast diversity of volatile organic compounds (VOCs), comprising an estimated tens of thousands of chemical species (Peñuelas and Llusià, 2004). Globally, biogenic VOC emissions exceed 10^9^ tons of carbon equivalents per year, representing a substantial fraction (up to approximately 10 %) of the carbon fixed by photosynthesis and subsequently released back into the atmosphere (Guenther *et al*., 1995; Peñuelas and Llusià, 2004). This substantial carbon investment highlights the fundamental importance of VOC emissions during plant life. Plant-derived VOCs perform numerous physiological and ecological functions (Peñuelas and Llusià, 2003; Sasaki *et al*., 2007; Vickers *et al*., 2009). Simultaneously, biogenic VOCs provide a major link between terrestrial ecosystems and atmospheric chemistry. Highly reactive compounds, such as isoprene, monoterpenes, and sesquiterpenes, strongly affect air quality by contributing to ozone formation, secondary organic aerosols, and cloud condensation nuclei, thereby influencing air quality and climate (Kavouras *et al*., 1998; Crounse *et al*., 2013; Ehn *et al*., 2014; Sharkey and Monson, 2017; Wennberg *et al*., 2018).

Experimental and observational approaches have been developed to quantify and interpret these substantial biogenic VOC emissions. At the ecosystem scale, flux measurements fundamentally constrain the net VOC exchange between forests and the atmosphere (Guenther *et al*., 1995; Karl *et al*., 2009). At the laboratory scale, leaf enclosure techniques and growth-chamber experiments have yielded comprehensive insights into species-specific emission capacities and physiological regulation under controlled conditions (Guenther *et al*., 2012). These approaches account for only a fraction of the overall fate of VOCs, from leaf surface release to their transport and transformation within the forest air and atmosphere. Flux measurements are integrated over large spatial domains, obscuring vertical gradients and short-term variability within forest canopies, whereas leaf-surface and chamber-based experiments isolate plants from their natural ecological and atmospheric contexts. Consequently, VOC variability within forest air, from the forest interior to the atmosphere directly above the canopy, remains poorly characterized. VOC dynamics within forest air, particularly near the forest floor and canopy, are closely associated with biotic interactions involving plants, insects, and rhizosphere-associated microorganisms (Holopainen and Gershenzon, 2010; Peñuelas *et al*., 2014). Numerous VOCs mediate plant–plant, plant–insect, and plant–microbe interactions, and their concentrations and compositions can vary notably with height, time of day, and local physiological activity. Therefore, resolving this variability is essential for understanding how biological interactions within forests are coupled with canopy-scale chemical dynamics.

Additional challenges result from analytical limitations. Measurements of biogenic VOCs have traditionally relied on gas chromatography coupled with electron-ionization mass spectrometry (GC/EI-MS), providing compound-resolved information for target species such as isoprene and monoterpenes (Tholl *et al*., 2006; Ortega and Helmig, 2008). Although this bottom-up, compound-focused approach has yielded fundamental insights into plant VOC emissions, it focuses on a limited subset of compounds within chemically complex forest atmospheres. Natural forest air contains diverse mixtures of VOCs derived from plants, insects, microbes, and in-canopy chemical processes. Therefore, targeted analyses risk overlooking non-target compounds that may contribute to dominant patterns of variability. Additionally, GC/EI-MS measurements are typically based on discrete sampling, limiting their ability to resolve rapid temporal dynamics and diurnal variability (Niederbacher *et al*., 2015). Therefore, observational frameworks that combine broad chemical coverage with high temporal resolution are required to capture process-level variability within and above forest canopies. These limitations highlight the need for non-targeted, high-resolution approaches to elucidate the emergent patterns of VOC organization in forest ecosystems.

In this study, we combined high-time-and high-mass-resolution measurements using proton-transfer-reaction time-of-flight mass spectrometry (PTR-ToF-MS) with positive matrix factorization (PMF) to investigate VOC emissions within a cool-temperate deciduous forest dominated by *Quercus crispula*. Compared with boreal and coniferous forest ecosystems, where forest-scale VOC measurements are relatively abundant, for example, in vertical distribution studies of spruce and pine stands (Rinne *et al*., 2016; Li *et al*., 2020, 2021; Petersen *et al*., 2023), observational data from temperate deciduous forests remain limited despite their wide geographic extent and ecological importance. Studies on mixed deciduous forests that capture VOC fluxes both above and below the canopy, including contributions from the forest floor, are rare (Stoy *et al*., 2021; Dumont *et al*., 2026). The PTR-ToF-MS plus PMF approach provides a non-targeted, process-oriented framework that enables the extraction of the dominant modes of VOC emissions without requiring a priori compound selection or complete compound-level identification. We hypothesized that a complex mixture of VOCs in forest air can be characterized by a limited number of recurring modes that represent distinct environmental and physiological conditions. To test this hypothesis, we applied this approach to forest-interior and canopy-top atmospheres and identified recurring daytime and nighttime patterns in VOC composition, which were then related to physiological and micrometeorological drivers. Through this analysis, we aimed to provide a forest-scale perspective on VOC dynamics and explore the links between leaf-level physiological processes and ecosystem-scale atmospheric exchanges.

## 2. Materials and methods

### 2.1. Measurement site description and field setup

To characterize forest-scale VOC dynamics under natural conditions, this study was conducted at the Shirakami Natural Science Park (40° 31’ 7” N, 140° 12’ 54” E, 245 m above sea level), a research forest managed by Hirosaki University in Aomori Prefecture, northern Japan. This region is characterized by a cool-temperate climate. In 2024, the mean air temperatures of the warmest month (August) and coldest month (January) were 23.4 and –1.1 ℃, respectively, and the annual precipitation was 2469 mm. The nearest settlement, Nishimeya Village (population of approximately 1000), is located approximately 10 km from the study site, indicating a low local population density and minimal influence from anthropogenic and industrial emissions.

The forest is composed of secondary cool-temperate deciduous broad-leaved vegetation dominated by *Quercus crispula*, with a well-developed understory. The major shrub and sub-canopy species include *Lindera umbellata* and *Clethra barbinervis*, whereas the evergreen dwarf shrubs include *Aucuba japonica* and *Ilex leucoclada* (Yamagishi and Ishikawa, 2012). A vegetation survey conducted in 2024 within a 25 × 45 m plot including the sampling port location showed that *Q. crispula* accounted for 54 % of the total basal area among trees with a diameter at breast height (DBH) ≥ 8 cm. Increment core analyses of several *Q. crispula* individuals indicated stand ages of approximately 60 years, with an average canopy height of approximately 15 m.

Several adaptations were implemented in the field setup to enable long-term real-time measurements of VOC emissions under forest conditions. First, a proton-transfer-reaction time-of-flight mass spectrometry (PTR-ToF-MS) unit was installed in an air-conditioned building to ensure stable operation of the instrument during extended field campaigns. At the same time, the building needed to be sufficiently close to the sampling locations to allow real-time measurements through a 50 m PTFE sampling line. Second, we secured a clean power supply without using generators that emit exhaust fumes. Third, a stable Internet connection was established to enable remote monitoring of instrument status and routine calibration. To meet these criteria, we selected the *Quercus crispula* forest in the Shirakami area, which is managed by Hirosaki University, as our experimental site.

### 2.2. Micrometeorological observations

Meteorological variables, including temperature, relative humidity, precipitation, snow depth, wind speed, and solar radiation, were measured continuously on-site using an automated meteorological system installed on a tower 10 m above ground level (Ishida, 2012). Continuous meteorological observations have been conducted since 2010, with a maximum temporal resolution of 10 min, and were used to characterize environmental drivers of VOC emissions, including forest-scale temperature response (‘*C_T_*’ used in Section 3.5).

In addition to long-term meteorological measurements, photosynthetically active radiation (PAR) was measured during the intensive VOC observation periods at air sampling heights (9 and 15 m above ground level). PAR sensors (MIJ-14 Type2, Environmental Measurement Japan, Fukuoka, Japan) were installed adjacent to each air sampling port to capture the local light conditions within the forest interior and at the canopy top. These measurements were recorded at a time resolution of 1 s, synchronized with the PTR-ToF-MS observations, and were used to derive the light-response term (‘*C_L_*’ used in Section 3.5) at each sampling height.

### 2.3. VOC measurements

To resolve the temporal variability of VOC emissions within and above the forest canopy, VOC measurements were conducted from August 29 to September 3, 2024 and from August 23 to September 7, 2025. Ambient forest air was sampled at two heights (9 and 15 m above ground level), corresponding to the forest interior and immediately above the canopy top, respectively (Supplementary Fig. S1). Within a 20 m radius of the canopy-top sampling port, *Q. crispula* accounted for approximately 80 % of the canopy leaf area. Air was drawn through two polytetrafluoroethylene (PTFE) tubes mounted on a rolling tower (approximately 10 m height) and extension poles (approximately 6 m length) using a microdiaphragm gas pump (NMS030. 1.2KPDC-B4, KNF, Germany). Each sampling tube was 50 m long, with outer and inner diameters of 6.35 and 4.35 mm, respectively. The flow rate through each tube was maintained at 3 L min^-1^, resulting in an air residence time of ˂10 s prior to reaching the analytical instrumentation. Sampling was conducted sequentially at each height for 5 min within a 40-min measurement cycle using a VICI Valco 1/4” multiposition valve system (GL Science, Japan). Several modifications were implemented to enable long-term field measurements, including the placement of a Teflon cover at the tube inlet to protect against rain, wind, and insects.

To achieve high-time-resolution characterization of VOC composition, a subsample of the air flow (0.2 L min^-1^) was analyzed using proton-transfer-reaction time-of-flight mass spectrometry (PTR-ToF-MS; custom Vocus Scout CI-TOF, TOFWERK AG, Thun, Switzerland), installed in a temperature-controlled laboratory. PTR-ToF-MS detects VOCs through proton-transfer reactions between H_3_O^+^ reagent ions and compounds with proton affinities higher than that of water, followed by high-resolution time-of-flight mass analysis of the resulting protonated molecules (Lindinger *et al*., 1998). The instrument consisted of a discharge source producing H_3_O^+^ reagent ions, a focusing ion-molecular reactor (FIMR) for proton-transfer ionization of VOCs by H_3_O^+^ which was operated at an electric field-to-number density ratio (*E/N*) of approximately 130 Td (1 Td = 10^-17^ V cm^-2^), a quadrupole ion guide, and a time-of-flight mass analyzer with a mass resolving power of approximately 4500 (*m*/Δ*m*, FWHM) at *m/z* 107. The configurations of the discharge source and the FIMR were equivalent to those described by Krechmer *et al*. (2018). Mass spectra were recorded at a time resolution of 1 s. Instrument background measurements were automatically performed every 320 min by introducing clean air generated using Vocus Zero Air (TOFWERK AG, Thun, Switzerland). Mass calibrations were conducted daily using a multicomponent standard cylinder containing approximately 1 ppmv each of acetonitrile, acetaldehyde, 1,2,4-trimethylbenzene, *m*-xylene, toluene, α-pinene, acrylonitrile, benzene, isoprene, methanol, and methyl vinyl ketone purged with pure N_2 (_NIPPON SANSO, Tokyo, Japan).

Mass spectra were recorded over the *m/z* range of 30-231, covering VOCs from small oxygenated compounds such as methanol and acetaldehyde to larger compounds associated with sesquiterpenes and their oxygenated derivatives. Elemental compositions were determined based on a combination of a limited set of atoms, such as H, C, O, N, and S atoms, and the high mass resolution of the ToF analyzer. The PTR-ToF-MS mass spectral data were processed using the Igor-based software package Tofware (version 3.2.5; TOFWERK AG).

### 2.4. Gas chromatographic analysis

Gas chromatographic (GC) separation was performed using a Sylph system (BallWave, Miyagi, Japan) equipped with a polyethylene glycol column (PEG20M; 30 m length, 0.25 mm inner diameter, 0.25 μm film thickness). Hydrogen was used as the carrier gas at a flow rate of 1 mL min^-1^. The oven temperature program was 50 °C (maintained for 5 min), increased to 90 °C at 5 °C min^-1^, then to 180 °C at 10 °C min^-1^, and finally maintained for 3 min. For GC analyses, 0.2 L of air sampled at the canopy-top height (15 m) was collected over a 10-min period. The GC outlet was connected to the PTR-ToF-MS inlet via a glass capillary (0.53-mm inner diameter; 60 cm length). The GC effluent was introduced into the PTR-ToF-MS together with ambient room air in the laboratory.

### 2.5. Concentration calculations

For the selected ions, ion intensities (in counts-per-second ‘cps’) were converted to ambient concentrations (in ppbv) by correcting for the quadrupole transmission efficiency, mass-dependent duty cycle of ToF extraction, hydrated cluster formation, and fragmentation. For the GC-separated data, when multiple chromatographic peaks were observed at a given *m/z*, including contributions from species other than the target VOC, raw ion intensities were apportioned based on the relative areas of the corresponding GC peaks. Corrected ion intensities were then converted to atmospheric concentrations using estimated proton-transfer reaction rate constants and the instrument functions obtained from the multicomponent standards described above, with an associated uncertainty of ˂50 % (Sekimoto *et al*., 2017).

### 2.6. PMF analysis

To interpret the VOC ion data acquired by PTR-ToF-MS, we applied positive matrix factorization (PMF; Paatero and Tapper, 1994; Paatero, 1997; Ulbrich *et al*., 2009). PMF is a bilinear unmixing model in which the data matrix is represented as a linear combination of a set of ‘factors’ with constant factor profiles, mass spectra (MS) of VOC ions, whose contributions vary over time (time courses, TCs):

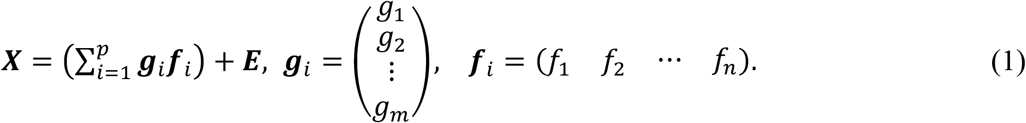

where ***X*** is an *m* × *n* matrix containing the measured data to be fit. In the present analysis, the *n* columns of ***X*** corresponded to the MS, and the *m* rows corresponded to the TC of each ion acquired at a 1-s resolution across individual sampling ports. *p* is the number of factors. The *m*-dimensional column vectors, {***g****_i_*}(*i* = 1 ∼ *p*), are the factor time course, while the *n*-dimensional row vectors, {***f****_i_*}(*i* = 1 ∼ *p*), are the factor mass-spectral profiles. The *m* × *n* matrix ***E*** contains the residuals not explained by the model, that is. ***E*** ≡ ***X – (Σ^p^_i=1_ g_i_ f_i_)***. A schematic representation of factorization is shown in Supplementary Fig. S2. PMF does not require a priori information regarding factor profiles or time courses. The optimal number of factors was determined based on residual structure, factor interpretability, and inter-factor correlations.

PMF was performed using the Source Finder (SoFi) interface and a multilinear engine (ME-2) (Paatero, 1999; Canonaco *et al*., 2013), applied to time courses of background-subtracted ion intensities (in counts-per-second) for selected ions. PMF was applied to 535 ions selected from the PTR-ToF-MS dataset (Supplementary Table S1). Ions were retained when they were monoisotopic, resolved from neighboring peaks, exhibited a signal-to-noise (S/N) ˃3, and showed enhancement above the instrumental background at least at one sampling position. Primary reagent ions (H_3O_^+^[H_2O_]_0-2_) and ions associated with instrumental contamination, including PTFE fragments and transition metals, were excluded.

## 3. Results

Weather conditions varied considerably during the study period, with notable differences between sunny (including partly cloudy) and rainy periods. Because rainfall strongly influences soil moisture and consequently VOC emissions, data collected within 24 h of precipitation events were excluded from subsequent analyses. The following sections focus on VOC observations during rain-free periods, beginning with an overview of the VOC ions selected for PMF analysis.

### 3.1. VOC ion composition and diel variability

The 535 ions mainly consisted of hydrocarbons (HC; 112 ions), oxygenates (O-HC; 316 ions), and nitrogen-and oxygen-containing hydrocarbons (N, O-HC; 58 ions). These ions accounted for 98 % of the measured total ion intensities in both the forest-interior and canopy-top atmospheres in 2024 (Fig. 1A inner circle). Approximately 67 % of the total intensity was accounted for by the 14 major ions displayed by their elemental compositions in Fig. 1A (outer circle). ‘no ID’ indicates ions that were not identified with mass accuracy within 20 ppm based on the above criteria and thus may include atoms other than H, C, O, N, and S.

**Fig. 1.**
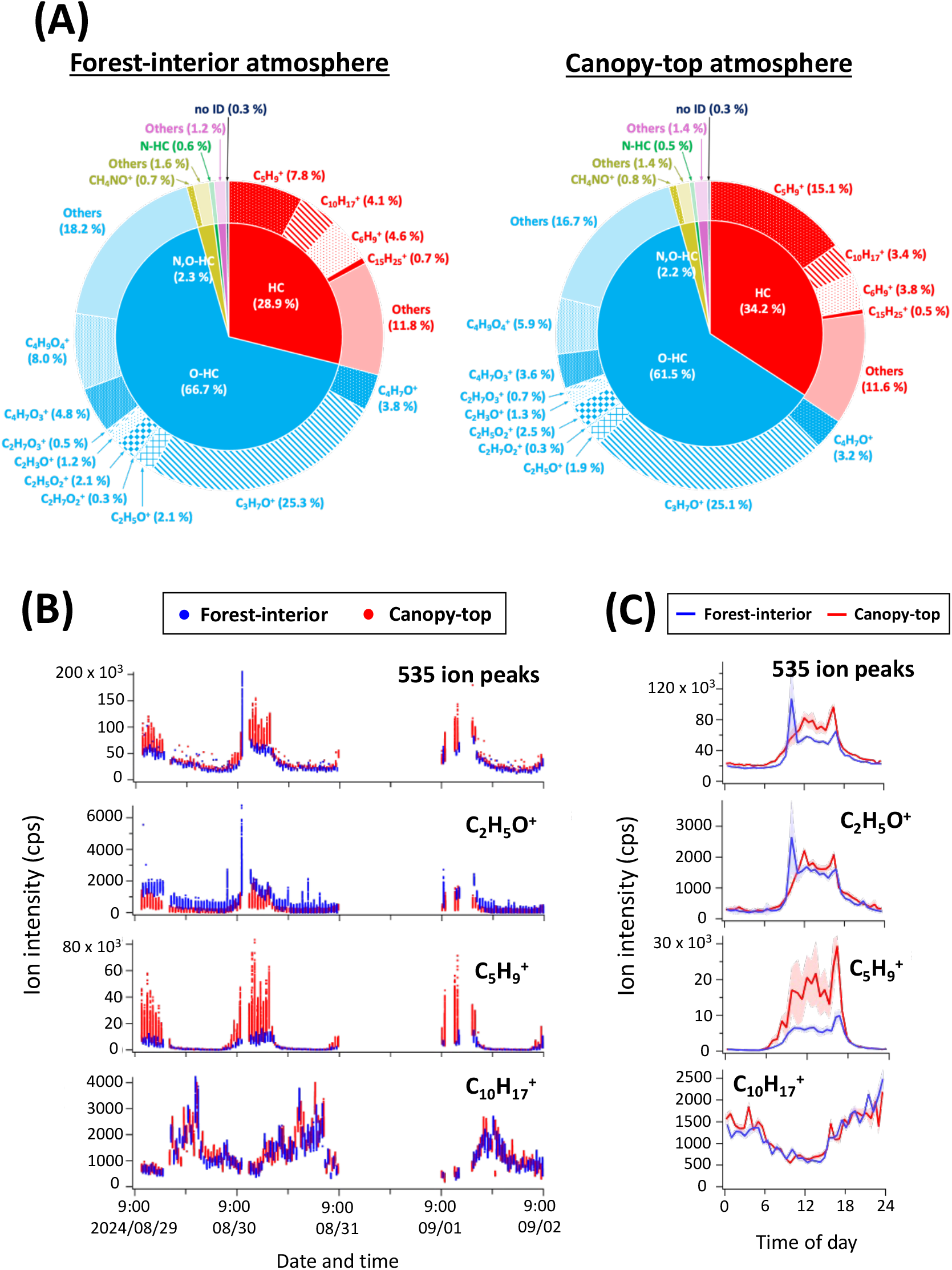
VOC ions detected using PTR-ToF-MS in the forest-interior and canopy-top atmospheres in 2024. (A) Fractions of VOC ions contributing to the total measured ion intensities (cps) during three rain-free days. The inner circle in each pie chart indicates the elemental composition of the ions, and the outer circle lists specific ion formulae. (B) Real-time monitoring of selected VOC ions during three rain-free days and (C) their average diurnal cycles. In (C), solid lines and shaded areas indicate the mean values and corresponding standard deviations, respectively.

Fig. 1B and C show the real-time monitoring of selected VOC ion signals originating from the forest-interior and canopy-top atmospheres over 36 h, together with their average diurnal cycles. Total VOC ion intensities (defined as the sum of counts-per-second ‘cps’ over all 535 ions) were relatively higher during daytime than at night, with average daytime-to-nighttime ratios of approximately 2–3, calculated from the mean values during 12:00-15:00 and 00:00-03:00 (Fig. 1C). The temporal behaviors of individual VOC ion signals were broadly classified into three categories.

i. signals showing limited diurnal variation or modest daytime enhancement (1 ≤ daytime-to-nighttime ratios ≤ 5), such as C_2_H_5_O^+^;
ii. signals that were significantly enhanced during the daytime (daytime-to-nighttime ratios > 5), such as C_5_H_9_^+^.
iii. signals that were higher during the nighttime (daytime-to-nighttime ratios < 1), that is,C_10_H_17_^+^

In the following analysis, the PMF was applied separately to the forest-interior and canopy-top datasets to parameterize the variability in the emitted VOC ion intensities.

### 3.2. Identification of dominant modes of VOC emissions

#### 3.2.1. Recurring daytime and nighttime modes in the forest interior

PMF was applied to the PTR-ToF-MS dataset consisting of 535 ions selected from the forest-interior atmospheres during the three rain-free days in 2024. The number of factors tested ranged from one to six. Based on an analysis of the residuals from each factor solution and inter-factor correlation, a two-factor parameterization was determined to be the optimal model for capturing the variability in the total ion intensities of the emitted VOCs (Fig. 2A). Residuals, defined as the difference between the measured total ion intensities and those calculated from the PMF fits, were ˂16 %. Although only 165 of the 535 ions were fitted within 50 % of their measured intensities, these ions accounted for 94 % of the total signal. The other 370 ions were poorly fitted because the difference between their measurements and the PMF reconstruction was higher than 50 %. Poorly fitted ions were generally those that contributed ˂0.3 % of the total intensity, consistent with previous findings (Ulbrich *et al*., 2009), suggesting that the dominant factors did not capture their variability.

**Fig. 2.**
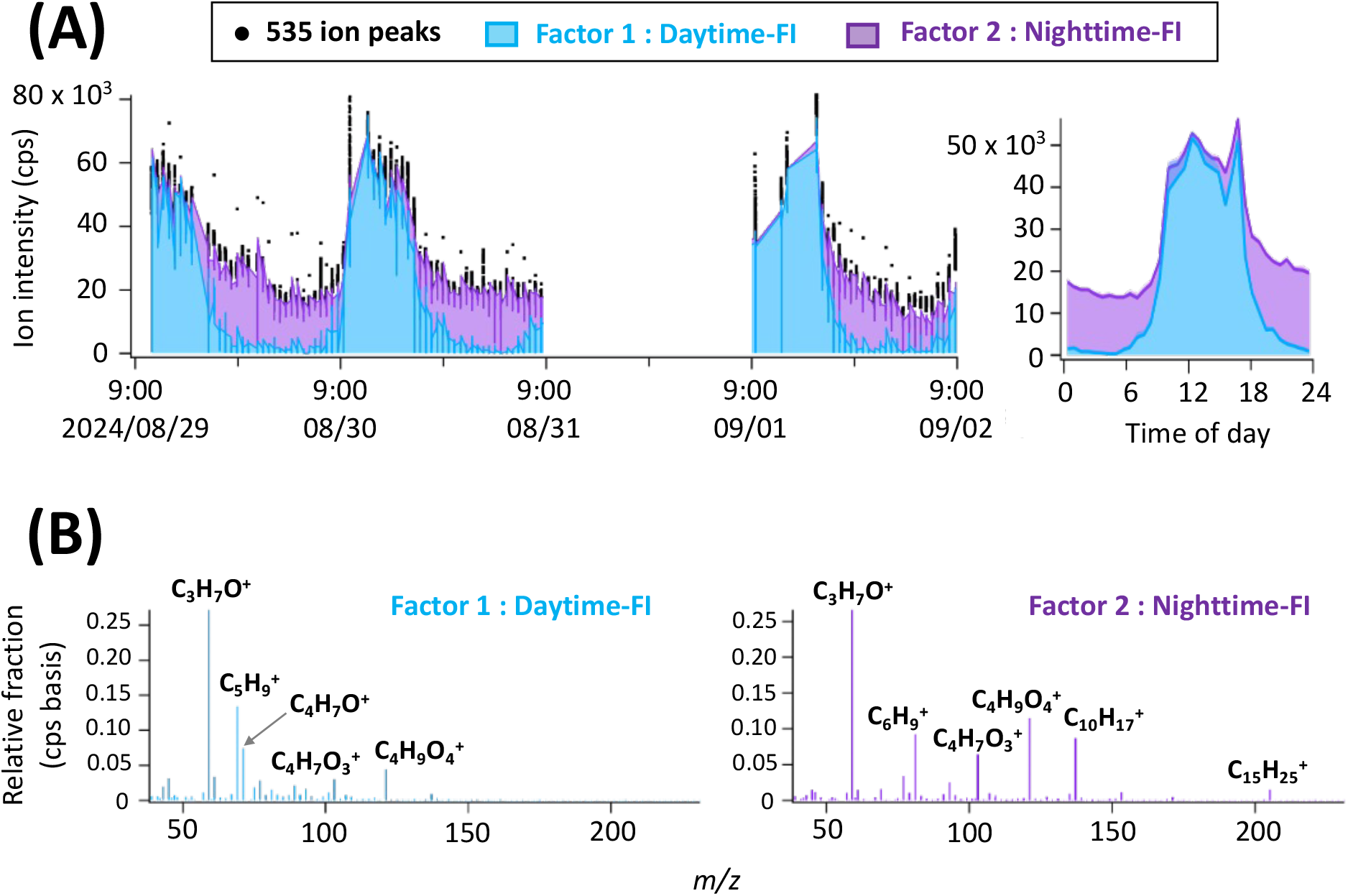
PMF results for the two-factor solution applied to VOC ion signals measured in forest-interior atmospheres during three rain-free days in 2024. (A) Time courses of 535 measured ion signals and PMF fits (left) and average diurnal cycles of PMF fits (right). Individual PMF fits (blue and purple) are shown as stacked rather than overlapped. (B) Mass spectral profiles of the daytime and nighttime forest interior (FI) factors.

The time courses for the first and second factors exhibited significant and recurring enhancements during the day and night, respectively (Fig. 2A). Accordingly, these factors are referred to as ‘daytime-FI’ and ‘nighttime-FI’ throughout this study (‘FI’ represents forest-interior). The mass spectral profiles of the daytime-and nighttime-FI factors are shown in Fig. 2B, and the relative fractions of individual ions in each factor are presented in Supplementary Table S1. Both profiles were dominated by C_3_H_7_O^+^; however, notable differences indicated the distinct variability patterns of the VOC ion intensities. The relative fractions of C_5_H_9_^+^ in the daytime-and nighttime-FI factors were 0.134 and 0.016, respectively; 0.009 and 0.087 for C_10_H_17_^+^; and 0.032 and 0.015 for C_2_H_5_O^+^, which are consistent with the results shown in Fig. 1. Further interpretations of these compositional differences and their relationships with environmental factors, particularly light intensity and temperature, are provided in Sections 3.3 and 3.5.

The optimal PMF model was determined based on the residual analysis and inter-factor correlation. residual for the one-factor solution was 27.1 %, whereas those for the two-, three-, four-, five-, and six-factor solutions were 15.8 %, 12.8 %, 11.4 %, 10.3 %, and 9.3 %, respectively. These results indicated that the PMF fit improved substantially from one to two factors; however, changes beyond these two factors were minimal, indicating that a two-factor solution was sufficient to describe the dominant variability in the measured VOC ion intensities. When more than two factors were incorporated, the time courses and mass spectral profiles of the additional factors largely represented the ‘splitting’ of the nighttime factor. Supplementary Fig. S3 shows the correlation between *n*-factor solutions (*n* = 3 and 4) and the PMF results from the daytime and nighttime factors for the same datasets as shown in Fig. 2. These results indicate that only two factors, daytime-and nighttime-FI, were required to explain most of the variability in VOC ion intensities from forest-interior atmospheres.

#### 3.2.2. Similar modes above the canopy

The PMF was applied to the PTR-ToF-MS dataset from the canopy-top atmospheres. The number of factors tested ranged from one to six. Based on the residual analysis and inter-factor correlation, most of the variability in VOC ion intensities from the canopy-top atmospheres was explained by two factors representing diurnal changes (Fig. 3A). The overall residuals were approximately 16 %.

**Fig. 3.**
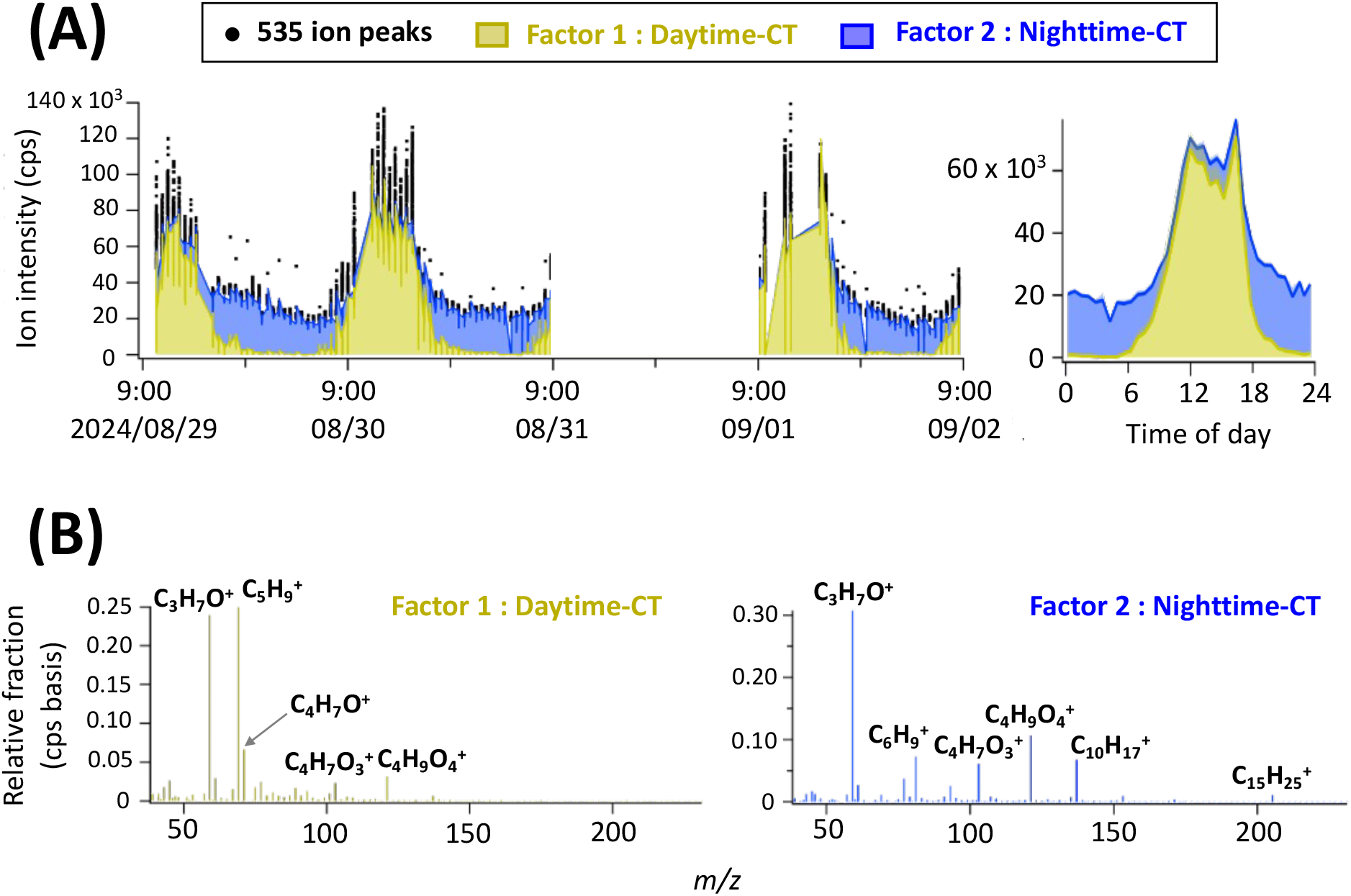
PMF results for the two-factor solution applied to VOC ion signals measured in canopy-top atmospheres during three rain-free days in 2024. (A) Time-courses of 535 measured ion signals and PMF fits (left), and average diurnal cycles of PMF fits (right). Individual PMF fits (yellow and dark blue) are shown as stacked rather than overlapped. (B) Mass spectral profiles of the daytime and nighttime canopy top (CT) factors.

The two factors are referred to as ‘daytime-CT’ and ‘nighttime-CT’ (‘CT’ represents canopy-top). The mass spectral profiles for the individual factors were partially similar to those from the forest-interior atmospheres (daytime-and nighttime-FI; Fig. 3B). Both profiles were dominated by C_3_H_7_O^+^. The relative fractions of C_10_H_17_^+^ and C_2_H_5_O^+^ in the daytime CT were 0.007 and 0.026, and those in the nighttime CT were 0.068 and 0.017, respectively. However, a notable difference was the significant enhancement of C_5_H_9_^+^, whose relative fraction was 0.250 in this case, which was almost double that of the forest-interior atmosphere (0.134). This result is consistent with the VOC emissions from the canopy dominated by *Q. crispula* foliage, as discussed in the next section.

### 3.3. Chemical characteristics of daytime and nighttime modes

The chemical characteristics of the mass spectral profiles of daytime-FI, daytime-CT, nighttime-FI, and nighttime-CT factors were examined (Figs. 2B and 3B), and multiple VOC ions were identified. Fig. 4A shows the VOC composition in the mass spectral profiles of daytime-and nighttime-FI factors from the forest-interior atmospheres. Both factors were dominated by oxygenates, followed by hydrocarbons and N-and O-containing hydrocarbons, with total fractions exceeding 0.98. The 14 ions highlighted in Fig. 1A accounted for approximately half of the total fraction (0.68 for daytime FI and 0.71 for nighttime FI), consistent with the raw measurement data. However, the degree to which each ion contributed to daytime-and nighttime-FI factors varied among species. The contributions of the ions to the daytime and nighttime factors are examined in the next section.

**Fig. 4.**
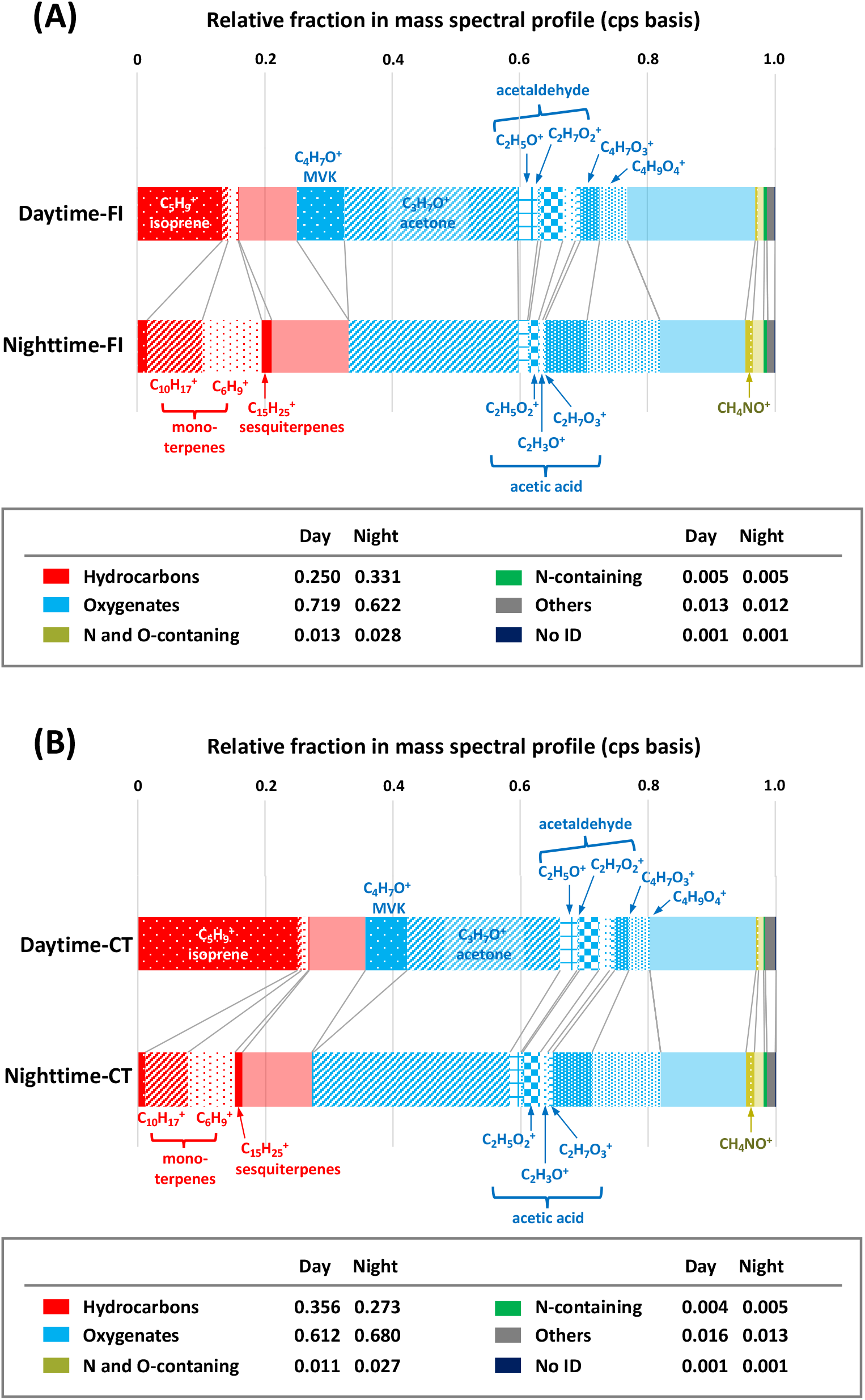
VOC ion composition of the mass spectral profiles of the daytime and nighttime factors obtained from (A) forest-interior (FI) atmospheres and (B) canopy-top (CT) atmospheres in 2024.

The canopy-top atmospheric results (Fig. 4B) were broadly similar to those from forest-interior atmospheres regarding the major contributors to daytime or nighttime factors. However, the fraction of C_5_H_9_^+^ in the daytime factor increased to approximately twice the value observed in the forest-interior atmospheres.

Gas chromatographic separation conducted prior to PTR-ToF-MS analysis confirmed the identities of several major ions (Supplementary Fig. S4). Based on the chromatographic retention times, the dominant ions C_5_H_9_^+^, C_4_H_7_O^+^, C_3_H_7_O^+^, and C_2_H_5_O^+^ were identified as protonated isoprene, protonated methyl vinyl ketone (MVK), protonated acetone, and protonated acetaldehyde, respectively. The C_2_H_7_O_2_^+^ ion exhibited chromatographic peaks at retention times identical to those of C_2_H_5_O^+^, indicating that C_2_H_7_O_2_^+^ was a hydrated cluster of protonated acetaldehyde formed in the PTR-ToF-MS reaction chamber. The C_10_H_17_^+^ ion exhibited two distinct chromatographic peaks. One peak corresponded to the retention time of α-pinene, whereas the other remained unidentified. As expected, C_6_H_9_^+^, a well-known fragment ion of protonated monoterpenes in PTR-MS (Müller *et al*., 2009), exhibited retention times identical to those of C_10_H_17_^+^, indicating that C_6_H_9_^+^ originated from monoterpene fragmentation in this dataset. These identified compounds have been widely reported in previous forest-atmosphere VOC studies. Isoprene emissions from *Q. crispula* leaves have been documented in Japan (Tani and Kawawata, 2008), and MVK is a well-known isoprene oxidation product (Atkinson, 1990; Warneke *et al*., 2001). MVK can also be formed as an artifact of other isoprene oxidation products (isoprene hydroxy hydroperoxides) during GC and/or PTR-MS analysis (Rivera-Rios *et al*., 2014). This artifact formation may occur under the present experimental conditions such that the detected C_4_H_7_O^+^ signals may include contributions from authentic forest emissions as well as from instrument-related processes, which cannot be distinguished in the present dataset. Acetone and acetaldehyde are commonly emitted by various tree species and produced in the atmosphere via VOC oxidation. Monoterpenes, represented by α-pinene in our experiment, are major VOCs emitted from forest vegetation. At the present forest site, *Magnolia obovata*, *Lindera umbellata*, *Larix kaempferi*, *Hydrangea petiolaris*, and *Sasa senanensis* were identified as the dominant monoterpene emitters, as confirmed by positioning the sampling port in close proximity to individual plants.

C_4_H_7_O_3_^+^ and C_4_H_9_O_4_^+^ were eluted as distinct chromatographic peaks at identical retention times (data not shown), indicating that the two ions originated from the same precursor VOC and that one of them likely represented an instrumental artifact. C_4_H_9_O_4_^+^ may represent a hydrated cluster of C_4_H_7_O_3_^+^ (protonated C_4_H_6_O_3_), whereas C_4_H_7_O_3_^+^ may represent a dehydrated fragment of C_4_H_9_O_4_^+^ (protonated C_4_H_8_O_4_). However, their chemical identities could not be assigned unambiguously. The other major ions present in the PTR-MS analysis (C_15_H_25_^+^, C_2_H_5_O_2_^+^, C_2_H_3_O^+^, C_2_H_7_O_3_^+^, and CH_4_NO^+^) were not detected as chromatographic peaks in the GC analysis, likely because of their low ambient concentrations and/or adsorption losses within the GC column. However, their chemical identities can be reasonably inferred from previous PTR-MS and forest atmosphere studies. Specifically, C_15_H_25_^+^ and C_2_H_5_O_2_^+^ most likely corresponded to protonated sesquiterpenes and acetic acid, respectively, both of which are frequently observed in atmospheric forest measurements (Hakola *et al*., 2003; Hellén *et al*., 2018; Kesselmeier and Staudt, 1999). C_2_H_3_O^+^ is a fragment of several oxygenated VOCs, including acetic acid (Koss *et al*., 2018), and C_2_H_7_O_3_^+^ may represent a hydrated cluster of protonated acetic acid. Consistent with this interpretation, the time courses of C_2_H_3_O^+^ and C_2_H_7_O_3_^+^ in both the forest-interior and canopy-top air masses strongly correlated with those of C_2_H_5_O_2_^+^ (Supplementary Fig. S5), indicating that these ions likely originated from the fragmentation and hydration of protonated acetic acid, respectively, thereby supporting the assignment of C_2_H_5_O_2_^+^ as acetic acid.

The CH_4_NO^+^ ion likely corresponded to protonated formamide, a compound commonly associated with biomass burning emissions (Koss *et al*., 2018; Stockwell *et al*., 2015). During the measurement period, controlled grassland burning was conducted in agricultural fields and orchards near the forest site, suggesting that the VOCs emitted from these fires were intermittently transported into the forest canopy.

In summary, the daytime factor was dominated by isoprene and its oxidation product, MVK, whereas the nighttime factor was enriched in monoterpenes and sesquiterpenes. Ion intensities (cps) were converted into atmospheric concentrations (ppbv) for eight compounds identified or inferred in this study (isoprene, MVK, acetone, acetaldehyde, α-pinene, acetic acid, sesquiterpenes, and formamide) (Supplementary Fig. S6). The isoprene concentrations showed an apparent difference between the FI and CT atmospheres. The mean daytime isoprene concentrations were approximately 4 ppbv in the FI and 8–10 ppbv in the CT, with CT values ranging from 5 to 30 ppbv. In contrast, daytime concentrations of other VOCs were generally comparable between FI and CT, with MVK at approximately 0.10–0.15 ppbv, acetone at 3–6 ppbv, acetaldehyde at 1–2 ppbv, and acetic acid at approximately 1–2 ppbv. For compounds with higher nighttime concentrations, α-pinene ranged from 0.6 to 0.8 ppbv and sesquiterpenes from 0.04 to 0.12 ppbv. Formamide exhibited relatively minimal diurnal variation and remained at approximately 0.1 ppbv. Therefore, MVK concentrations may indicate contributions from forest emissions, as well as artifact formation within the instrument.

### 3.4. VOC ions characterizing daytime and nighttime modes

The contributions of individual ions to the daytime and nighttime factors varied among the chemical species. To quantify these contributions, the normalized fractions for daytime and nighttime (*NF*_daytime_ and *NF*_nighttime_) were calculated as follows:

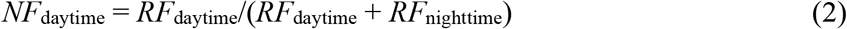

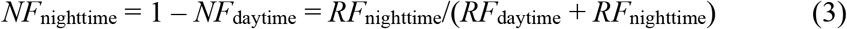

where *RF*_daytime_ and *RF*_nighttime_ represent the relative fractions of individual VOC ions in the mass spectral profiles of daytime and nighttime factors, respectively (Fig. 4). This analysis was performed separately for the FI and CT datasets.

We focused on ions that contributed strongly to either the daytime or nighttime factor, defined as ions with *NF*_daytime_ or *NF*_nighttime_ ≥ 75 %. The number of ions with *NF*_daytime_ ≥ 75 % was 43 and 35 for the FI and CT datasets, respectively. Table 1a lists the selected ions corresponding to the 12 most abundant species, based on their total fractions (*RF*_daytime_ and *RF*_nighttime_) for each dataset. Of these, nine ions were common to both datasets, indicating a high degree of similarity in daytime factors between the FI and CT atmospheres. These common ions included isoprene (C_5_H_9_^+^) and MVK (C_4_H_7_O^+^) as well as other known isoprene oxidation products such as C_5_H_9_O_2_^+^, C_5_H_9_O_3_^+^, C_4_H_7_O_2_^+^, and C_4_H_9_O_2_^+^ (Li *et al*., 2020). In addition, C_3_OH^+^ strongly correlated with isoprene in the canopy-top atmosphere (Supplementary Fig. S7). No corresponding chromatographic peak was observed for this ion, and its chemical identity remains unresolved. However, the lack of a GC peak at the retention time of isoprene indicates that C_3_OH^+^ is unlikely to be an artifact formed from protonated isoprene, suggesting the presence of one or more neutral precursor species with an elemental composition of C_3_O in the forest atmosphere.

**Table 1.** Summary of the twelve most abundant ion species with (a) *NF*_daytime_ ≥ 75 % and (b) *NF*_nighttime_ ≥ 75 % observed in the forest-interior and canopy-top atmospheres. Relative fractions (*RF*) of the daytime and nighttime factors, total fractions, and *NF*_daytime_ are also shown.

|  | Forest-interior (FI) |  |  |  |  | Canopy-top (CT) |  |  |  |  |
| --- | --- | --- | --- | --- | --- | --- | --- | --- | --- | --- |
| | Ion formula<sup>a</sup> | $RF_{\text{daytime}}$ | $RF_{\text{nighttime}}$ | Total fraction<sup>b</sup> | $NF_{\text{daytime}} (\%)$ | Ion formula<sup>a</sup> | $RF_{\text{daytime}}$ | $RF_{\text{nighttime}}$ | Total fraction<sup>b</sup> | $NF_{\text{daytime}} (\%)$ |
| (a) $RF_{\text{daytime}} \geq 75\%$ | **C5H9+** | 0.1343 | 0.0158 | 0.1501 | 89.5 | **C5H9+** | 0.2499 | 0.0113 | 0.2612 | 95.7 |
|  | **C4H7O+** | 0.0745 | 0.0000 | 0.0745 | 100.0 | **C4H7O+** | 0.0664 | 0.0028 | 0.0692 | 96.0 |
|  | **C4H9O2+** | 0.0215 | 0.0026 | 0.0241 | 89.2 | **C3H7O2+** | 0.0175 | 0.0046 | 0.0221 | 79.1 |
|  | **C3H7O2+** | 0.0190 | 0.0032 | 0.0222 | 85.5 | **C4H9O2+** | 0.0168 | 0.0033 | 0.0201 | 83.7 |
|  | **C5H9O2+** | 0.0116 | 0.0027 | 0.0143 | 81.3 | C5H7+ | 0.0151 | 0.0031 | 0.0182 | 82.9 |
|  | C4H11O2+ | 0.0081 | 0.0027 | 0.0108 | 75.3 | **C5H9O2+** | 0.0099 | 0.0029 | 0.0128 | 77.3 |
|  | **C4H7O2+** | 0.0079 | 0.0010 | 0.0089 | 88.7 | C3H5+ | 0.0095 | 0.0018 | 0.0114 | 84.1 |
|  | **C2H7O+** | 0.0075 | 0.0002 | 0.0077 | 97.4 | C3HO+ | 0.0084 | 0.0028 | 0.0111 | 75.2 |
|  | CH7O2+ | 0.0051 | 0.0017 | 0.0069 | 74.8 | **C4H7O2+** | 0.0066 | 0.0018 | 0.0084 | 78.7 |
|  | C5H7O2+ | 0.0051 | 0.0014 | 0.0065 | 77.8 | **C2H7O+** | 0.0061 | 0.0009 | 0.0070 | 87.1 |
|  | **C2H9O2+** | 0.0033 | 0.0000 | 0.0033 | 100.0 | **C2H9O2+** | 0.0030 | 0.0000 | 0.0030 | 100.0 |
|  | **C5H9O3+** | 0.0030 | 0.0000 | 0.0030 | 100.0 | **C5H9O3+** | 0.0027 | 0.0000 | 0.0027 | 99.5 |
| (b) $RF_{\text{nighttime}} \geq 75\%$ | **C6H9+** | 0.0154 | 0.0925 | 0.1080 | 14.3 | C4H9O4+ | 0.0321 | 0.1067 | 0.1387 | 23.1 |
|  | **C10H17+** | 0.0092 | 0.0870 | 0.0962 | 9.6 | **C6H9+** | 0.0114 | 0.0731 | 0.0845 | 13.5 |
|  | **C15H25+** | 0.0011 | 0.0151 | 0.0162 | 6.8 | **C10H17+** | 0.0066 | 0.0679 | 0.0744 | 8.8 |
|  | **CH4NO+** | 0.0038 | 0.0118 | 0.0156 | 24.4 | **CH4NO+** | 0.0039 | 0.0128 | 0.0167 | 23.5 |
|  | **C6H7+** | 0.0034 | 0.0108 | 0.0142 | 23.8 | **C15H25+** | 0.0007 | 0.0111 | 0.0118 | 5.5 |
|  | **C10H17O+** | 0.0016 | 0.0116 | 0.0132 | 12.1 | C7H7+ | 0.0027 | 0.0081 | 0.0108 | 24.8 |
|  | **C10H15+** | 0.0018 | 0.0094 | 0.0112 | 16.1 | **C10H17O+** | 0.0010 | 0.0092 | 0.0103 | 10.2 |
|  | **C6H7O3+** | 0.0014 | 0.0048 | 0.0062 | 22.1 | **C6H7+** | 0.0018 | 0.0081 | 0.0099 | 18.3 |
|  | **H6NO2+** | 0.0012 | 0.0043 | 0.0055 | 22.2 | **C10H15+** | 0.0011 | 0.0076 | 0.0087 | 12.9 |
|  | **C10H19O2+** | 0.0000 | 0.0046 | 0.0046 | 0.1 | **H6NO2+** | 0.0014 | 0.0045 | 0.0059 | 23.0 |
|  | C9H11+ | 0.0000 | 0.0032 | 0.0032 | 0.0 | **C6H7O3+** | 0.0011 | 0.0043 | 0.0054 | 20.6 |
|  | C10H11+ | 0.0001 | 0.0027 | 0.0028 | 2.9 | **C10H19O2+** | 0.0000 | 0.0037 | 0.0037 | 0.1 |

a: Ion formulae shown in bold are common to forest-interior and canopy-top datasets.
b: Total fraction = $RF_{\text{daytime}} + RF_{\text{nighttime}}$

In contrast, the number of ions with *NF*_nighttime_ ≥ 75 % was 254 and 314 for the FI and CT datasets, respectively. Similar to the daytime, the 12 most abundant species based on their total fractions are listed in Table 1b, of which 10 ions were common to both datasets. Notably, terpene-related ions other than monoterpenes (C_10_H_17_^+^) and sesquiterpenes (C_15_H_25_^+^), including C H ^+^, C_10_H_17_O^+^, and C_10_H_19_O_2_^+^, were also identified, indicating that nighttime factors include a broader suite of terpene-related compounds and their oxidation products.

### 3.5. Light and temperature controls on the daytime mode

The enhancement of daytime and nighttime factors in the forest-interior and canopy-top atmospheres was compared with the temperature and light intensity (photosynthetically active radiation, PAR) measured in close proximity to the individual VOC sampling ports.

The daytime factor was dominated by isoprene (Fig. 4), whose emissions are strongly controlled by PAR and temperature (Guenther *et al*., 1993). Therefore, we tested whether enhancement of the daytime factor was associated with variations in temperature and light intensity.

The relative enhancement of the daytime factor (*RE*_daytime_) is defined as the time-course-based fractional contribution of the daytime factor to the sum of the daytime and nighttime factors, generally expressed as follows:

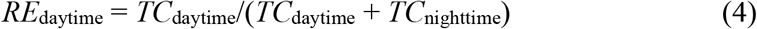

This formulation was applied separately to the FI and CT datasets as *RE*_daytime-FI_ and *RE*_daytime-CT_, where *TC*_daytime-FI_, *TC*_nighttime-FI_, *TC*_daytime-CT_, and *TC*_nighttime-CT_ represent PMF-derived time courses for the respective daytime and nighttime factors.

Model G93 (Guenther *et al*., 1993) is a well-established model used to estimate isoprene emission rates as functions of light and temperature.

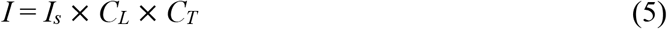

where *I* is the isoprene emission rate at temperature *T* (K) and PAR flux *L* (μmol m^-2^ s^-1^), *I_s_* is the isoprene emission rate at a standard temperature, *T_S_* (K), and standardized PAR flux. *C*_L_ and *C*_T_ are the light-and temperature-dependent factors, respectively. *C_L_* is defined as follows:

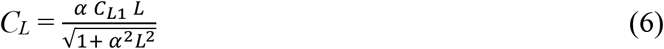

where α (= 0.0027) and *C_L_*_1_ (= 1.066) are empirical coefficients determined from the emission rate measurements of selected plant species, such as eucalyptus, sweet gum, aspen, and velvet bean. This formulation simulates a near-linear increase in isoprene emissions up to a saturation point and resembles the equations used to describe light dependence in photosynthesis (Smith, 1937; Harley and Tenhunen, 1991). *C_T_* is expressed as follows:

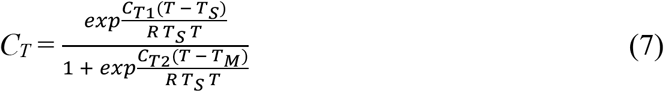

where *R* is a constant (= 8.314 J K^-1^ mol^-1^), and *C_T_*_1_ (= 95,000 J mol^-1^), *C_T_*_2_ (= 230,000 J mol^-1^), and *T_M_* (= 314 K) are empirical coefficients derived from the same datasets used to determine α and *C_L_*_1_. The standard temperature was *T*_S_ = 303 K (30 °C), as defined in G93. This equation simulates the temperature response of the enzymatic activity (Johnson *et al*., 1942; Sharpe and DeMichele, 1977). Because *I_s_* is constant, we compared *RE*_daytime_ with the product *C_L_* × *C_T_*.

Fig. 5 shows the relationship between *RE*_daytime_ and *C_L_* × *C_T_* for a single day (0:00–24:00 on August 30, 2024). As shown in Fig. 5A, *RE*_daytime_ and *C_L_* × *C_T_* are plotted as functions of time, along with temperature (*T*) and PAR (*L*). Both *RE*_daytime_ and *C_L_* × *C_T_* increased as PAR began to increase after 05:00. The values of *C_L_* × *C_T_* reached their maximum between 12:30 and 14:30, whereas *RE*_daytime_ remained elevated over a broader interval from 12:00 to 16:30. Notably, even after *C_L_* × *C_T_* declined to approximately zero after 18:00 (sunset), *RE*_daytime_ persisted at appreciable levels. Fig. 5B shows a direct comparison between *RE*_daytime_ and *C_L_* × *C_T_*. Notably, the *RE*_daytime_ during the afternoon and evening was substantially higher than that during the morning. *RE*_daytime_ varied even under similar values of *C_L_* × *C_T_* depending on the time of day (*RE*_daytime-FI_ = 0.46 at *C_L_* × *C_T_* = 0.05 [08:37]; *RE*_daytime-FI_ = 0.79 at *C_L_* × *C_T_* = 0.06 [16:49]; *RE*_daytime-CT_ = 0.64 at *C_L_* × *C_T_* = 0.43 [08:57]; *RE*_daytime-CT_ = 0.88 at *C_L_* × *C_T_* = 0.41 [15:09]). A comparable behavior was also observed on another day (09:00 on September 1, 2024, to 09:00 on September 2, 2024; Supplementary Fig. S8).

**Fig. 5.**
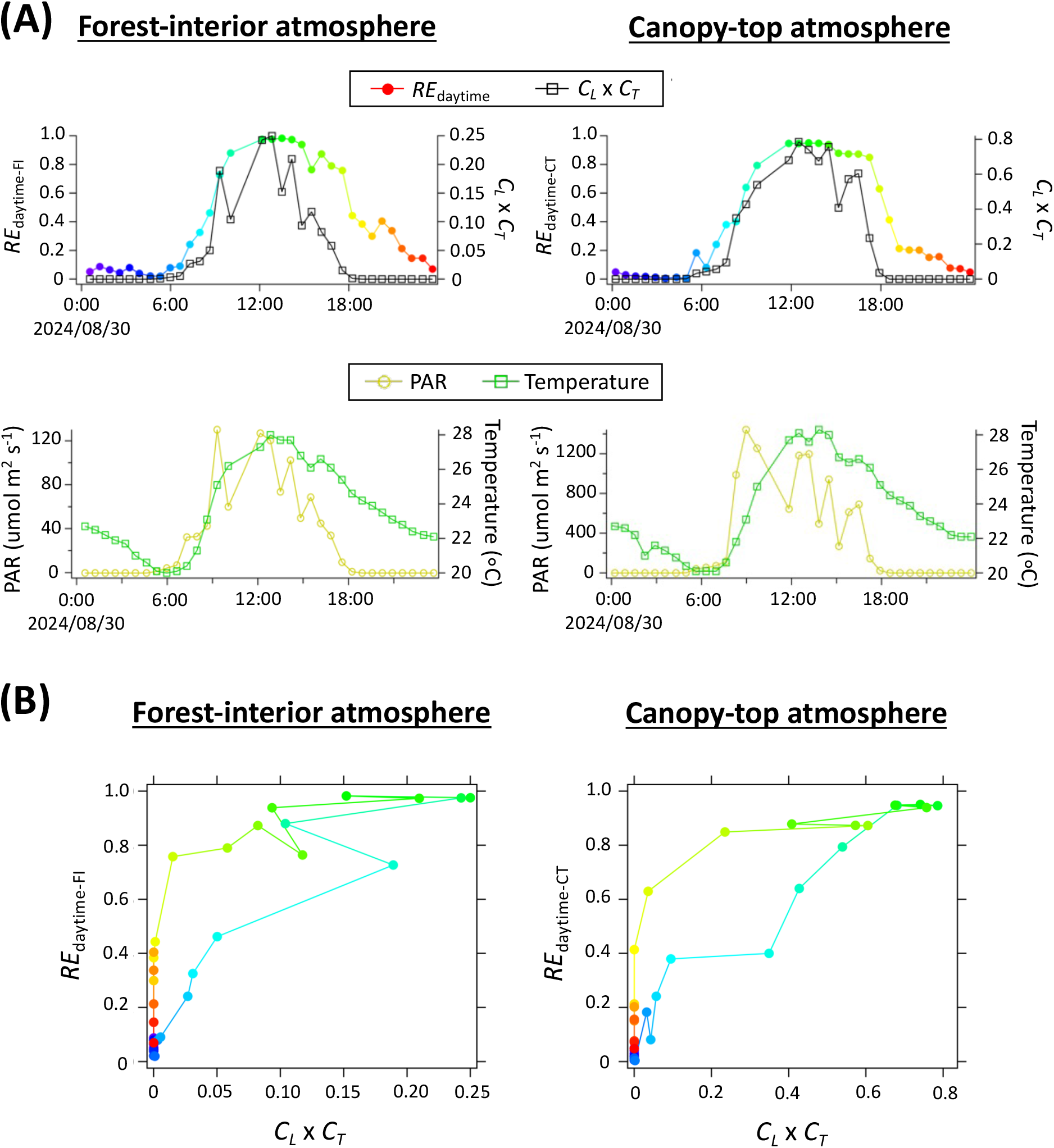
Relationship between *RE*_daytime-FI_, *RE*_daytime-CT_, and *C_L_* × *C_T_* for a single day (0:00– 24:00 on August 30, 2024). (A) Time courses of *RE*_daytime-FI_ and *RE*_daytime-CT_, *C_L_* × *C_T_*, PAR, and temperature. (B) Scatter plots of *RE*_daytime-FI_ or *RE*_daytime-CT_ versus *C_L_* × *C_T_*. Color codes in (B) indicate time of day and correspond to those used for *RE*_daytime_ in (A).

Collectively, *RE*_daytime_ exhibited a pronounced morning–afternoon asymmetry; for comparable values of *C_L_* × *C_T_*, afternoon (and early evening) values were systematically higher than morning values. This behavior suggests the presence of additional controls beyond the simple light and temperature dependences captured by G93. Possible explanations include leaf-level physiological hysteresis, canopy-scale transport processes, and atmospheric chemical processing. However, because we did not measure the gene expression of isoprene, monoterpenes, and sesquiterpene synthases, boundary-layer development, or atmospheric oxidants, such as OH, O_3_ and NO*_x_*, this interpretation should be regarded as hypothesis-forming.

### 3.6. Interannual reproducibility of daytime and nighttime VOC modes

To evaluate the interannual reproducibility of the PMF-derived VOC patterns, we applied the same analysis framework to an independent dataset acquired during the summer of 2025 under rain-free conditions (September 4–6, 2025), which was comparable to that of 2024. The PTR-ToF-MS measurements and data selection criteria were identical to those described above.

The PMF analysis of the 2025 dataset resulted in a distinct separation into two dominant factors characterized by daytime and nighttime enhancement, consistent with the results obtained for the 2024 observations (Supplementary Fig. S9A and S10A). As in 2024, the two-factor solution captured most of the variability in total VOC ion intensities, with an overall residual of approximately 20 %.

The mass spectral profiles of the daytime and nighttime factors in 2025 closely resembled those obtained from the 2024 dataset (Supplementary Fig. S9B, S10B, and Supplementary Table S1). The daytime factor was dominated by isoprene (C_5_H_9_^+^) and its oxidation products, including MVK (C_4_H_7_O^+^) and other products (C_5_H_9_O_2_^+^, C_5_H_9_O_3_^+^, C_4_H_7_O_2_^+^, and C_4_H_9_O_2_^+^), as well as C_3_OH^+^, whereas the nighttime factor was enriched in monoterpenes (C_10_H_17_^+^), sesquiterpenes (C H ^+^), and related terpene-derived ions (C_10_H_17_O^+^). Correlation coefficients (*r*^2^) between the mass spectral profiles obtained in 2024 and 2025 ranged from 0.65 to 0.80 for the daytime and nighttime factors across the forest-interior and canopy-top atmospheres, based on relative ion fractions evaluated on a logarithmic scale (Supplementary Fig. S9C and S10C). Strong correlations between isoprene and C_3_OH^+^ were consistently observed in the 2025 dataset (Supplementary Fig. S7B).

A relationship between the relative enhancement of the daytime factor (*RE*_daytime_) and the combined light and temperature response terms (*C_L_* × *C_T_*) was consistently observed in both years (Supplementary Fig. S11). As observed in 2024, *RE*_daytime_ increased with increasing *C_L_* × *C_T_* during the morning; however, it remained elevated in the afternoon and early evening, resulting in pronounced morning–afternoon asymmetry under comparable environmental conditions.

These results indicate that the separation of daytime and nighttime VOC factors, their associated chemical characteristics, and their relationship with light and temperature are recurring features of forest-scale VOC variability in this ecosystem across the years.

## 4. Discussion

In this study, we applied positive matrix factorization (PMF) to forest-scale VOC measurements in a *Quercus crispula* forest to identify the recurring and dominant modes of VOC emissions and to explore their environmental, physiological, and micrometeorological drivers. Hereafter, the PMF-derived factors are discussed as the recurring modes of VOC emissions associated with distinct environmental and physiological conditions. In the following sections, we discuss these interpretations, assess their robustness across years and sampling locations, and discuss the methodological and biological implications of our findings.

### 4.1. Physiological interpretation of recurring daytime and nighttime VOC modes

Using PTR-ToF-MS combined with PMF, we identified two recurring modes of forest VOC variability in a *Q. crispula* forest. These patterns were consistent with well-established plant metabolic controls: a daytime, light-, and temperature-dependent isoprene/MVK-dominated mode and a nighttime mode characterized by enhanced contributions of monoterpenes and sesquiterpenes. The daytime mode is consistent with chloroplastic MEP (non-mevalonate) pathway regulation and isoprene synthase (*IspS*) activity (Sharkey and Monson, 2017), including the circadian regulation of *IspS* transcripts and enzyme activity reported for poplar (Loivamäki *et al*., 2007) and oil palm (Wilkinson *et al*., 2006). This behavior is also consistent with the biochemical framework underlying the G93 algorithm, which describes isoprene emission as a function of light and temperature response (*C_L_* × *C_T_*) (Guenther *et al*., 1993; Sharkey and Loreto, 1996; Karl *et al*., 2009). In contrast, the nighttime mode aligns with storage-and diffusion-driven terpenoid emissions under conditions of reduced photochemical oxidation, as previously reported for monoterpenes and sesquiterpenes (Schade and Goldstein, 2002; Hakola, 2003; Holzke, 2006).

### 4.2. Generality and reproducibility of forest VOC variability

To evaluate the robustness of the daytime and nighttime VOC modes, we applied the same PMF analysis to an independent dataset from the summer of 2025 and assessed the reproducibility of (i) an apparent separation into daytime and nighttime factors, (ii) the factor mass spectra, particularly the dominance of isoprene/MVK during daytime and monoterpenes/sesquiterpenes at night, and (iii) the relationship between daytime factor enhancement (*RE*_daytime_) and the light-and temperature-response term (*C_L_* × *C_T_*), including the observed morning–afternoon asymmetry. The high degree of reproducibility across the years indicates that these factors represent recurring modes of forest-scale VOC variability rather than site-or year-specific anomalies.

Previous studies of European forests further support the validity of this PMF-derived interpretation. The extracted compositional fingerprints, daytime enhancement of isoprene and MVK and nighttime dominance of monoterpenes and sesquiterpenes, are consistent with observations from boreal forest ecosystems, including coniferous and mixed forests in France and Finland (Li *et al*., 2020, 2021). Dominant emitted terpenoids differ among forest types, being isoprene-dominated in the present temperate broad-leaved forest and monoterpene-dominated in boreal forests. The emergence of comparable daytime and nighttime emission modes suggests that these patterns are consistent with general physiological and micrometeorological controls, rather than depending on differences in the dominant emitted chemical species. Oxygenated VOCs, such as acetone and acetaldehyde, contribute to both factors, consistent with their mixed sources and sinks, including direct biogenic emissions, secondary atmospheric formation, deposition, and forest-scale mixing and entrainment processes (Karl *et al*., 2009). These results suggest that daytime and nighttime VOC modes represent common features of forest VOC variability across contrasting forest ecosystems.

### 4.3. A morning–afternoon asymmetry in relation to environmental drivers: complex physiological and forest-scale mechanisms

The daytime factor covaried with isoprene and exhibited a strong dependence on photosynthetically active radiation and temperature when evaluated against the combined light and temperature response terms (*C_L_* × *C_T_*; Guenther *et al*., 1993). However, a systematic morning– afternoon asymmetry was observed; for comparable values of *C_L_* × *C_T_*, the afternoon and early evening values of the daytime factor enhancement (*RE*_daytime_) consistently exceeded those observed in the morning. This behavior may be explained by two non-exclusive mechanisms involving plant physiology and forest-scale atmospheric processes.

1. **Leaf-level metabolic asymmetry:** Afternoon enhancement of daytime factors may result from the diel dynamics of substrate pools in the MEP pathway and the regulation of *IspS*, including circadian modulation and temperature-dependent enzyme kinetics that increase emission efficiency later in the day, even under similar instantaneous drivers (Brüggemann and Schnitzler, 2002; Wilkinson *et al*., 2006; Loivamäki *et al*., 2007; Wiberley *et al*., 2009). Transgenic poplar studies have further indicated that substrate availability (DMAPP), *IspS* localization, and kinetics exert primary short-term control over isoprene emissions, with circadian regulation of *IspS* contributing to diel asymmetry (Loivamäki *et al*., 2007).
2. **Forest-scale atmospheric transport and oxidative-loss asymmetry:** At the canopy scale, morning boundary layer growth and entrainment promote the dilution of emitted VOCs, whereas reduced oxidative loss and enhanced in-canopy accumulation later in the day can sustain elevated daytime-factor contributions, even as *C_L_* × *C_T_* declines toward sunset (Stroud *et al*., 2005; Wolfe *et al*., 2011; Sarkar *et al*., 2020). Large-eddy simulation studies that explicitly resolve turbulent eddies within and above forest canopies, together with field observations, have demonstrated that coupled emissions, transport, and chemistry can generate these phase lags (Clifton *et al*., 2022).

Although the present dataset precludes quantitatively partitioning the relative contributions of these mechanisms, the close correspondence between the PMF-derived factor composition and established plant emission behavior supports the interpretation that physiological processes are important in influencing forest-scale VOC variability in *Q. crispula* forests. Although we did not directly measure *IspS* gene expression, enzyme activity, leaf-level VOC emission rates, boundary-layer development, or atmospheric oxidants (OH, O_3_, and NO*_x_*), the mechanisms discussed above can be considered as integrative interpretations consistent with our observations and previous studies rather than direct causal attribution.

### 4.4. Implications of unassigned VOC ions for daytime and nighttime modes

In addition to the well-characterized VOC species, our PMF analysis revealed ions that could not be unambiguously assigned to known compounds but consistently contributed to the identified daytime and nighttime factors. These unassigned signals indicate that forest-scale VOC variability is not completely captured by the currently identified biogenic VOCs and their well-established oxidation products.

For example, the ion C_3_OH⁺ exhibited a strong temporal correlation with isoprene in the canopy-top atmosphere but lacked a corresponding chromatographic peak in the GC/PTR-ToF-MS analysis. The absence of a peak at the retention time of isoprene suggests that C_3_OH⁺ is unlikely to originate from instrumental fragmentation of protonated isoprene; rather, it indicates the presence of a neutral precursor with the elemental composition C_3_O in the forest atmosphere. The close coupling between C₃OH⁺ and isoprene suggests that this species may be associated with processes occurring under active photosynthetic conditions, either through plant metabolism or rapid in-canopy chemical transformation.

These observations suggest that although the dominant modes of forest-scale VOC variability can be predominantly interpreted regarding established physiological and micrometeorological controls, additional unassigned chemical components also contribute to the variability captured by the daytime and nighttime VOC modes. This highlights the value of non-targeted analytical frameworks for capturing these signals and motivates further efforts to constrain their chemical identity and origin using complementary approaches.

### 4.5. Methodological advantages and limitations of the PTR-ToF-MS plus PMF framework

Pairing high-resolution PTR-ToF-MS with positive matrix factorization (PMF) enables a non-targeted, process-oriented characterization of forest VOC variability by extracting co-varying ensembles of ions that represent dominant modes of VOC variability, such as daytime isoprene-rich and nighttime terpene-rich patterns. When used alone, PTR-ToF-MS measurements primarily provide mass-to-charge (*m/z*) ratios and corresponding elemental compositions of detected ions, indicating that compound-resolved identification frequently becomes a prerequisite for mechanistic interpretation. In contrast, coupling PTR-ToF-MS with PMF enables dominant modes of variability to be identified even when individual compounds cannot be confidently assigned. As demonstrated in this study, this framework remains effective when the chromatographic separation is incomplete or when unassigned ions contribute substantially to the observed variability. Therefore, the PMF-based analysis provides a top-down perspective, whereby dominant modes of variability are initially identified from covariance structures in the data, and chemically or physiologically informative ions associated with these processes can subsequently be prioritized for targeted identification. This approach complements conventional bottom-up compound-resolved analyses and provides a more integrated perspective on complex forest-scale VOC dynamics.

However, this study had several limitations. Because PMF does not require prior specifications of tracers, the attribution of extracted factors inherently requires interpretation, and factor time courses integrate the combined effects of emissions, atmospheric transport, and chemical processing. Therefore, a robust interpretation depends on independent constraints, as highlighted by the morning–afternoon asymmetry observed in this study, including environmental drivers (light and temperature), plant physiological information (enzyme expression or activity), and atmospheric context (oxidant levels, turbulence, and boundary-layer development). In addition, low-abundance ions tend to be underrepresented in PMF solutions, which potentially limits the identification of processes associated with chemically minor but mechanistically informative species.

### 4.6. Broader implications and perspectives

Plant VOC emissions have been extensively studied from both the physiological and ecological perspectives. Specifically, isoprene has attracted particular attention because of its global abundance and diverse roles in plant responses to environmental stress (Guenther *et al*., 1995; Loreto *et al*., 2001; Sasaki *et al*., 2007; Vickers *et al*., 2009; Tattini *et al*., 2014; Zuo *et al*., 2025). In addition to its direct physiological functions, isoprene emissions are closely linked to broader metabolic regulation, including interactions with isoprenoid and phenylpropanoid metabolism and plant hormone signaling pathways (Monson *et al*., 2021). These observations suggest that VOC emissions are closely associated with regulated physiological processes rather than resulting solely from passive byproducts of metabolism.

The present study extends this perspective from the leaf to the forest scale. The recurring daytime and nighttime VOC modes were reproducible across years and were associated with distinct chemical compositions and environmental responses. The daytime mode was primarily characterized by isoprene and its oxidation products, whereas the nighttime mode was enriched in monoterpenes and sesquiterpenes. Comparable daytime and nighttime VOC modes have also been reported in contrasting forest ecosystems, despite substantial differences in the dominant emitted compounds (Li *et al*., 2020, 2021). Together, these observations suggest that common physiological and environmental constraints may shape forest-scale VOC variability across a wide range of forest types.

From a broader perspective, integrating non-targeted, high-time-resolution VOC measurements with process-oriented analyses provides a pathway for linking ecological interactions within forests with atmospheric chemistry and climate processes (Satake *et al*., 2024). By providing a forest-scale perspective on recurring VOC variability, the PTR-ToF-MS plus PMF framework facilitates linking leaf-level physiological processes and ecosystem-scale atmospheric exchanges. Extending this approach across forest types, seasons, and environmental conditions, while resolving vertical heterogeneity within forest atmospheres, including poorly characterized dynamics near the forest floor, will further advance our understanding of forest VOC variability. Combining these observations with complementary physiological, ecological, and micrometeorological measurements will provide comprehensive insights into how biological activity within forests is expressed by atmospheric-scale VOC dynamics.

## Supplementary data

The following supplementary data are available at online.

**Table S1.** Relative fractions in the mass spectral profiles (daytime FI, nighttime FI, daytime CT, and nighttime CT) for individual ions obtained from the 2024 and 2025 datasets.

**Fig. S1**. Study site and sampling locations.

**Fig. S2.** Schematic representation of PMF factorization.

**Fig. S3.** Comparison of time courses for the forest-interior atmospheres in the 2024 dataset from 2-, 3-, and 4-factor PMF solutions.

**Fig. S4.** GC/PTR-ToF-MS results of canopy-top atmospheres during daytime and nighttime on August 24, 2025.

**Fig. S5.** Correlation among C_2_H_5_O_2_^+^, C_2_H_7_O_3_^+^, and C_2_H_3_O^+^ in the 2024 dataset.

**Fig. S6.** Time courses of concentrations of eight VOCs in the 2024 dataset.

**Fig. S7.** Correlation between C_3_OH^+^ and C_5_H_9_^+^ (isoprene).

**Fig. S8.** Relationship between *RE*_daytime-FI_, *RE*_daytime-CT_, and *C_L_* × *C_T_* from September 1 to 2, 2024.

**Fig. S9.** PMF results for the two-factor solution applied to VOC ion signals measured in forest-interior atmospheres during three rain-free days in the 2025 datasets.

**Fig. S10.** PMF results for the two-factor solution applied to VOC ion signals measured in canopy-top atmospheres during three rain-free days in the 2025 datasets.

**Fig. S11.** Relationship between *RE*_daytime-FI_, *RE*_daytime-CT_, and *C_L_* × *C_T_* from September 4 to 7, 2024.

## Acknowledgements

We thank Ms. Saki Oe, Mr. Haruki Nagata, Mr. Yuta Hamamoto, and CHABUDAI for their support during the field campaigns. We also thank Associate Professor Sachinobu Ishida (Hirosaki University) for providing meteorological data. This study was supported by the JST PRESTO (Grant No. JPMJPR21D7) and JSPS KAKENHI (Grant Nos. JP23H04969 and JP24K01518).

## Author contribution

K.S., Y.S., Y.K., and T.S. designed the experiments. K.S., Y.S., Y.K., D.F., A.K., A.M. and S.T. performed the experiments. H.Y. and K.S. managed the forest site and constructed the rolling tower. K.S., T.S., and K.Y. drafted the manuscript. All authors revised and approved the final manuscript.

## Data availability

The primary data supporting the findings of this study are available from the corresponding author upon request.

## Figure legends and Alt text for accessibility

**Alt text:** VOC measurements from forest-interior and canopy-top air show that a small number of ions account for a large fraction of total VOC signal intensity. Time-course and diurnal-cycle data indicate that individual ions exhibit distinct temporal behaviors, including daytime enhancement, nighttime enhancement, or relatively weak diurnal variation. Total VOC abundance is generally higher during daytime than at night.

**Alt text:** Positive matrix factorization (PMF) applied to forest-interior VOC measurements in 2024 identifies two dominant factors. One factor is enhanced during daytime, whereas the other is enhanced during nighttime. The two factors have distinct mass-spectral profiles, indicating differences in the VOC ions contributing to daytime and nighttime variability.

**Alt text:** PMF analysis of canopy-top VOC measurements in 2024 also identifies two dominant factors with contrasting daytime and nighttime behavior. The daytime factor contributes most strongly during daylight hours, whereas the nighttime factor dominates after sunset. Their mass-spectral profiles differ, reflecting different chemical compositions associated with daytime and nighttime atmospheric conditions.

**Alt text:** Chemical compositions of PMF-derived daytime and nighttime factors are compared for forest-interior and canopy-top atmospheres. Daytime factors are characterized by larger contributions from isoprene-related ions and their oxidation products, whereas nighttime factors contain greater contributions from monoterpene-, sesquiterpene-, and other terpene-related ions. Similar patterns are observed in both sampling locations.

**Alt text:** The contribution of the daytime VOC factor increases with increasing light and temperature conditions represented by the product *C_L_* × *C_T_*. However, the relationship differs between morning and afternoon periods, with daytime-factor enhancement remaining higher in the afternoon and early evening than would be expected from light and temperature alone. This pattern indicates a pronounced morning–afternoon asymmetry in forest-scale VOC variability.

## Abbreviations

Cps: Counts-per-second
CT: Canopy top
FI: Forest interior
MS: Mass spectra
MVK: Methyl vinyl ketone
NF: Normalized fraction
PAR: Photosynthetically active radiation
PMF: Positive matrix factorization
PTR-ToF-MS: Proton-transfer-reaction time-of-flight mass spectrometry
RE: Relative enhancement
RF: Relative fractions
TC: time course
VOC: Volatile organic compounds

## References

Atkinson R. 1990. Gas-phase tropospheric chemistry of organic compounds: A review. Atmospheric Environment 24A, 1–41.

Brüggemann N, Schnitzler J-P. 2002. Comparison of isoprene emission, intercellular isoprene concentration and photosynthetic performance in water-limited oak (Quercus pubescens Willd. and Quercus robur L.) saplings. Plant Biology 4, 456–463.

Clifton OE, Patton EG, Wang S, Barth M, Orlando J, Schwantes RH. 2022. Large-eddy simulation for investigating coupled forest canopy and turbulence influences on atmospheric chemistry. Journal of Advances in Modeling Earth Systems 14, e2022MS003078.

Crounse JD, Nielsen LB, Jørgensen S, Kjaergaard HG, Wennberg PO. 2013. Autoxidation of organic compounds in the atmosphere. Journal of Physical Chemistry Letters 4, 3513–3520.

Dumont C, Verreyken BW, Schoon N, Bergmans B, Heinesch B, Amelynck C. 2026. Multi-year observations of BVOCs and ozone: concentrations and fluxes measured above and below the canopy in a mixed temperate forest. Earth System Science Data 18, 617–654.

Ehn M, Thornton JA, Kleist E, et al. 2014. A large source of low-volatility secondary organic aerosol. Nature 506, 476–479.

Guenther AB, Zimmerman PR, Harley PC, Monson RK, Fall R. 1993. Isoprene and monoterpene emission variability: model evaluations and sensitivity analyses. Journal of Geophysical Research 98, 12609–12617.

Guenther A, Hewitt CN, Erickson D, et al. 1995. A global model of natural volatile organic compound emissions. Journal of Geophysical Research 100, 8873–8892.

Guenther AB, Jiang X, Heald CL, Sakulyanontvittaya T, Duhl TA, Emmons LK, Wang X. 2012. The Model of Emissions of Gases and Aerosols from Nature (MEGAN2.1): an extended and updated framework for modeling biogenic emissions. Geoscientific Model Development 5, 1471–1492.

Hakola H, Tarvainen V, Laurila T, Hiltunen V, Hellén H, Keronen P. 2003. Seasonal variation of VOC concentrations above a boreal coniferous forest. Atmospheric Environment 37, 1623–1634.

Harley PC, Tenhunen JD. 1991. Modeling the photosynthetic response of C₃ leaves to environmental factors In: Boote KJ, Loomis RS, eds. Modeling Crop Photosynthesis — from Biochemistry to Canopy. CSSA Special Publications. ISSN: 2165-9745.

Hellén H, Praplan AP, Tykkä T, Ylivinkka I, Vakkari V, Bäck J, Petäjä T, Kulmala M, Hakola H. 2018. Long-term measurements of volatile organic compounds highlight the importance of sesquiterpenes for the atmospheric chemistry of a boreal forest. Atmospheric Chemistry and Physics 18, 13839–13863.

Holopainen JK, Gershenzon J. 2010. Multiple stress factors and the emission of plant VOCs. Trends in Plant Science 15, 176–184.

Holzke C, Hoffmann T, Jaeger L, Koppmann R, Zimmer W. 2006. Diurnal and seasonal variation of monoterpene and sesquiterpene emissions from Scots pine (*Pinus sylvestris* L.). Atmospheric Environment 40, 3174–3185.

Ishida S. 2012. General meteorological conditions of the Shirakami Natural Science Park, 2011, Bulletin of the Shirakami Institute for Environmental Sciences, Hirosaki University, SHIRAKAMI-SANCHI 1: 19–27. ISSN 2186-9219

Johnson FH, Eyring H, Williams RW. 1942. The nature of enzyme inhibitions in bacterial luminescence: Sulfanilamide, urethane, temperature and pressure. Journal of Cellular and Comparative Physiology 20, 247–268.

Karl T, Apel E, Hodzic A, Riemer DD, Blake DR, Wiedinmyer C. 2009. Emissions of volatile organic compounds inferred from airborne flux measurements over a megacity. Atmospheric Chemistry and Physics 9, 271–285.

Kavouras IG, Mihalopoulos N, Stephanou EG. 1998. Formation of atmospheric particles from organic acids produced by forests. Nature 395, 683–686.

Kesselmeier J, Staudt M. 1999. Biogenic volatile organic compounds (VOC): an overview on emission, physiology and ecology. Journal of Atmospheric Chemistry 33, 23–88.

Koss AR, Sekimoto K, Gilman JB, et al. 2018. Non-methane organic gas emissions from biomass burning: identification, quantification, and emission factors from PTR-ToF during the FIREX 2016 laboratory experiment. Atmospheric Chemistry and Physics 18, 3299–3319. 10.5194/acp-18-3299-2018.

Krechmer JE, Lopez-Hilfiker F, Koss A, et al. 2018. Evaluation of a new reagent-ion source and focusing ion-molecule reactor for use in proton-transfer-reaction mass spectrometry. Analytical Chemistry 90, 12011–12018.

Li HY, Riva M, Rantala P, et al. 2020. Terpenes and their oxidation products in the French Landes forest: insights from Vocus PTR-TOF measurements. Atmospheric Chemistry and Physics 20, 1941–1959.

Li HY, Canagaratna MR, Riva M, et al. 2021. Atmospheric organic vapors in two European pine forests measured by a Vocus PTR-TOF: insights into monoterpene and sesquiterpene oxidation processes. Atmospheric Chemistry and Physics 21, 4123–4147.

Loivamäki M, Louis S, Cinege G, Zimmer I, Fischbach RJ, Schnitzler JP. 2007. Circadian rhythms of isoprene biosynthesis in grey poplar leaves. Plant Physiology 143, 540–551.

Loreto F, Mannozzi M, Maris C, Nascetti P, Ferranti F, Pasqualini S. 2001. Ozone quenching properties of isoprene and its antioxidant role in leaves. Plant Physiology 126, 993–1000.

Monson RK, Weraduwage SM, Rosenkranz M, Schnitzler JP, Sharkey TD. 2021. Leaf isoprene emission as a trait that mediates the growth–defense tradeoff in the face of climate stress. Oecologia 197, 885–902.

Müller M, Mielke LH, Breitenlechner M, McLuckey SA, Shepson PB, Wisthaler A, Hansel A. 2009. MS/MS studies for the selective detection of isomeric biogenic VOCs using a Townsend discharge triple quadrupole tandem MS and a PTR-linear ion trap MS. Atmospheric Measurement Techniques 2, 703–712.

Niederbacher B, Winkler JB, Schnitzler JP. 2015. Volatile organic compounds as non-invasive markers for plant phenotyping. Journal of Experimental Botany 66, 5403–5416.

Ortega J, Helmig D. 2008. Approaches for quantifying reactive and low-volatility biogenic VOCs. Atmospheric Chemistry and Physics 8, 4989–5002.

Paatero P, Tapper U. 1994. Positive matrix factorization: A non-negative factor model with optimal utilization of error estimates. Environmetrics 5, 111–126.

Paatero P. 1997. Least squares formulation of robust non-negative factor analysis. Chemometrics and Intelligent Laboratory Systems 37, 23–35.

Peñuelas J, Llusià J. 2003. BVOCs: Plant defense against climate warming? Trends in Plant Science 8, 105–109.

Peñuelas J, Llusià J. 2004. Plant VOC emissions: Making use of the unavoidable. Trends in Ecology and Evolution 19, 402–404.

Peñuelas J, Asensio D, Tholl D, Wenke K, Rosenkranz M, Piechulla B, Schnitzler J. 2014. Biogenic volatile emissions from the soil. Plant, Cell and Environment 37, 1866–1891.

Petersen AK, Holst T, Mölder M, Kljun N, Rinne J. 2023. Vertical distribution of sources and sinks of volatile organic compounds within a boreal forest canopy. Atmospheric Chemistry and Physics 23, 7839–7856.

Rinne J, Taipale R, Markkanen T, Ruuskanen TM, Hellén H, Kajos MK, Vesala T, Kulmala M. 2016. Isoprene and monoterpene fluxes measured above a boreal forest. Biogeosciences 13, 1913–1930.

Rivera-Rios JC, Nguyen TB, Crounse JD, et al. 2014. Conversion of hydroperoxides to carbonyls in field and laboratory instrumentation: Observational bias in diagnosing pristine versus anthropogenically controlled atmospheric chemistry. Geophysical Research Letters 41, 8645–8651.

Sarkar C, Guenther AB, Park JH, et al. 2020. PTR-ToF-MS eddy covariance measurements of isoprene and monoterpene fluxes from an eastern Amazonian rainforest. Atmospheric Chemistry and Physics 20, 7179–7191.

Sasaki K, Saito T, Lämsä M, Oksman-Caldentey KM, Suzuki M, Ohyama K, Muranaka T, Ohara K, Yazaki K. 2007. Plants utilize isoprene emission as a thermotolerance mechanism. Plant and Cell Physiology 48, 1254–1262.

Satake A, Hagiwara T, Nagano AJ, Yamaguchi N, Sekimoto K, Shiojiri K, Sudo K. 2024. Plant molecular phenology and climate feedbacks mediated by BVOCs. Annual Review of Plant Biology 75, 605–627.

Schade GW, Goldstein AH. 2002. Seasonal measurements of volatile organic compounds in a coniferous forest. Journal of Geophysical Research 107, 8440.

Sekimoto K, Li SM, Yuan B, Koss A, Coggon M, Warneke C, De Gouw J. 2017. PTR-MS sensitivity calculation for organic trace gases. International Journal of Mass Spectrometry 421, 71–94.

Sharkey TD, Loreto F. 1996. Water stress, temperature, and light effects on the capacity for isoprene emission. Plant Physiology 110, 1067–1074.

Sharkey TD, Monson RK. 2017. Isoprene research – 60 years later, the biology is still enigmatic. Plant, Cell and Environment 40, 1671–1678.

Sharpe PJH, DeMichele DW. 1977. Reaction kinetics of poikilotherm development. Journal of Theoretical Biology 64, 649–670.

Smith EL. 1937. The influence of light and carbon dioxide on photosynthesis. Journal of General Physiology 20, 807–830.

Stockwell CE, Veres PR, Williams J, Yokelson RJ. 2015. Characterization of biomass burning emissions from cooking fires, peat, crop residue, and other fuels with high-resolution proton-transfer-reaction time-of-flight mass spectrometry. Atmospheric Chemistry and Physics 15, 845–865.

Stoy PC, Trowbridge AM, Siqueira MB, Freire LS, Phillips RP, Jacobs L, Wiesner S, Monson RK, Novick KA. 2021. Vapor pressure deficit helps explain biogenic volatile organic compound fluxes from the forest floor and canopy of a temperate deciduous forest. Oecologia 195, 165–178.

Stroud C, Makar P, Karl T, Guenther A, Geron C, Turnipseed A, Nemitz E, Baker B, Potosnak M, Fuentes JD. 2005. Role of canopy-scale photochemistry in modifying biogenic-atmosphere exchange of reactive terpene species: Results from the CELTIC field study. Journal of Geophysical Research 110, D17303.

Tani A, Kawawata Y. 2008. Isoprene emission from the major native Quercus spp. in Japan. Atmospheric Environment 42, 4540–4550.

Tattini M, Velikova V, Vickers C, Brunetti C, Di Ferdinando M, Trivellini A, Fineschi S, Agati G, Ferrini F, Loreto F. 2014. Isoprene production in transgenic tobacco alters isoprenoid, non-structural carbohydrate and phenylpropanoid metabolism, and protects photosynthesis from drought stress. Plant, Cell and Environment 37, 1950–1964.

Tholl D, Boland W, Hansel A, Loreto F, Röse US, Schnitzler JP. 2006. Practical approaches to plant volatile analysis. The Plant Journal 45, 540–560.

Ulbrich IM, Canagaratna MR, Zhang Q, Worsnop DR, Jimenez JL. 2009. Interpretation of organic components from positive matrix factorization of aerosol mass spectrometric data. Atmospheric Chemistry and Physics 9, 2891–2918.

Vickers CE, Possell M, Cojocariu CI, Velikova VB, Laothawornkitkul J, Ryan A, Mullineaux PM, Nicholas Hewitt C. 2009. Isoprene synthesis protects transgenic tobacco plants from oxidative stress. Plant, Cell and Environment 32, 520–531.

Warneke C, Holzinger R, Hansel A, et al. 2001. Isoprene and its oxidation products measured over the tropical rainforest of Surinam. Journal of Atmospheric Chemistry 38, 167–185.

Wennberg PO, Bates KH, Crounse JD, et al. 2018. Gas-phase reactions of isoprene and its major oxidation products. Chemical Reviews 118, 3337–3390.

Wilkinson MJ, Owen SM, Possell M, Hartwell J, Gould P, Hall A, Vickers C, Nicholas Hewitt C. 2006. Circadian control of isoprene emissions from oil palm (Elaeis guineensis). The Plant Journal 47, 960–968.

Wiberley AE, Donohue AR, Westphal MM, Sharkey TD. 2009. Regulation of isoprene emission from poplar leaves throughout a day. Plant, Cell and Environment 32, 693–703.

Wolfe GM, Thornton JA, McKay M, Goldstein AH. 2011. Forest–atmosphere exchange of ozone: sensitivity to very reactive biogenic VOC emissions and implications for in-canopy photochemistry. Atmospheric Chemistry and Physics 11, 7875–7891.

Yamagishi H, Ishikawa Y. 2012. Flora of the Shirakami Natural Science Part, Hirosaki University, Bulletin of the Shirakami Institute for Environmental Sciences, Hirosaki University, SHIRAKAMI-SANCHI 1: 28–38. ISSN 2186-9219

Zuo, Z., Weraduwage SM, Huang T, Sharkey TD. 2025. How volatile isoprenoids improve plant thermotolerance. Trends in Plant Science 30, 1237–1250.

