## Supplementary Data (Table S1 and Figures S1-S11) for "Recurring daytime and nighttime modes of VOC emissions in a cool-temperate oak forest"

**Table S1.** Relative fractions in the mass spectral profiles (daytime FI, nighttime FI, daytime CT, and nighttime CT) for individual ions obtained from the 2024 and 2025 datasets.

**Fig. S1**. Study site and sampling locations.

**Fig. S2.** Schematic representation of PMF factorization.

**Fig. S3.** Comparison of time courses for the forest-interior atmospheres in the 2024 dataset from 2-, 3-, and 4-factor PMF solutions.

**Fig. S4.** GC/PTR-ToF-MS results of canopy-top atmospheres during daytime and nighttime on August 24, 2025.

**Fig. S5.** Correlation among $C_2H_5O_2^+$, $C_2H_7O_3^+$, and $C_2H_3O^+$ in the 2024 dataset.

**Fig. S6.** Time courses of concentrations of eight VOCs in the 2024 dataset.

**Fig. S7.** Correlation between $C_3OH^+$ and $C_5H_9^+$ (isoprene).

**Fig. S8.** Relationship between $RE_{\text{daytime-FI}}$, $RE_{\text{daytime-CT}}$, and $C_L \times C_T$ from September 1 to 2, 2024.

**Fig. S9.** PMF results for the two-factor solution applied to VOC ion signals measured in forest-interior atmospheres during three rain-free days in the 2025 datasets.

**Fig. S10.** PMF results for the two-factor solution applied to VOC ion signals measured in canopy-top atmospheres during three rain-free days in the 2025 datasets.

**Fig. S11.** Relationship between $RE_{\text{daytime-FI}}$, $RE_{\text{daytime-CT}}$, and $C_L \times C_T$ from September 4 to 7, 2024.

**Table S1. Relative fractions in the mass spectral profiles (daytime FI, nighttime FI, daytime CT, and nighttime CT) for individual ions obtained from the 2024 and 2025 datasets.**

| | | | 2024 | | | | 2025 | | | |
|---|---|---|---|---|---|---|---|---|---|---|
| | | | Forest-interior (FI) atmospheres | | Canopy-top (CT) atmospheres | | Forest-interior (FI) atmospheres | | Canopy-top (CT) atmospheres | |
| ID | Exact m/z | Ion formula | Daytime-FI | Nighttime-FI | Daytime-CT | Nighttime-CT | Daytime-FI | Nighttime-FI | Daytime-CT | Nighttime-CT |
| 1 | 39.02348 | C3H3+ | 0.00627399 | 0.00565692 | 0.00929861 | 0.00578456 | 0.00294222 | 0.00217803 | 0.0054137 | 0.00258638 |
| 2 | 41.03912 | C3H5+ | 0.00607564 | 0.00209644 | 0.00954433 | 0.00180706 | 0.00380398 | 0.00161489 | 0.00638187 | 0.0014862 |
| 3 | 42.033826 | C2H4N+ | 0.00336425 | 0.00374615 | 0.00270588 | 0.00340192 | 0.0033305 | 0.00288346 | 0.00260709 | 0.00301121 |
| 4 | 43.01839 | C2H3O+ | 0.0199034 | 0.00745026 | 0.01771971 | 0.01269959 | 0.02183221 | 0.00507043 | 0.01984374 | 0.00609834 |
| 5 | 43.05478 | C3H7+ | 0.00126457 | 0.00038887 | 0.00104345 | 0.00036234 | 0.00076728 | 4.52E-05 | 0.00072733 | 5.81E-05 |
| 6 | 44.049476 | C2H6N+ | 0.00011496 | 0.00019503 | 9.75E-05 | 0.00017577 | 5.33E-06 | 1.16E-05 | 6.39E-06 | 1.22E-05 |
| 7 | 45.03404 | C2H5O+ | 0.03187993 | 0.01453387 | 0.02603635 | 0.0171861 | 0.02440376 | 0.0153526 | 0.02255329 | 0.0206815 |
| 8 | 46.02874 | CH4NO+ | 0.00379715 | 0.01177367 | 0.00394118 | 0.01279996 | 0.00053152 | 0.00109098 | 0.00136747 | 0.00114923 |
| 9 | 47.049141 | C2H7O+ | 0.00749995 | 0.00019968 | 0.00609089 | 0.00090001 | 0.00413069 | 0.00391269 | 0.00355477 | 0.00389334 |
| 10 | 48.008005 | H2NO2+ | 0.00481779 | 0.00437238 | 0.00468021 | 0.00582312 | 0.00027213 | 0.00044466 | 0.00038546 | 0.00065883 |
| 11 | 49.010648 | CH5S+ | 0.00043909 | 2.15E-05 | 0.00030728 | 4.50E-06 | 0.00053672 | 3.21E-05 | 0.00030285 | 4.85E-05 |
| 12 | 51.02348 | C4H3+ | 1.19E-05 | 0.00037358 | 1.82E-05 | 0.00029193 | 2.36E-08 | 1.42E-05 | 3.80E-08 | 1.29E-05 |
| 13 | 51.044056 | CH7O2+ | 0.00514072 | 0.00172915 | 0.0045572 | 0.00190112 | 0.00955639 | 0.00187076 | 0.00760432 | 0.00232304 |
| 14 | 51.994366 | C3O+ | 0.00083033 | 0.00017617 | 0.00080199 | 0.00025177 | 5.13E-07 | 1.50E-06 | 5.31E-07 | 1.45E-06 |
| 15 | 52.039305 | H6NO2+ | 0.00122679 | 0.00428852 | 0.00135276 | 0.00452346 | 6.40E-06 | 3.73E-05 | 6.60E-06 | 3.67E-05 |
| 16 | 53.002191 | C3HO+ | 0.00559921 | 0.00326505 | 0.00836331 | 0.00275821 | 0.00467765 | 0.00361889 | 0.00739775 | 0.00338687 |
| 17 | 53.03912 | C4H5+ | 0.00179495 | 0.00116983 | 0.00249777 | 0.00094608 | 0.00137708 | 0.00141775 | 0.00203618 | 0.001361 |
| 18 | 53.991567 | <unknown0012> | 0.00026477 | 0.00015684 | 0.00027308 | 0.00016377 | 5.28E-07 | 1.68E-06 | 4.26E-07 | 1.59E-06 |
| 19 | 54.033826 | C3H4N+ | 0.00067238 | 0.00051884 | 0.00055846 | 0.00049017 | 1.36E-05 | 1.15E-05 | 8.20E-06 | 1.13E-05 |
| 20 | 57.069877 | C4H9+ | 0.01150409 | 0.01081156 | 0.00967415 | 0.01167682 | 0.00753044 | 0.0146996 | 0.00734714 | 0.01509606 |
| 21 | 58.02874 | C2H4NO+ | 0.00034639 | 0.00062775 | 0.00037032 | 0.00056399 | 5.46E-06 | 1.87E-06 | 7.20E-06 | 1.69E-06 |
| 22 | 58.041316 | C3H6O+ | 3.22E-06 | 5.62E-07 | 6.92E-06 | 6.34E-08 | 0.00050477 | 6.08E-05 | 0.00057464 | 0.00012336 |
| 23 | 58.069198 | <unknown0021> | 2.70E-07 | 7.87E-07 | 2.41E-07 | 6.34E-07 | 3.70E-07 | 1.22E-06 | 2.73E-07 | 1.16E-06 |
| 24 | 59.04969 | C3H7O+ | 0.27194373 | 0.2662174 | 0.23942894 | 0.30736092 | 0.3884483 | 0.36360269 | 0.30152219 | 0.35904668 |
| 25 | 60.04439 | C2H6NO+ | 0.00202958 | 0.00444302 | 0.00175281 | 0.00451072 | 0.00058179 | 0.00083926 | 0.00102749 | 0.00117986 |
| 26 | 61.02895 | C2H5O2+ | 0.03369561 | 0.01463572 | 0.02953028 | 0.02727874 | 0.04498224 | 0.00274181 | 0.04110324 | 0.00458851 |
| 27 | 61.039639 | CH5N2O+ | 1.77E-07 | 6.00E-07 | 1.26E-07 | 4.65E-07 | 3.14E-07 | 1.05E-06 | 2.33E-07 | 9.90E-07 |
| 28 | 62.023655 | CH4NO2+ | 0.00014552 | 0.00059512 | 0.00013012 | 0.00048637 | 1.05E-06 | 1.74E-05 | 8.07E-07 | 2.48E-05 |
| 29 | 63.02348 | C5H3+ | 1.84E-06 | 2.19E-06 | 1.86E-06 | 1.22E-06 | 7.88E-05 | 4.68E-06 | 0.00010262 | 5.59E-06 |
| 30 | 63.044056 | C2H7O2+ | 0.00457692 | 0.00215291 | 0.00388957 | 0.00265583 | 0.00276385 | 0.00196128 | 0.00251164 | 0.0021317 |
| 31 | 64.039305 | CH6NO2+ | 0.00018755 | 0.00043451 | 0.0001589 | 0.0004461 | 5.77E-07 | 1.99E-06 | 5.25E-07 | 1.91E-06 |
| 32 | 65.03912 | C5H5+ | 0 | 1.27E-05 | 0 | 7.39E-06 | 3.07E-06 | 4.41E-06 | 3.42E-06 | 1.04E-05 |
| 33 | 65.059706 | C2H9O2+ | 0.0033473 | 0 | 0.0030027 | 0 | 0.0022928 | 0.00054541 | 0.00200698 | 0.00067714 |
| 34 | 66.046402 | C5H6+ | 0.00014397 | 3.86E-05 | 0.00026566 | 1.36E-05 | 4.90E-06 | 8.10E-06 | 6.00E-05 | 1.46E-06 |
| 35 | 67.01839 | C4H3O+ | 1.75E-07 | 6.12E-07 | 1.24E-07 | 4.89E-07 | 3.19E-07 | 1.16E-06 | 2.31E-07 | 1.08E-06 |
| 36 | 67.05477 | C5H7+ | 0.0090535 | 0.00406126 | 0.01506279 | 0.00311513 | 0.00615002 | 0.00391474 | 0.01136583 | 0.00369618 |
| 37 | 68.049476 | C4H6N+ | 0 | 2.29E-05 | 0 | 8.37E-06 | 3.09E-07 | 1.95E-06 | 2.12E-07 | 1.70E-06 |
| 38 | 68.062052 | C5H8+ | 5.36E-05 | 0 | 0.00018111 | 0 | 0.00304226 | 0 | 0.00614958 | 0 |
| 39 | 69.03404 | C4H5O+ | 6.68E-08 | 1.94E-06 | 7.40E-08 | 9.88E-07 | 2.16E-07 | 3.69E-06 | 2.01E-07 | 1.89E-06 |
| 40 | 69.07043 | C5H9+ | 0.13430402 | 0.01577984 | 0.24987613 | 0.01132379 | 0.12971832 | 0.01551855 | 0.25783428 | 0.014138 |
| 41 | 70.004931 | C3H2O2+ | 8.30E-05 | 9.71E-05 | 6.95E-05 | 9.51E-05 | 7.24E-07 | 7.36E-06 | 2.20E-07 | 6.34E-06 |
| 42 | 70.041316 | C4H6O+ | 0.00010591 | 0.00030218 | 0 | 9.64E-06 | 3.28E-06 | 2.62E-06 | 2.46E-07 | 2.75E-06 |
| 43 | 71.01331 | C3H3O2+ | 0 | 0.00085398 | 0 | 0.00078712 | 4.30E-05 | 0.0009655 | 0.000124 | 0.00137046 |
| 44 | 71.04969 | C4H7O+ | 0.07447787 | 0 | 0.06641995 | 0.00279772 | 0.0703379 | 0.00158343 | 0.05874973 | 0.00368263 |
| 45 | 71.08607 | C5H11+ | 0 | 9.93E-06 | 0 | 5.55E-06 | 0 | 0.00502673 | 0 | 0.00547025 |
| 46 | 72.04439 | C3H6NO+ | 0.00032355 | 0.00046363 | 0.00032943 | 0.00040697 | 6.36E-07 | 2.58E-06 | 5.19E-07 | 2.88E-06 |
| 47 | 72.080776 | C4H10N+ | 1.77E-07 | 6.47E-07 | 1.24E-07 | 5.18E-07 | 3.48E-07 | 1.41E-06 | 2.67E-07 | 1.35E-06 |
| 48 | 74.023655 | C2H4NO2+ | 0.00051716 | 0.00051455 | 0.00052984 | 0.00057263 | 0.00028598 | 0.00037584 | 0.00030598 | 0.00045591 |
| 49 | 74.06004 | C3H8NO+ | 0.00175512 | 0.00158834 | 0.0012693 | 0.00137307 | 0.00111064 | 0.00071071 | 0.00111087 | 0.00075839 |
| 50 | 75.00822 | C2H3O3+ | 1.76E-07 | 6.05E-07 | 1.25E-07 | 4.81E-07 | 3.11E-07 | 1.14E-06 | 2.30E-07 | 1.07E-06 |

**Table S1.** Continued.

| | | | 2024 | | | | 2025 | | | |
|---|---|---|---|---|---|---|---|---|---|---|
| | | | Forest-interior (FI) atmospheres | | Canopy-top (CT) atmospheres | | Forest-interior (FI) atmospheres | | Canopy-top (CT) atmospheres | |
| ID | Exact m/z | Ion formula | Daytime-FI | Nighttime-FI | Daytime-CT | Nighttime-CT | Daytime-FI | Nighttime-FI | Daytime-CT | Nighttime-CT |
| 51 | 75.0446 | C3H7O2+ | 0.01895785 | 0.00322678 | 0.01751355 | 0.00462638 | 0.03090961 | 0 | 0.02645667 | 0.00101322 |
| 52 | 76.039305 | C2H6NO2+ | 0.00028228 | 0.00046688 | 0.00024367 | 0.00043798 | 1.62E-06 | 3.75E-06 | 1.70E-06 | 6.34E-06 |
| 53 | 76.07569 | C3H10NO+ | 2.84E-07 | 7.43E-07 | 2.13E-07 | 6.58E-07 | 3.25E-07 | 1.17E-06 | 2.41E-07 | 1.09E-06 |
| 54 | 77.02387 | C2H5O3+ | 1.67E-06 | 0 | 1.24E-05 | 0 | 0.00851615 | 0 | 0.00661126 | 0 |
| 55 | 77.059706 | C3H9O2+ | 0.02851868 | 0.03425402 | 0.02443527 | 0.0376164 | 0.01340286 | 0.01257734 | 0.01045076 | 0.01310435 |
| 56 | 78.054955 | C2H8NO2+ | 0.00033961 | 0.00075304 | 0.00026117 | 0.00067901 | 5.27E-06 | 0.00012059 | 6.19E-06 | 6.94E-05 |
| 57 | 79.03897 | C2H7O3+ | 0.00773451 | 0.00385627 | 0.00727253 | 0.00757776 | 0.00439294 | 0 | 0.00426675 | 0 |
| 58 | 79.05477 | C6H7+ | 0.00339215 | 0.0108458 | 0.00181877 | 0.00811814 | 0.0040355 | 0.01780891 | 0.0030331 | 0.01702731 |
| 59 | 80.049476 | C5H6N+ | 0.00059966 | 0.00081692 | 0.00049312 | 0.00075109 | 0.00042329 | 1.11E-05 | 0.00037422 | 2.12E-05 |
| 60 | 81.03404 | C5H5O+ | 0.00010177 | 0 | 9.53E-05 | 4.99E-07 | 0.00022133 | 1.44E-05 | 0.00022273 | 4.86E-05 |
| 61 | 81.07043 | C6H9+ | 0.01542682 | 0.09254855 | 0.01139855 | 0.0731044 | 0.01239494 | 0.11957735 | 0.01009194 | 0.11457554 |
| 62 | 83.04969 | C5H7O+ | 0.00593245 | 0.00242594 | 0.00505699 | 0.00240482 | 0.00634586 | 0.00206774 | 0.00522538 | 0.00255631 |
| 63 | 83.08607 | C6H11+ | 0.00819366 | 0.0063356 | 0.00684723 | 0.0062523 | 0.00431077 | 0.0045366 | 0.00467036 | 0.00511816 |
| 64 | 84.04439 | C4H6NO+ | 0.00016497 | 0.00038501 | 0.0001505 | 0.00033678 | 5.73E-06 | 6.18E-06 | 7.03E-06 | 5.96E-06 |
| 65 | 84.080776 | C5H10N+ | 7.52E-07 | 2.62E-06 | 5.23E-07 | 2.09E-06 | 0.00027998 | 0.00107094 | 0.00017244 | 0.00077031 |
| 66 | 85.02895 | C4H5O2+ | 0.00134464 | 0.00087036 | 0.00111423 | 0.00095386 | 0.00187312 | 0.0011538 | 0.00165095 | 0.00128261 |
| 67 | 85.06534 | C5H9O+ | 0.00523275 | 0.0026493 | 0.00471669 | 0.00247196 | 0.0054949 | 0.00350812 | 0.00468239 | 0.00377222 |
| 68 | 85.10172 | C6H13+ | 0.00066112 | 0.00084033 | 0.0001974 | 0.00094175 | 0.00074517 | 0.00128322 | 0.00069153 | 0.00157445 |
| 69 | 86.023655 | C3H4NO2+ | 5.52E-05 | 0.00017145 | 6.48E-05 | 0.00013694 | 1.08E-06 | 2.01E-06 | 8.22E-07 | 1.99E-06 |
| 70 | 86.06004 | C4H8NO+ | 0.0001444 | 0.00023269 | 0.00013933 | 0.0002141 | 1.98E-06 | 3.02E-06 | 2.15E-06 | 2.83E-06 |
| 71 | 87.02348 | C7H3+ | 1.72E-07 | 8.12E-07 | 1.15E-07 | 6.61E-07 | 3.10E-07 | 1.18E-06 | 2.29E-07 | 1.12E-06 |
| 72 | 87.0446 | C4H7O2+ | 0.00788269 | 0.00100735 | 0.00658605 | 0.00177994 | 0.01085301 | 0.00131576 | 0.00962528 | 0.00243318 |
| 73 | 87.08099 | C5H11O+ | 0.00228308 | 1.27E-05 | 0.00172409 | 8.14E-06 | 0.00419539 | 0.00638403 | 0.00309466 | 0.00643263 |
| 74 | 88.039305 | C3H6NO2+ | 0.00037875 | 0.00049327 | 0.00035286 | 0.00051381 | 0.00021496 | 0.00014574 | 0.00023836 | 0.00021957 |
| 75 | 88.07569 | C4H10NO+ | 5.00E-06 | 6.26E-07 | 3.13E-06 | 7.60E-07 | 2.21E-06 | 2.87E-06 | 2.04E-06 | 2.66E-06 |
| 76 | 89.02387 | C3H5O3+ | 0.00037576 | 0.00100644 | 0.00079204 | 0.00093225 | 4.07E-06 | 1.75E-06 | 6.41E-06 | 0.00094177 |
| 77 | 89.03912 | C7H5+ | 1.77E-07 | 6.11E-07 | 1.24E-07 | 4.89E-07 | 3.14E-07 | 1.06E-06 | 2.33E-07 | 9.92E-07 |
| 78 | 89.06026 | C4H9O2+ | 0.02146492 | 0.00259916 | 0.01680018 | 0.00327829 | 0.01343567 | 0.00385467 | 0.01274964 | 0.00390992 |
| 79 | 90.046402 | C7H6+ | 8.48E-05 | 0.00021579 | 6.20E-05 | 0.00018315 | 7.30E-07 | 2.48E-06 | 3.72E-07 | 5.67E-06 |
| 80 | 90.066188 | C2H8N3O+ | 2.45E-07 | 6.13E-07 | 1.80E-07 | 4.96E-07 | 4.92E-07 | 1.30E-06 | 3.77E-07 | 1.22E-06 |
| 81 | 91.00314 | C2H3O4+ | 1.75E-07 | 6.91E-07 | 1.20E-07 | 5.54E-07 | 3.12E-07 | 1.27E-06 | 2.31E-07 | 1.19E-06 |
| 82 | 91.01839 | C6H3O+ | 1.78E-07 | 5.98E-07 | 1.25E-07 | 4.80E-07 | 3.13E-07 | 1.13E-06 | 2.32E-07 | 1.06E-06 |
| 83 | 91.03952 | C3H7O3+ | 0.00119838 | 1.11E-05 | 0.00103813 | 3.60E-05 | 0.00322466 | 1.99E-06 | 0.00245693 | 8.94E-06 |
| 84 | 91.05477 | C7H7+ | 0.00391277 | 0.00897532 | 0.00267296 | 0.00812069 | 3.42E-06 | 0.00629621 | 2.87E-06 | 0.0058766 |
| 85 | 91.075356 | C4H11O2+ | 0.00812688 | 0.00267045 | 0.00661263 | 0.00263535 | 0.00360714 | 2.34E-05 | 0.0033109 | 1.36E-05 |
| 86 | 92.034219 | C2H6NO3+ | 0.0001078 | 0.00036448 | 8.61E-05 | 0.00029796 | 1.16E-06 | 1.47E-06 | 9.49E-07 | 1.43E-06 |
| 87 | 92.062052 | C7H8+ | 1.64E-06 | 3.80E-06 | 1.18E-06 | 4.08E-06 | 1.10E-05 | 0.0022089 | 2.29E-05 | 0.00216289 |
| 88 | 93.01878 | C2H5O4+ | 2.31E-05 | 0 | 1.03E-06 | 1.09E-06 | 1.84E-05 | 0.00025816 | 5.20E-06 | 1.45E-05 |
| 89 | 93.03404 | C6H5O+ | 1.86E-07 | 5.81E-07 | 1.33E-07 | 4.66E-07 | 3.57E-07 | 8.32E-07 | 3.26E-07 | 8.08E-07 |
| 90 | 93.07043 | C7H9+ | 0.01646141 | 0.02542734 | 0.01293008 | 0.02574075 | 0.01098121 | 0.02970092 | 0.00873468 | 0.03018149 |
| 91 | 95.01331 | C5H3O2+ | 1.89E-07 | 5.79E-07 | 1.42E-07 | 4.63E-07 | 6.80E-07 | 1.01E-06 | 5.59E-07 | 9.62E-07 |
| 92 | 95.04969 | C6H7O+ | 0.00569124 | 0.00713475 | 0.00430096 | 0.0068306 | 0.00277521 | 0.00358964 | 0.00249003 | 0.00478303 |
| 93 | 95.08607 | C7H11+ | 1.13E-06 | 1.66E-05 | 0 | 2.11E-05 | 0.00143063 | 0.01094644 | 0.0014189 | 0.01056669 |
| 94 | 96.089329 | C2H12N2O2+ | 7.40E-07 | 1.60E-06 | 5.27E-07 | 1.35E-06 | 6.49E-07 | 1.89E-06 | 5.51E-07 | 1.81E-06 |
| 95 | 97.02895 | C5H5O2+ | 0.00166305 | 0.00203863 | 0.00140941 | 0.00234263 | 0.00137718 | 0.00083522 | 0.00124962 | 0.00117969 |
| 96 | 97.06534 | C6H9O+ | 0.00175116 | 0.00189968 | 0.00144471 | 0.00184303 | 0.0019086 | 0.00240092 | 0.00155923 | 0.0025011 |
| 97 | 97.10172 | C7H13+ | 0.00208084 | 2.86E-06 | 0.00184204 | 1.87E-05 | 9.59E-06 | 1.21E-05 | 0.00085439 | 2.10E-05 |
| 98 | 98.034888 | C3H4N3O+ | 7.80E-05 | 0.00016917 | 5.21E-05 | 0.00013712 | 3.74E-05 | 1.42E-05 | 1.60E-05 | 1.80E-05 |
| 99 | 98.06004 | C5H8NO+ | 8.87E-05 | 1.55E-05 | 0.00014778 | 2.51E-05 | 8.53E-07 | 1.94E-06 | 8.65E-07 | 1.78E-06 |
| 100 | 98.109002 | C7H14+ | 2.70E-07 | 7.61E-07 | 1.83E-07 | 6.22E-07 | 5.39E-07 | 1.65E-06 | 4.13E-07 | 1.56E-06 |

**Table S1.** Continued.

| | | | 2024 | | | | 2025 | | | |
|---|---|---|---|---|---|---|---|---|---|---|
| | | | Forest-interior (FI) atmospheres | | Canopy-top (CT) atmospheres | | Forest-interior (FI) atmospheres | | Canopy-top (CT) atmospheres | |
| ID | Exact m/z | Ion formula | Daytime-FI | Nighttime-FI | Daytime-CT | Nighttime-CT | Daytime-FI | Nighttime-FI | Daytime-CT | Nighttime-CT |
| 101 | 99.02348 | C8H3+ | 1.98E-07 | 7.15E-07 | 1.37E-07 | 5.83E-07 | 0 | 3.34E-05 | 0 | 4.77E-05 |
| 102 | 99.0446 | C5H7O2+ | 0.00507394 | 0.00144624 | 0.0043668 | 0.00157679 | 0.0097936 | 0.00091636 | 0.00789676 | 0.00113226 |
| 103 | 99.08099 | C6H11O+ | 0.00560719 | 0.003443 | 0.00440451 | 0.00333086 | 0.00458216 | 0.00235723 | 0.00357679 | 0.00295941 |
| 104 | 99.11738 | C7H15+ | 1.77E-07 | 5.89E-07 | 1.26E-07 | 4.72E-07 | 3.18E-07 | 1.09E-06 | 2.41E-07 | 1.04E-06 |
| 105 | 100.039305 | C4H6NO2+ | 0.00022361 | 0.000152 | 0.00019859 | 0.00015648 | 0.00022564 | 0 | 0.00018241 | 0 |
| 106 | 100.07569 | C5H10NO+ | 2.37E-06 | 4.34E-06 | 1.36E-06 | 3.05E-06 | 6.98E-07 | 1.47E-06 | 5.45E-07 | 1.48E-06 |
| 107 | 101.02387 | C4H5O3+ | 0.00019846 | 0.0003252 | 0.00017708 | 0.00037467 | 0.00045445 | 0.00149248 | 0.00055211 | 0.00163778 |
| 108 | 101.03912 | C8H5+ | 1.78E-07 | 5.83E-07 | 1.26E-07 | 4.68E-07 | 3.14E-07 | 1.06E-06 | 2.33E-07 | 9.90E-07 |
| 109 | 101.06026 | C5H9O2+ | 0.01163731 | 0.00268311 | 0.00987877 | 0.00290821 | 0.01742902 | 0.00104578 | 0.01429398 | 0.001687 |
| 110 | 101.09664 | C6H13O+ | 1.62E-07 | 1.10E-06 | 9.79E-08 | 9.88E-07 | 0 | 0.00077547 | 0 | 0.00225394 |
| 111 | 102.018569 | C3H4NO3+ | 2.74E-05 | 5.13E-05 | 1.70E-05 | 7.70E-05 | 7.14E-07 | 2.01E-06 | 5.53E-07 | 2.08E-06 |
| 112 | 102.054955 | C4H8NO2+ | 2.73E-05 | 0.00012243 | 5.69E-05 | 0.00010799 | 6.85E-07 | 1.82E-06 | 5.97E-07 | 1.74E-06 |
| 113 | 102.09134 | C5H12NO+ | 0.00112606 | 0.00326163 | 0.00079995 | 0.00254887 | 0.00062091 | 0.00159022 | 0.00048031 | 0.00144135 |
| 114 | 103.01839 | C7H3O+ | 1.78E-07 | 5.82E-07 | 1.26E-07 | 4.67E-07 | 3.18E-07 | 1.12E-06 | 2.36E-07 | 1.05E-06 |
| 115 | 103.03952 | C4H7O3+ | 0.0300875 | 0.06452179 | 0.02312227 | 0.06144287 | 0.00249211 | 0.00053456 | 0.00200586 | 0.00063184 |
| 116 | 103.0759 | C5H11O2+ | 1.78E-07 | 5.82E-07 | 1.27E-07 | 4.67E-07 | 0.00133468 | 0.00054309 | 0.001165 | 0.00061711 |
| 117 | 104.042259 | <unknown0071> | 0 | 0.00031743 | 0.00018713 | 0.00030891 | 0.00028495 | 0 | 0.00023821 | 1.41E-06 |
| 118 | 104.070605 | C4H10NO2+ | 1.84E-07 | 5.86E-07 | 1.32E-07 | 4.70E-07 | 5.92E-07 | 1.39E-06 | 4.49E-07 | 1.31E-06 |
| 119 | 105.01878 | C3H5O4+ | 0.00017587 | 0.00048591 | 0.00014973 | 0.00046528 | 0.00014987 | 0.0007154 | 0.00017963 | 0.00095706 |
| 120 | 105.03404 | C7H5O+ | 1.83E-07 | 5.96E-07 | 1.28E-07 | 4.81E-07 | 3.20E-07 | 1.08E-06 | 2.39E-07 | 1.01E-06 |
| 121 | 105.05517 | C4H9O3+ | 0.00302706 | 0.00168663 | 0.00248668 | 0.00169831 | 0.00328038 | 0.0001041 | 0.0026148 | 0.00026417 |
| 122 | 105.07043 | C8H9+ | 2.80E-07 | 1.02E-06 | 2.04E-07 | 8.59E-07 | 5.52E-07 | 5.14E-06 | 4.68E-07 | 5.04E-06 |
| 123 | 105.091006 | C5H13O2+ | 0.00178116 | 6.76E-06 | 0.00141275 | 7.38E-06 | 0.00059466 | 0 | 0.00031766 | 4.84E-06 |
| 124 | 106.04987 | C3H8NO3+ | 8.45E-05 | 0.00021131 | 6.61E-05 | 0.0001823 | 5.02E-07 | 1.37E-06 | 3.98E-07 | 1.31E-06 |
| 125 | 106.077702 | C8H10+ | 5.82E-07 | 9.62E-07 | 4.16E-07 | 7.95E-07 | 1.02E-06 | 3.75E-06 | 8.41E-07 | 3.49E-06 |
| 126 | 106.094124 | <unknown0076> | 2.80E-07 | 8.62E-07 | 1.98E-07 | 6.79E-07 | 4.18E-07 | 1.33E-06 | 3.11E-07 | 1.24E-06 |
| 127 | 107.04969 | C7H7O+ | 0.00176794 | 0.00166843 | 0.00174836 | 0.00229729 | 0.00132439 | 0.00031986 | 0.00051702 | 0 |
| 128 | 107.08607 | C8H11+ | 0.00735111 | 0.00997687 | 0.00517074 | 0.0081602 | 0.00422482 | 0.01091196 | 0.00347023 | 0.00966014 |
| 129 | 108.087386 | <unknown0079> | 4.68E-06 | 8.83E-06 | 2.92E-06 | 7.63E-06 | 1.31E-06 | 4.26E-06 | 9.84E-07 | 5.03E-06 |
| 130 | 109.02895 | C6H5O2+ | 0.00497793 | 0.00334596 | 0.00444005 | 0.00467481 | 0.002678 | 0.00407767 | 0.00332392 | 0.00800999 |
| 131 | 109.06534 | C7H9O+ | 1.79E-07 | 5.82E-07 | 1.26E-07 | 4.67E-07 | 3.41E-07 | 1.13E-06 | 2.80E-07 | 1.02E-06 |
| 132 | 109.10172 | C8H13+ | 0.00376559 | 0.00688946 | 0.0028739 | 0.00563067 | 0.00232959 | 0.00793572 | 0.00206477 | 0.00744276 |
| 133 | 110.063412 | C3H12NOS+ | 3.31E-07 | 1.19E-06 | 1.93E-07 | 8.39E-07 | 7.07E-07 | 1.88E-06 | 5.50E-07 | 1.65E-06 |
| 134 | 111.02348 | C9H3+ | 1.78E-07 | 5.82E-07 | 1.26E-07 | 4.67E-07 | 3.14E-07 | 1.06E-06 | 2.33E-07 | 9.91E-07 |
| 135 | 111.0446 | C6H7O2+ | 0.00075305 | 0 | 0.00078811 | 8.17E-05 | 0.00099132 | 0.00141538 | 0.00121025 | 0.00209984 |
| 136 | 111.08099 | C7H11O+ | 1.93E-07 | 5.79E-07 | 1.48E-07 | 4.66E-07 | 1.33E-05 | 1.05E-07 | 1.06E-05 | 4.18E-07 |
| 137 | 111.11738 | C8H15+ | 0.00171854 | 0.00175589 | 0.00131953 | 0.00162631 | 0.00027528 | 0.00117778 | 0.00052552 | 0.00132975 |
| 138 | 112.079062 | C3H14NOS+ | 5.53E-05 | 3.73E-05 | 4.85E-05 | 3.92E-05 | 1.83E-06 | 4.66E-06 | 1.38E-06 | 4.65E-06 |
| 139 | 113.02387 | C5H5O3+ | 0.00012134 | 1.47E-05 | 6.72E-06 | 1.05E-05 | 0.00171201 | 0 | 0.00141164 | 5.19E-05 |
| 140 | 113.03912 | C9H5+ | 1.82E-07 | 5.90E-07 | 1.29E-07 | 4.73E-07 | 3.14E-07 | 1.06E-06 | 2.33E-07 | 9.99E-07 |
| 141 | 113.06026 | C6H9O2+ | 0.00227523 | 0.00133928 | 0.0019284 | 0.00151014 | 0.00304221 | 0.0023228 | 0.0026523 | 0.00287027 |
| 142 | 113.09664 | C7H13O+ | 1.90E-07 | 5.82E-07 | 1.46E-07 | 4.70E-07 | 4.38E-07 | 1.12E-06 | 3.81E-07 | 1.09E-06 |
| 143 | 114.018569 | C4H4NO3+ | 4.99E-05 | 0.00011993 | 3.04E-05 | 0.0001016 | 7.13E-07 | 2.41E-06 | 5.28E-07 | 2.53E-06 |
| 144 | 114.054955 | C5H8NO2+ | 2.51E-06 | 5.26E-06 | 1.29E-06 | 4.42E-06 | 5.65E-07 | 1.61E-06 | 5.03E-07 | 1.51E-06 |
| 145 | 114.09134 | C6H12NO+ | 3.42E-07 | 6.05E-07 | 4.91E-07 | 4.88E-07 | 6.99E-07 | 1.50E-06 | 3.30E-07 | 1.13E-06 |
| 146 | 115.01839 | C8H3O+ | 4.91E-07 | 1.11E-06 | 3.51E-07 | 8.97E-07 | 6.50E-07 | 3.06E-05 | 1.95E-07 | 5.83E-05 |
| 147 | 115.03952 | C5H7O3+ | 0.0005671 | 0.00079005 | 0.00047006 | 0.00076867 | 0.0014539 | 0.00051112 | 0.00113368 | 0.00054309 |
| 148 | 115.05477 | C9H7+ | 1.82E-07 | 6.08E-07 | 1.28E-07 | 4.85E-07 | 3.20E-07 | 1.17E-06 | 2.37E-07 | 1.09E-06 |
| 149 | 115.0759 | C6H11O2+ | 0.00253478 | 0.00178874 | 0.00206693 | 0.00171931 | 0.00307708 | 0.00240776 | 0.00266881 | 0.00253273 |
| 150 | 115.11229 | C7H15O+ | 2.02E-07 | 6.28E-07 | 1.58E-07 | 5.30E-07 | 7.19E-07 | 2.30E-06 | 8.27E-07 | 3.23E-06 |

**Table S1.** Continued.

| | | | 2024 | | | | 2025 | | | |
|---|---|---|---|---|---|---|---|---|---|---|
| | | | Forest-interior (FI) atmospheres | | Canopy-top (CT) atmospheres | | Forest-interior (FI) atmospheres | | Canopy-top (CT) atmospheres | |
| ID | Exact m/z | Ion formula | Daytime-FI | Nighttime-FI | Daytime-CT | Nighttime-CT | Daytime-FI | Nighttime-FI | Daytime-CT | Nighttime-CT |
| 151 | 116.034219 | C4H6NO3+ | 6.96E-05 | 0.00014743 | 5.26E-05 | 0.00012642 | 1.15E-06 | 2.12E-06 | 1.23E-06 | 1.97E-06 |
| 152 | 116.070605 | C5H10NO2+ | 8.76E-07 | 1.56E-06 | 6.33E-07 | 1.31E-06 | 5.45E-07 | 1.45E-06 | 4.39E-07 | 1.36E-06 |
| 153 | 116.10699 | C6H14NO+ | 5.34E-06 | 6.68E-06 | 5.18E-06 | 5.26E-06 | 1.70E-06 | 6.43E-06 | 1.38E-06 | 5.37E-06 |
| 154 | 117.03404 | C8H5O+ | 3.32E-06 | 0.00030722 | 0 | 0.0002645 | 6.69E-05 | 0.0003758 | 4.70E-05 | 0.00042407 |
| 155 | 117.05517 | C5H9O3+ | 0.00301652 | 0 | 0.00269457 | 1.42E-05 | 0.00392678 | 0 | 0.00310673 | 0 |
| 156 | 117.07043 | C9H9+ | 1.75E-07 | 6.45E-07 | 1.22E-07 | 5.11E-07 | 3.07E-07 | 1.38E-06 | 2.29E-07 | 1.30E-06 |
| 157 | 117.09155 | C6H13O2+ | 0.00373536 | 0.00306851 | 0.00282052 | 0.00288177 | 0.00127265 | 0.00087983 | 0.00100252 | 0.0010766 |
| 158 | 118.04987 | C4H8NO3+ | 0.00014243 | 0.00022176 | 0.00011928 | 0.00019973 | 1.83E-06 | 1.77E-06 | 1.81E-06 | 1.53E-06 |
| 159 | 118.086255 | C5H12NO2+ | 5.92E-07 | 7.30E-07 | 5.29E-07 | 6.01E-07 | 7.09E-07 | 1.47E-06 | 5.66E-07 | 1.41E-06 |
| 160 | 119.01331 | C7H3O2+ | 1.78E-07 | 5.82E-07 | 1.26E-07 | 4.67E-07 | 3.16E-07 | 1.06E-06 | 2.34E-07 | 9.97E-07 |
| 161 | 119.03443 | C4H7O4+ | 0.00052912 | 0.000103 | 0.00047358 | 0.00015241 | 0.00066436 | 0.00081092 | 0.00059314 | 0.00088925 |
| 162 | 119.07082 | C5H11O3+ | 2.92E-07 | 5.84E-07 | 2.05E-07 | 4.76E-07 | 6.12E-06 | 6.56E-07 | 4.27E-06 | 7.85E-07 |
| 163 | 119.08607 | C9H11+ | 1.10E-06 | 0.00321933 | 0 | 0.00293475 | 1.29E-06 | 4.93E-05 | 1.33E-06 | 0.00063985 |
| 164 | 120.031548 | <unknown0095> | 0.00028208 | 0.00068206 | 0.00020243 | 0.00062615 | 2.33E-06 | 6.51E-06 | 2.15E-06 | 7.90E-06 |
| 165 | 121.02895 | C7H5O2+ | 1.78E-07 | 5.82E-07 | 1.26E-07 | 4.67E-07 | 0.00013907 | 0.00047932 | 0.00012665 | 0.00057557 |
| 166 | 121.05009 | C4H9O4+ | 0.04414783 | 0.11519272 | 0.03207099 | 0.10667431 | 3.13E-05 | 0 | 2.16E-05 | 0 |
| 167 | 121.10172 | C9H13+ | 6.01E-07 | 5.90E-07 | 3.30E-07 | 5.07E-07 | 0.00211266 | 0.00656237 | 0.00185512 | 0.00641919 |
| 168 | 123.00822 | C6H3O3+ | 0 | 0.00019243 | 5.55E-05 | 0.00016696 | 4.69E-05 | 0.00022135 | 3.43E-05 | 0.00025473 |
| 169 | 123.0446 | C7H7O2+ | 0.00049025 | 0.00055312 | 0.00038367 | 0.00049915 | 0.00032445 | 0.00014478 | 0.0002922 | 0.00022497 |
| 170 | 123.08099 | C8H11O+ | 2.17E-07 | 6.78E-07 | 1.61E-07 | 5.27E-07 | 2.73E-06 | 2.58E-06 | 1.72E-06 | 2.23E-06 |
| 171 | 123.11738 | C9H15+ | 0.0015359 | 0.00288303 | 0.00115315 | 0.00259705 | 0.00107531 | 0.00335602 | 0.00160619 | 0.0033423 |
| 172 | 125.03912 | C10H5+ | 0.00117499 | 0.00109741 | 0.0009621 | 0.00152626 | 0.00070762 | 0.00094756 | 0.00071103 | 0.00125114 |
| 173 | 125.06026 | C7H9O2+ | 2.61E-07 | 5.69E-07 | 2.36E-07 | 4.82E-07 | 3.14E-07 | 1.05E-06 | 2.33E-07 | 9.90E-07 |
| 174 | 125.09664 | C8H13O+ | 0.00104389 | 0 | 0.00090291 | 1.44E-05 | 0.00139702 | 7.07E-05 | 0.00109552 | 3.67E-05 |
| 175 | 125.13303 | C9H17+ | 0.00081691 | 0.00011242 | 0.00044237 | 6.92E-05 | 2.74E-06 | 3.43E-05 | 0.00050412 | 0.00024488 |
| 176 | 126.099894 | C3H14N2O3+ | 4.35E-07 | 1.26E-06 | 3.26E-07 | 9.86E-07 | 5.54E-07 | 1.58E-06 | 4.20E-07 | 1.49E-06 |
| 177 | 126.140302 | C9H18+ | 2.31E-07 | 7.04E-07 | 1.64E-07 | 5.65E-07 | 4.06E-07 | 1.31E-06 | 3.28E-07 | 1.26E-06 |
| 178 | 127.03897 | C6H7O3+ | 0.00136012 | 0.00480533 | 0.00111572 | 0.00431126 | 0.00090517 | 0.0021388 | 0.00104076 | 0.00391434 |
| 179 | 127.11229 | C8H15O+ | 2.91E-07 | 5.61E-07 | 2.52E-07 | 4.40E-07 | 0.00200702 | 7.93E-05 | 0.0017693 | 0.00024437 |
| 180 | 127.14867 | C9H19+ | 2.48E-07 | 5.79E-07 | 1.71E-07 | 4.72E-07 | 3.21E-07 | 1.12E-06 | 2.48E-07 | 1.09E-06 |
| 181 | 128.070605 | C6H10NO2+ | 1.88E-07 | 6.25E-07 | 1.34E-07 | 5.00E-07 | 1.18E-06 | 1.99E-06 | 1.08E-06 | 1.81E-06 |
| 182 | 128.114945 | <unknown0108> | 7.55E-07 | 6.24E-07 | 5.39E-07 | 5.32E-07 | 4.31E-07 | 1.36E-06 | 3.22E-07 | 1.28E-06 |
| 183 | 129.054621 | C6H9O3+ | 0.00119237 | 0.00136337 | 0.00092674 | 0.00141226 | 0.0010314 | 0.00127379 | 0.00085157 | 0.00169489 |
| 184 | 129.09155 | C7H13O2+ | 1.78E-07 | 5.82E-07 | 1.26E-07 | 4.67E-07 | 5.92E-06 | 1.70E-05 | 6.49E-05 | 6.77E-06 |
| 185 | 129.12794 | C8H17O+ | 8.71E-06 | 2.94E-08 | 5.79E-06 | 5.04E-07 | 0.00030754 | 0.00010642 | 0.00035314 | 0.00041979 |
| 186 | 130.056669 | <unknown0110> | 2.43E-06 | 2.98E-06 | 1.87E-06 | 2.67E-06 | 7.27E-07 | 1.59E-06 | 6.08E-07 | 1.49E-06 |
| 187 | 130.088542 | <unknown0111> | 2.24E-07 | 7.11E-07 | 1.64E-07 | 5.66E-07 | 4.30E-07 | 1.36E-06 | 3.31E-07 | 1.28E-06 |
| 188 | 130.129847 | <unknown0112> | 3.25E-07 | 9.43E-07 | 2.47E-07 | 7.86E-07 | 4.19E-07 | 1.34E-06 | 3.22E-07 | 1.30E-06 |
| 189 | 131.03444 | C5H7O4+ | 1.78E-07 | 5.82E-07 | 1.26E-07 | 4.67E-07 | 3.31E-07 | 1.05E-06 | 2.47E-07 | 9.84E-07 |
| 190 | 131.04968 | C9H7O+ | 0.00205606 | 0.00059724 | 0.00141318 | 0.00072505 | 0.00177944 | 0.00045382 | 0.00137717 | 0.00067451 |
| 191 | 131.08607 | C10H11+ | 8.27E-05 | 0.0027264 | 0.00014897 | 0.00261824 | 0.00077697 | 0.00144927 | 0.0006885 | 0.00163599 |
| 192 | 131.10721 | C7H15O2+ | 1.78E-07 | 5.82E-07 | 1.26E-07 | 4.67E-07 | 3.14E-07 | 1.05E-06 | 2.33E-07 | 9.90E-07 |
| 193 | 132.10479 | <unknown0116> | 1.78E-07 | 5.82E-07 | 1.26E-07 | 4.67E-07 | 3.14E-07 | 1.05E-06 | 2.33E-07 | 9.90E-07 |
| 194 | 133.05008 | C5H9O4+ | 0.00034227 | 0.00020655 | 0.00024786 | 0.00017057 | 0.00059148 | 0.00032358 | 0.00043771 | 0.00032594 |
| 195 | 133.06534 | C9H9O+ | 1.86E-07 | 6.07E-07 | 1.30E-07 | 4.84E-07 | 3.35E-07 | 1.12E-06 | 2.50E-07 | 1.05E-06 |
| 196 | 133.08647 | C6H13O3+ | 1.99E-06 | 3.71E-06 | 1.39E-06 | 2.50E-06 | 8.83E-05 | 0 | 7.12E-05 | 5.51E-06 |
| 197 | 133.10173 | C10H13+ | 4.77E-07 | 2.13E-06 | 3.27E-07 | 1.98E-06 | 6.82E-07 | 6.53E-06 | 5.46E-07 | 6.15E-06 |
| 198 | 133.122306 | C7H17O2+ | 3.49E-07 | 6.73E-07 | 2.92E-07 | 5.84E-07 | 6.55E-07 | 1.51E-06 | 6.60E-07 | 1.53E-06 |
| 199 | 134.08117 | C5H12NO3+ | 2.14E-06 | 3.66E-06 | 1.42E-06 | 3.04E-06 | 5.37E-07 | 1.70E-06 | 4.40E-07 | 1.64E-06 |
| 200 | 135.02347 | C11H3+ | 1.93E-07 | 6.02E-07 | 1.35E-07 | 4.81E-07 | 3.66E-07 | 1.25E-06 | 2.73E-07 | 1.15E-06 |

**Table S1.** Continued.

| | | | 2024 | | | | 2025 | | | |
|---|---|---|---|---|---|---|---|---|---|---|
| | | | Forest-interior (FI) atmospheres | | Canopy-top (CT) atmospheres | | Forest-interior (FI) atmospheres | | Canopy-top (CT) atmospheres | |
| ID | Exact m/z | Ion formula | Daytime-FI | Nighttime-FI | Daytime-CT | Nighttime-CT | Daytime-FI | Nighttime-FI | Daytime-CT | Nighttime-CT |
| 201 | 135.02934 | C4H7O5+ | 4.83E-07 | 2.06E-06 | 4.81E-07 | 1.90E-06 | 6.65E-07 | 1.96E-06 | 5.18E-07 | 2.04E-06 |
| 202 | 135.0446 | C8H7O2+ | 2.17E-07 | 8.40E-07 | 1.65E-07 | 7.02E-07 | 4.35E-07 | 1.39E-06 | 3.35E-07 | 1.31E-06 |
| 203 | 135.06573 | C5H11O4+ | 2.51E-07 | 8.05E-07 | 1.70E-07 | 6.36E-07 | 7.79E-07 | 1.61E-06 | 4.48E-07 | 1.52E-06 |
| 204 | 135.08099 | C9H11O+ | 2.46E-06 | 3.23E-07 | 1.64E-06 | 3.52E-07 | 2.00E-06 | 7.93E-07 | 2.30E-06 | 7.50E-07 |
| 205 | 135.11737 | C10H15+ | 0.00179847 | 0.0093996 | 0.00112208 | 0.00758127 | 0.00183476 | 0.01562146 | 0.00149573 | 0.01471642 |
| 206 | 136.024049 | C3H6NO5+ | 0.00041008 | 0.00038816 | 0.00010122 | 9.37E-05 | 9.10E-07 | 1.97E-06 | 6.77E-07 | 1.88E-06 |
| 207 | 136.060434 | C4H10NO4+ | 1.98E-07 | 7.25E-07 | 1.49E-07 | 6.61E-07 | 4.25E-07 | 1.32E-06 | 3.19E-07 | 1.22E-06 |
| 208 | 136.07569 | C8H10NO+ | 1.27E-06 | 6.25E-07 | 1.08E-06 | 5.34E-07 | 1.76E-06 | 1.19E-06 | 1.14E-06 | 1.16E-06 |
| 209 | 136.124652 | C10H16+ | 1.79E-07 | 6.58E-06 | 4.19E-08 | 4.96E-06 | 5.71E-07 | 1.37E-05 | 3.38E-07 | 1.25E-05 |
| 210 | 137.00861 | C3H5O6+ | 1.78E-07 | 5.83E-07 | 1.27E-07 | 4.68E-07 | 3.53E-07 | 1.07E-06 | 2.68E-07 | 1.00E-06 |
| 211 | 137.02386 | C7H5O3+ | 1.78E-07 | 5.92E-07 | 1.26E-07 | 4.76E-07 | 3.47E-07 | 1.17E-06 | 2.62E-07 | 1.11E-06 |
| 212 | 137.03912 | C11H5+ | 1.56E-07 | 1.01E-06 | 1.05E-07 | 7.99E-07 | 3.83E-07 | 1.55E-06 | 2.83E-07 | 1.42E-06 |
| 213 | 137.045 | C4H9O5+ | 3.67E-07 | 5.09E-07 | 2.33E-07 | 4.37E-07 | 4.70E-07 | 1.03E-06 | 3.57E-07 | 9.84E-07 |
| 214 | 137.06026 | C8H9O2+ | 4.25E-07 | 5.36E-07 | 2.41E-07 | 4.59E-07 | 1.39E-06 | 7.83E-07 | 1.07E-06 | 7.86E-07 |
| 215 | 137.09663 | C9H13O+ | 1.79E-07 | 5.82E-07 | 1.29E-07 | 4.66E-07 | 3.18E-07 | 1.05E-06 | 2.36E-07 | 9.89E-07 |
| 216 | 137.13303 | C10H17+ | 0.0091927 | 0.08702728 | 0.0065604 | 0.06786627 | 0.01110891 | 0.15520703 | 0.0088023 | 0.14524362 |
| 217 | 138.029803 | C5H4N3O2+ | 7.16E-07 | 1.00E-06 | 4.44E-07 | 7.84E-07 | 4.00E-06 | 7.87E-06 | 2.74E-06 | 7.85E-06 |
| 218 | 138.054955 | C7H8NO2+ | 3.68E-07 | 7.13E-07 | 2.61E-07 | 5.77E-07 | 4.55E-07 | 1.29E-06 | 3.87E-07 | 1.30E-06 |
| 219 | 138.087581 | <unknown0128> | 2.37E-07 | 5.70E-07 | 1.67E-07 | 4.64E-07 | 4.35E-07 | 1.03E-06 | 3.23E-07 | 9.82E-07 |
| 220 | 139.02426 | C3H7O6+ | 1.85E-07 | 5.87E-07 | 1.30E-07 | 4.70E-07 | 3.37E-07 | 1.18E-06 | 2.51E-07 | 1.11E-06 |
| 221 | 139.03952 | C7H7O3+ | 0 | 2.84E-05 | 0.00015674 | 0.00032041 | 0.00040635 | 0.00077164 | 0.00037779 | 0.0009776 |
| 222 | 139.05478 | C11H7+ | 2.48E-07 | 9.87E-07 | 1.81E-07 | 8.54E-07 | 3.45E-07 | 1.11E-06 | 2.56E-07 | 1.04E-06 |
| 223 | 139.0759 | C8H11O2+ | 1.19E-06 | 1.25E-06 | 4.70E-07 | 8.25E-07 | 4.39E-07 | 1.19E-06 | 3.75E-07 | 1.10E-06 |
| 224 | 139.11229 | C9H15O+ | 0.0032364 | 0.00388782 | 0.00247082 | 0.00347839 | 0.00439052 | 0.00597294 | 0.00334591 | 0.00593764 |
| 225 | 139.14868 | C10H19+ | 1.83E-07 | 6.01E-07 | 1.26E-07 | 4.83E-07 | 3.22E-07 | 1.24E-06 | 2.51E-07 | 1.18E-06 |
| 226 | 140.006082 | <unknown0129> | 1.56E-06 | 3.61E-06 | 1.46E-06 | 2.61E-06 | 4.55E-07 | 1.66E-06 | 3.42E-07 | 1.53E-06 |
| 227 | 140.072028 | <unknown0131> | 5.18E-07 | 1.41E-06 | 5.01E-07 | 1.02E-06 | 8.10E-07 | 1.60E-06 | 6.48E-07 | 1.45E-06 |
| 228 | 140.151385 | <unknown0133> | 2.11E-07 | 6.94E-07 | 1.50E-07 | 5.54E-07 | 3.83E-07 | 1.36E-06 | 2.99E-07 | 1.28E-06 |
| 229 | 141.05518 | C7H9O3+ | 0.00029971 | 0.00029055 | 0.00023314 | 0.00024991 | 0.00035444 | 0.00020195 | 0.00027721 | 0.00021939 |
| 230 | 141.07042 | C11H9+ | 1.78E-07 | 5.82E-07 | 1.26E-07 | 4.67E-07 | 3.15E-07 | 1.06E-06 | 2.34E-07 | 9.91E-07 |
| 231 | 141.09155 | C8H13O2+ | 4.98E-07 | 6.57E-07 | 4.62E-07 | 5.20E-07 | 0.00122794 | 0 | 0.00087044 | 0 |
| 232 | 141.12794 | C9H17O+ | 0.00082408 | 0.00066637 | 0.00058819 | 0.00071285 | 0.00019758 | 0.00063452 | 0.00105984 | 0.00099737 |
| 233 | 142.086255 | C7H12NO2+ | 7.87E-06 | 9.80E-06 | 2.40E-06 | 1.48E-05 | 7.21E-07 | 1.99E-06 | 5.82E-07 | 1.93E-06 |
| 234 | 143.01331 | C9H3O2+ | 0.00030965 | 0.00033606 | 0.00022477 | 0.00027881 | 0.00010439 | 0.00055716 | 8.82E-05 | 0.00065973 |
| 235 | 143.04968 | C10H7O+ | 0.0001453 | 0.00111682 | 0.00013332 | 0.00101188 | 0.00049386 | 0.00074212 | 0.00043082 | 0.0008837 |
| 236 | 143.07082 | C7H11O3+ | 1.78E-07 | 5.82E-07 | 1.26E-07 | 4.67E-07 | 3.14E-07 | 1.05E-06 | 2.33E-07 | 9.90E-07 |
| 237 | 143.08607 | C11H11+ | 1.78E-07 | 5.82E-07 | 1.26E-07 | 4.67E-07 | 3.57E-07 | 1.05E-06 | 2.71E-07 | 9.87E-07 |
| 238 | 143.10721 | C8H15O2+ | 6.25E-06 | 7.56E-07 | 4.94E-06 | 6.78E-07 | 0.0018118 | 0.00188595 | 0.00139285 | 0.0015861 |
| 239 | 143.14359 | C9H19O+ | 0.00191761 | 0.00140528 | 0.00116554 | 0.00115892 | 0.00018671 | 0 | 0.00169148 | 0.00033669 |
| 240 | 144.107365 | <unknown0140> | 4.23E-07 | 1.03E-06 | 3.26E-07 | 8.39E-07 | 4.67E-07 | 1.40E-06 | 3.57E-07 | 1.34E-06 |
| 241 | 145.0137 | C5H5O5+ | 0.00010703 | 0.00011765 | 8.22E-05 | 0.00011857 | 8.09E-05 | 0.00020415 | 6.73E-05 | 0.00022116 |
| 242 | 145.06534 | C10H9O+ | 0.00038998 | 0.00083443 | 0.00033187 | 0.00076424 | 0.00051946 | 0.00057009 | 0.00041908 | 0.00058453 |
| 243 | 145.08647 | C7H13O3+ | 1.78E-07 | 5.82E-07 | 1.26E-07 | 4.67E-07 | 3.29E-07 | 1.06E-06 | 2.48E-07 | 9.99E-07 |
| 244 | 145.10173 | C11H13+ | 1.79E-07 | 5.82E-07 | 1.26E-07 | 4.68E-07 | 5.45E-07 | 1.38E-06 | 4.43E-07 | 1.30E-06 |
| 245 | 145.12285 | C8H17O2+ | 1.19E-06 | 9.58E-07 | 1.23E-06 | 8.53E-07 | 1.94E-06 | 6.11E-06 | 2.30E-06 | 6.04E-06 |
| 246 | 145.969167 | <unknown0141> | 2.45E-07 | 7.30E-07 | 1.78E-07 | 5.91E-07 | 4.06E-07 | 1.42E-06 | 2.99E-07 | 1.32E-06 |
| 247 | 146.014329 | <unknown0142> | 1.03E-06 | 2.01E-06 | 7.23E-07 | 1.63E-06 | 4.87E-07 | 1.47E-06 | 3.84E-07 | 1.39E-06 |
| 248 | 146.125175 | <unknown0145> | 4.77E-07 | 1.19E-06 | 3.73E-07 | 9.62E-07 | 4.04E-07 | 1.33E-06 | 3.11E-07 | 1.25E-06 |
| 249 | 146.977158 | H3O9+ | 2.36E-07 | 6.64E-07 | 1.70E-07 | 5.48E-07 | 4.30E-07 | 1.58E-06 | 3.19E-07 | 1.46E-06 |
| 250 | 146.99297 | C4H3O6+ | 4.31E-06 | 1.59E-05 | 2.37E-06 | 9.13E-06 | 2.85E-06 | 7.86E-06 | 2.22E-06 | 7.81E-06 |

**Table S1.** Continued.

| | | | 2024 | | | | 2025 | | | |
|---|---|---|---|---|---|---|---|---|---|---|
| | | | Forest-interior (FI) atmospheres | | Canopy-top (CT) atmospheres | | Forest-interior (FI) atmospheres | | Canopy-top (CT) atmospheres | |
| ID | Exact m/z | Ion formula | Daytime-FI | Nighttime-FI | Daytime-CT | Nighttime-CT | Daytime-FI | Nighttime-FI | Daytime-CT | Nighttime-CT |
| 251 | 147.06573 | C6H11O4+ | 0.00039387 | 0.00047338 | 0.00030159 | 0.00041962 | 0.00040136 | 0.0002756 | 0.00028316 | 0.00023763 |
| 252 | 147.08099 | C10H11O+ | 1.81E-07 | 5.97E-07 | 1.28E-07 | 4.81E-07 | 3.34E-07 | 1.13E-06 | 2.51E-07 | 1.06E-06 |
| 253 | 147.10211 | C7H15O3+ | 1.85E-07 | 6.46E-07 | 1.31E-07 | 5.19E-07 | 4.50E-07 | 2.59E-06 | 3.48E-07 | 2.57E-06 |
| 254 | 147.11737 | C11H15+ | 3.61E-07 | 2.15E-06 | 2.69E-07 | 1.75E-06 | 7.21E-07 | 3.51E-06 | 6.00E-07 | 3.10E-06 |
| 255 | 147.137956 | C8H19O2+ | 0.0005255 | 0 | 0.00047479 | 0 | 1.40E-06 | 2.06E-06 | 1.44E-06 | 2.21E-06 |
| 256 | 147.972792 | <unknown0147> | 1.99E-06 | 2.81E-06 | 9.77E-07 | 2.35E-06 | 4.99E-07 | 1.72E-06 | 3.75E-07 | 1.62E-06 |
| 257 | 148.060434 | C5H10NO4+ | 9.43E-05 | 9.18E-05 | 7.67E-05 | 8.09E-05 | 2.71E-06 | 1.80E-06 | 2.10E-06 | 1.82E-06 |
| 258 | 148.971679 | C3HO7+ | 1.05E-06 | 1.44E-06 | 6.50E-07 | 1.30E-06 | 7.03E-07 | 2.61E-06 | 4.57E-07 | 2.33E-06 |
| 259 | 149.00861 | C4H5O6+ | 2.03E-07 | 6.85E-07 | 1.46E-07 | 5.49E-07 | 4.42E-07 | 1.67E-06 | 3.24E-07 | 1.59E-06 |
| 260 | 149.02386 | C8H5O3+ | 2.80E-07 | 8.05E-07 | 2.42E-07 | 7.18E-07 | 6.00E-07 | 1.61E-06 | 6.83E-07 | 1.51E-06 |
| 261 | 149.03912 | C12H5+ | 2.35E-07 | 6.84E-07 | 1.69E-07 | 5.61E-07 | 4.05E-07 | 1.24E-06 | 3.02E-07 | 1.17E-06 |
| 262 | 149.045 | C5H9O5+ | 2.33E-07 | 7.29E-07 | 1.62E-07 | 5.83E-07 | 4.81E-07 | 1.43E-06 | 3.45E-07 | 1.34E-06 |
| 263 | 149.06026 | C9H9O2+ | 2.13E-07 | 7.63E-07 | 1.50E-07 | 6.07E-07 | 4.31E-07 | 1.42E-06 | 3.27E-07 | 1.35E-06 |
| 264 | 149.08139 | C6H13O4+ | 3.51E-07 | 1.07E-06 | 2.49E-07 | 8.32E-07 | 6.38E-07 | 1.83E-06 | 4.40E-07 | 1.65E-06 |
| 265 | 149.09663 | C10H13O+ | 4.29E-07 | 7.95E-07 | 3.07E-07 | 6.69E-07 | 1.40E-06 | 2.54E-06 | 1.26E-06 | 2.22E-06 |
| 266 | 149.13303 | C11H17+ | 0.0001909 | 0.00245844 | 0.00014791 | 0.00191514 | 0.00026618 | 0.00263443 | 0.00034994 | 0.00232858 |
| 267 | 149.974761 | <unknown0152> | 2.46E-07 | 7.40E-07 | 1.67E-07 | 5.83E-07 | 3.99E-07 | 1.36E-06 | 3.00E-07 | 1.27E-06 |
| 268 | 150.050932 | C3H8N3O4+ | 4.18E-06 | 6.06E-06 | 9.89E-06 | 1.23E-05 | 5.23E-07 | 1.48E-06 | 3.91E-07 | 1.41E-06 |
| 269 | 150.09134 | C9H12NO+ | 2.86E-07 | 7.68E-07 | 1.98E-07 | 6.09E-07 | 5.30E-07 | 1.64E-06 | 4.00E-07 | 1.55E-06 |
| 270 | 150.134307 | <unknown0155> | 2.83E-07 | 8.45E-07 | 2.03E-07 | 6.73E-07 | 4.49E-07 | 1.56E-06 | 3.43E-07 | 1.47E-06 |
| 271 | 151.00313 | C7H3O4+ | 4.74E-07 | 9.50E-07 | 3.44E-07 | 8.08E-07 | 6.88E-07 | 2.51E-06 | 5.26E-07 | 2.59E-06 |
| 272 | 151.03952 | C8H7O3+ | 8.67E-05 | 0.00023592 | 6.57E-05 | 0.00019947 | 7.55E-05 | 0.00016326 | 5.01E-05 | 0.00020171 |
| 273 | 151.06065 | C5H11O5+ | 1.88E-07 | 5.96E-07 | 1.33E-07 | 4.79E-07 | 4.36E-07 | 1.19E-06 | 3.20E-07 | 1.12E-06 |
| 274 | 151.0759 | C9H11O2+ | 2.12E-07 | 5.96E-07 | 1.49E-07 | 4.80E-07 | 4.51E-07 | 1.14E-06 | 3.42E-07 | 1.09E-06 |
| 275 | 151.11229 | C10H15O+ | 0.00130181 | 0.00298548 | 0.00099152 | 0.00242413 | 0.00170919 | 0.00596809 | 0.00134673 | 0.00523088 |
| 276 | 151.14868 | C11H19+ | 3.74E-07 | 8.32E-07 | 2.87E-07 | 6.90E-07 | 8.42E-07 | 1.53E-06 | 1.14E-06 | 1.64E-06 |
| 277 | 153.01878 | C7H5O4+ | 1.82E-07 | 5.95E-07 | 1.30E-07 | 4.81E-07 | 3.19E-07 | 1.17E-06 | 2.39E-07 | 1.10E-06 |
| 278 | 153.03404 | C11H5O+ | 1.74E-07 | 7.58E-07 | 1.23E-07 | 6.05E-07 | 3.21E-07 | 1.12E-06 | 2.38E-07 | 1.05E-06 |
| 279 | 153.03992 | C4H9O6+ | 1.90E-07 | 6.65E-07 | 1.35E-07 | 5.45E-07 | 3.73E-07 | 1.32E-06 | 2.82E-07 | 1.25E-06 |
| 280 | 153.05518 | C8H9O3+ | 3.08E-06 | 9.87E-07 | 1.94E-06 | 9.09E-07 | 0.00128563 | 0 | 0.00093345 | 0 |
| 281 | 153.07042 | C12H9+ | 2.08E-07 | 6.34E-07 | 1.50E-07 | 5.05E-07 | 4.06E-07 | 1.16E-06 | 3.06E-07 | 1.07E-06 |
| 282 | 153.09155 | C9H13O2+ | 2.28E-07 | 5.71E-07 | 1.65E-07 | 4.63E-07 | 1.01E-06 | 9.33E-07 | 7.95E-07 | 8.95E-07 |
| 283 | 153.12794 | C10H17O+ | 0.00159488 | 0.01159945 | 0.00104744 | 0.00923555 | 0.00175401 | 0.02210786 | 0.00140584 | 0.02009971 |
| 284 | 153.16432 | C11H21+ | 1.78E-07 | 5.82E-07 | 1.26E-07 | 4.67E-07 | 3.40E-07 | 1.06E-06 | 2.63E-07 | 9.99E-07 |
| 285 | 155.01331 | C10H3O2+ | 2.59E-07 | 7.23E-07 | 1.89E-07 | 5.93E-07 | 4.05E-07 | 1.30E-06 | 2.92E-07 | 1.22E-06 |
| 286 | 155.03444 | C7H7O4+ | 2.82E-07 | 9.15E-07 | 2.18E-07 | 7.66E-07 | 1.26E-06 | 6.22E-06 | 1.06E-06 | 7.28E-06 |
| 287 | 155.04968 | C11H7O+ | 2.49E-07 | 7.90E-07 | 1.75E-07 | 6.38E-07 | 4.08E-07 | 1.30E-06 | 3.11E-07 | 1.20E-06 |
| 288 | 155.07082 | C8H11O3+ | 2.49E-07 | 6.81E-07 | 1.63E-07 | 5.39E-07 | 7.74E-07 | 1.43E-06 | 5.80E-07 | 1.31E-06 |
| 289 | 155.08607 | C12H11+ | 2.93E-07 | 9.24E-07 | 1.91E-07 | 7.09E-07 | 4.51E-07 | 1.53E-06 | 3.30E-07 | 1.48E-06 |
| 290 | 155.10721 | C9H15O2+ | 4.45E-06 | 2.58E-06 | 4.60E-06 | 2.22E-06 | 0.00078671 | 3.87E-05 | 0.00065752 | 2.56E-05 |
| 291 | 155.14359 | C10H19O+ | 1.76E-06 | 4.84E-06 | 7.31E-07 | 5.08E-06 | 8.30E-07 | 7.93E-06 | 7.66E-07 | 8.02E-06 |
| 292 | 155.17998 | C11H23+ | 2.19E-07 | 6.42E-07 | 1.69E-07 | 5.15E-07 | 4.52E-07 | 1.28E-06 | 4.55E-07 | 1.29E-06 |
| 293 | 156.106309 | <unknown0165> | 5.03E-07 | 1.35E-06 | 3.94E-07 | 1.08E-06 | 4.95E-07 | 1.45E-06 | 3.68E-07 | 1.37E-06 |
| 294 | 156.146055 | <unknown0166> | 2.93E-07 | 7.75E-07 | 2.08E-07 | 6.32E-07 | 3.94E-07 | 1.31E-06 | 3.00E-07 | 1.23E-06 |
| 295 | 157.02896 | C10H5O2+ | 1.28E-06 | 9.36E-07 | 6.87E-07 | 8.17E-07 | 2.17E-05 | 0.00031148 | 9.30E-05 | 0.00037185 |
| 296 | 157.05008 | C7H9O4+ | 0.00014666 | 0.00030283 | 7.63E-05 | 0.00054828 | 3.61E-06 | 8.83E-06 | 2.21E-06 | 9.03E-06 |
| 297 | 157.06534 | C11H9O+ | 1.87E-07 | 6.36E-07 | 1.34E-07 | 5.06E-07 | 4.55E-07 | 1.42E-06 | 3.46E-07 | 1.32E-06 |
| 298 | 157.08647 | C8H13O3+ | 1.79E-07 | 5.83E-07 | 1.27E-07 | 4.68E-07 | 5.77E-07 | 1.25E-06 | 4.47E-07 | 1.16E-06 |
| 299 | 157.10173 | C12H13+ | 2.13E-07 | 6.08E-07 | 1.52E-07 | 4.83E-07 | 5.50E-07 | 1.57E-06 | 4.05E-07 | 1.46E-06 |
| 300 | 157.12285 | C9H17O2+ | 0.00012312 | 0.00092765 | 3.74E-06 | 0.00089185 | 0.00070772 | 0.0010781 | 0.00068826 | 0.00096837 |

**Table S1.** Continued.

| | | | 2024 | | | | 2025 | | | |
|---|---|---|---|---|---|---|---|---|---|---|
| | | | Forest-interior (FI) atmospheres | | Canopy-top (CT) atmospheres | | Forest-interior (FI) atmospheres | | Canopy-top (CT) atmospheres | |
| ID | Exact m/z | Ion formula | Daytime-FI | Nighttime-FI | Daytime-CT | Nighttime-CT | Daytime-FI | Nighttime-FI | Daytime-CT | Nighttime-CT |
| 301 | 157.15924 | C10H21O+ | 0.00162837 | 0.00042225 | 0.00114152 | 0.00042749 | 6.22E-07 | 1.31E-06 | 1.04E-06 | 1.21E-06 |
| 302 | 158.08117 | C7H12NO3+ | 4.80E-07 | 1.10E-06 | 3.03E-07 | 8.78E-07 | 4.95E-07 | 1.39E-06 | 3.85E-07 | 1.30E-06 |
| 303 | 158.117555 | C8H16NO2+ | 2.83E-07 | 7.92E-07 | 2.09E-07 | 6.51E-07 | 4.10E-07 | 1.33E-06 | 3.20E-07 | 1.24E-06 |
| 304 | 159.0446 | C10H7O2+ | 0.00013317 | 0.00036067 | 0.00011111 | 0.00031935 | 0.00022361 | 0.00040275 | 0.00018253 | 0.00045192 |
| 305 | 159.08099 | C11H11O+ | 1.80E-07 | 5.84E-07 | 1.27E-07 | 4.69E-07 | 5.11E-07 | 1.17E-06 | 3.73E-07 | 1.10E-06 |
| 306 | 159.10211 | C8H15O3+ | 5.16E-07 | 8.74E-07 | 3.25E-07 | 6.55E-07 | 1.35E-05 | 7.61E-06 | 6.45E-06 | 6.48E-06 |
| 307 | 159.11737 | C12H15+ | 2.84E-07 | 1.21E-06 | 2.07E-07 | 9.65E-07 | 4.23E-07 | 1.60E-06 | 3.27E-07 | 1.53E-06 |
| 308 | 159.1385 | C9H19O2+ | 3.42E-07 | 9.04E-07 | 3.59E-07 | 1.07E-06 | 5.33E-07 | 1.75E-06 | 2.14E-06 | 1.86E-06 |
| 309 | 160.133205 | C8H18NO2+ | 2.78E-07 | 8.26E-07 | 2.29E-07 | 8.14E-07 | 4.15E-07 | 1.37E-06 | 3.50E-07 | 1.33E-06 |
| 310 | 160.169591 | C9H22NO+ | 5.03E-07 | 7.87E-07 | 2.85E-07 | 6.16E-07 | 3.57E-07 | 1.18E-06 | 2.78E-07 | 1.12E-06 |
| 311 | 161.00861 | C5H5O6+ | 5.88E-05 | 9.43E-05 | 4.45E-05 | 8.53E-05 | 5.02E-05 | 0.00014166 | 3.63E-05 | 0.00014821 |
| 312 | 161.08139 | C7H13O4+ | 2.02E-05 | 1.81E-05 | 2.30E-05 | 9.04E-06 | 4.69E-05 | 4.45E-06 | 3.42E-05 | 4.53E-06 |
| 313 | 161.09663 | C11H13O+ | 1.78E-07 | 5.83E-07 | 1.26E-07 | 4.68E-07 | 3.18E-07 | 1.06E-06 | 2.37E-07 | 1.00E-06 |
| 314 | 161.11777 | C8H17O3+ | 1.78E-07 | 5.93E-07 | 1.26E-07 | 4.74E-07 | 3.80E-07 | 1.34E-06 | 2.84E-07 | 1.27E-06 |
| 315 | 161.13303 | C12H17+ | 3.18E-07 | 1.25E-06 | 3.08E-07 | 1.67E-06 | 2.66E-06 | 4.94E-06 | 1.66E-06 | 4.96E-06 |
| 316 | 161.153606 | C9H21O2+ | 0.00191848 | 0.00103873 | 0.00131124 | 0.00020106 | 1.80E-06 | 2.19E-06 | 0.00033728 | 0 |
| 317 | 163.01839 | C12H3O+ | 1.80E-07 | 5.89E-07 | 1.28E-07 | 4.73E-07 | 3.34E-07 | 1.16E-06 | 2.47E-07 | 1.09E-06 |
| 318 | 163.02426 | C5H7O6+ | 1.43E-05 | 6.42E-05 | 2.53E-05 | 5.45E-05 | 2.17E-06 | 3.41E-06 | 2.05E-06 | 3.74E-06 |
| 319 | 163.14868 | C12H19+ | 0.00025979 | 0.00089727 | 0.00017477 | 0.00071565 | 9.57E-05 | 0.00052993 | 0.00011561 | 0.00044688 |
| 320 | 164.034219 | C8H6NO3+ | 2.26E-06 | 2.16E-06 | 7.81E-07 | 1.31E-06 | 4.90E-07 | 1.36E-06 | 3.94E-07 | 1.34E-06 |
| 321 | 165.01878 | C8H5O4+ | 1.81E-07 | 5.88E-07 | 1.29E-07 | 4.73E-07 | 3.53E-07 | 1.18E-06 | 2.61E-07 | 1.10E-06 |
| 322 | 165.03404 | C12H5O+ | 8.96E-05 | 0.0001222 | 6.75E-05 | 0.00011415 | 1.32E-05 | 1.38E-05 | 7.20E-06 | 1.41E-05 |
| 323 | 165.12794 | C11H17O+ | 7.46E-05 | 0.00081545 | 4.73E-05 | 0.00062182 | 8.66E-05 | 0.00093306 | 7.43E-05 | 0.00076572 |
| 324 | 165.16432 | C12H21+ | 2.99E-07 | 6.60E-07 | 2.31E-07 | 5.65E-07 | 4.18E-07 | 1.49E-06 | 3.74E-07 | 1.50E-06 |
| 325 | 166.166343 | <unknown0189> | 2.31E-07 | 7.35E-07 | 1.71E-07 | 5.96E-07 | 3.78E-07 | 1.31E-06 | 2.94E-07 | 1.24E-06 |
| 326 | 167.01331 | C11H3O2+ | 1.91E-07 | 6.15E-07 | 1.38E-07 | 4.91E-07 | 3.52E-07 | 1.18E-06 | 2.55E-07 | 1.11E-06 |
| 327 | 167.01918 | C4H7O7+ | 2.03E-07 | 6.44E-07 | 1.49E-07 | 5.14E-07 | 4.06E-07 | 1.36E-06 | 3.03E-07 | 1.29E-06 |
| 328 | 167.03444 | C8H7O4+ | 1.60E-06 | 1.75E-06 | 3.77E-06 | 3.40E-06 | 8.65E-07 | 2.32E-06 | 1.09E-06 | 2.17E-06 |
| 329 | 167.05556 | C5H11O6+ | 2.24E-07 | 7.53E-07 | 1.41E-07 | 5.81E-07 | 5.03E-07 | 1.35E-06 | 3.56E-07 | 1.26E-06 |
| 330 | 167.08607 | C13H11+ | 2.33E-07 | 6.84E-07 | 1.55E-07 | 5.36E-07 | 4.91E-07 | 1.41E-06 | 3.64E-07 | 1.33E-06 |
| 331 | 167.10721 | C10H15O2+ | 1.55E-05 | 1.71E-05 | 2.62E-05 | 1.57E-05 | 0.00040871 | 0.00129087 | 0.00033464 | 0.00070782 |
| 332 | 167.14359 | C11H19O+ | 2.70E-07 | 9.29E-07 | 2.09E-07 | 7.74E-07 | 6.15E-07 | 2.08E-06 | 5.78E-07 | 2.05E-06 |
| 333 | 167.17998 | C12H23+ | 2.39E-07 | 6.88E-07 | 1.76E-07 | 5.59E-07 | 3.97E-07 | 1.32E-06 | 3.85E-07 | 1.34E-06 |
| 334 | 168.050263 | C4H10NO6+ | 1.20E-06 | 2.03E-06 | 8.40E-07 | 2.15E-06 | 5.68E-07 | 1.65E-06 | 4.16E-07 | 1.57E-06 |
| 335 | 169.0137 | C7H5O5+ | 1.97E-07 | 6.27E-07 | 1.42E-07 | 5.08E-07 | 3.91E-07 | 1.34E-06 | 2.99E-07 | 1.27E-06 |
| 336 | 169.02896 | C11H5O2+ | 1.91E-07 | 6.11E-07 | 1.36E-07 | 4.95E-07 | 3.42E-07 | 1.16E-06 | 2.53E-07 | 1.09E-06 |
| 337 | 169.03482 | C4H9O7+ | 5.15E-07 | 1.53E-06 | 3.59E-07 | 1.20E-06 | 6.48E-07 | 1.82E-06 | 4.95E-07 | 1.69E-06 |
| 338 | 169.08647 | C9H13O3+ | 2.44E-07 | 6.84E-07 | 1.68E-07 | 5.41E-07 | 8.73E-07 | 1.35E-06 | 6.16E-07 | 1.28E-06 |
| 339 | 169.10173 | C13H13+ | 2.40E-07 | 6.46E-07 | 1.65E-07 | 5.19E-07 | 4.04E-07 | 1.38E-06 | 2.97E-07 | 1.29E-06 |
| 340 | 169.12285 | C10H17O2+ | 0.0009298 | 0.00208469 | 0.00072914 | 0.00166151 | 0.0014542 | 0.00338431 | 0.00118375 | 0.0027972 |
| 341 | 169.15924 | C11H21O+ | 1.85E-07 | 6.29E-07 | 1.30E-07 | 5.14E-07 | 3.57E-07 | 1.19E-06 | 2.75E-07 | 1.14E-06 |
| 342 | 169.19563 | C12H25+ | 2.03E-07 | 5.95E-07 | 1.54E-07 | 4.84E-07 | 3.66E-07 | 1.16E-06 | 3.69E-07 | 1.17E-06 |
| 343 | 171.02347 | C14H3+ | 2.03E-07 | 6.15E-07 | 1.43E-07 | 4.96E-07 | 3.83E-07 | 1.25E-06 | 2.84E-07 | 1.18E-06 |
| 344 | 171.02934 | C7H7O5+ | 2.77E-07 | 7.19E-07 | 2.09E-07 | 6.00E-07 | 3.95E-07 | 1.40E-06 | 2.96E-07 | 1.35E-06 |
| 345 | 171.0446 | C11H7O2+ | 2.19E-07 | 1.15E-06 | 1.49E-07 | 8.89E-07 | 5.56E-07 | 1.47E-06 | 4.12E-07 | 1.34E-06 |
| 346 | 171.08099 | C12H11O+ | 7.72E-07 | 7.75E-07 | 4.78E-07 | 6.57E-07 | 3.05E-06 | 7.68E-06 | 2.29E-06 | 7.75E-06 |
| 347 | 171.10211 | C9H15O3+ | 2.36E-07 | 6.00E-07 | 1.79E-07 | 4.86E-07 | 7.76E-07 | 1.16E-06 | 6.13E-07 | 1.14E-06 |
| 348 | 171.11737 | C13H15+ | 2.26E-07 | 6.22E-07 | 1.59E-07 | 4.99E-07 | 4.59E-07 | 1.37E-06 | 3.49E-07 | 1.28E-06 |
| 349 | 171.1385 | C10H19O2+ | 6.66E-06 | 0.00463241 | 2.86E-06 | 0.00369282 | 5.91E-06 | 0.00527274 | 4.71E-06 | 0.00480784 |
| 350 | 171.1749 | C11H23O+ | 3.92E-07 | 5.42E-07 | 2.92E-07 | 4.51E-07 | 5.68E-07 | 1.02E-06 | 1.55E-06 | 8.09E-07 |

**Table S1.** Continued.

| | | | 2024 | | | | 2025 | | | |
|---|---|---|---|---|---|---|---|---|---|---|
| | | | Forest-interior (FI) atmospheres | | Canopy-top (CT) atmospheres | | Forest-interior (FI) atmospheres | | Canopy-top (CT) atmospheres | |
| ID | Exact m/z | Ion formula | Daytime-FI | Nighttime-FI | Daytime-CT | Nighttime-CT | Daytime-FI | Nighttime-FI | Daytime-CT | Nighttime-CT |
| 351 | 172.134802 | <unknown0201> | 4.26E-07 | 9.50E-07 | 3.19E-07 | 7.57E-07 | 4.22E-07 | 1.46E-06 | 3.17E-07 | 1.38E-06 |
| 352 | 173.00861 | C6H5O6+ | 1.95E-07 | 6.17E-07 | 1.38E-07 | 4.96E-07 | 3.80E-07 | 1.25E-06 | 2.86E-07 | 1.21E-06 |
| 353 | 173.02386 | C10H5O3+ | 1.68E-06 | 4.41E-06 | 1.18E-06 | 4.59E-06 | 1.03E-06 | 3.26E-06 | 7.42E-07 | 3.12E-06 |
| 354 | 173.08139 | C8H13O4+ | 6.91E-07 | 2.03E-06 | 4.41E-07 | 1.55E-06 | 2.89E-06 | 3.38E-06 | 2.98E-06 | 3.38E-06 |
| 355 | 173.09663 | C12H13O+ | 2.03E-07 | 6.17E-07 | 1.45E-07 | 4.96E-07 | 3.81E-07 | 1.23E-06 | 2.82E-07 | 1.16E-06 |
| 356 | 173.11777 | C9H17O3+ | 3.50E-07 | 8.36E-07 | 2.78E-07 | 6.80E-07 | 1.71E-06 | 3.37E-06 | 9.52E-07 | 2.58E-06 |
| 357 | 173.13303 | C13H17+ | 3.39E-07 | 1.08E-06 | 2.32E-07 | 8.78E-07 | 4.88E-07 | 1.57E-06 | 3.76E-07 | 1.48E-06 |
| 358 | 173.15416 | C10H21O2+ | 2.47E-07 | 8.27E-07 | 1.80E-07 | 6.78E-07 | 4.07E-07 | 1.37E-06 | 3.53E-07 | 1.34E-06 |
| 359 | 174.148855 | C9H20NO2+ | 2.53E-07 | 7.83E-07 | 1.81E-07 | 6.34E-07 | 3.87E-07 | 1.26E-06 | 2.90E-07 | 1.17E-06 |
| 360 | 174.185241 | C10H24NO+ | 5.82E-07 | 7.73E-07 | 4.07E-07 | 6.30E-07 | 3.46E-07 | 1.15E-06 | 2.61E-07 | 1.08E-06 |
| 361 | 174.98788 | C5H3O7+ | 2.81E-07 | 7.46E-07 | 2.08E-07 | 6.13E-07 | 4.62E-07 | 1.66E-06 | 3.52E-07 | 1.62E-06 |
| 362 | 175.03952 | C10H7O3+ | 5.62E-05 | 0.00011914 | 4.56E-05 | 0.00011213 | 9.87E-05 | 0.00017058 | 7.62E-05 | 0.00017171 |
| 363 | 175.09703 | C8H15O4+ | 3.35E-07 | 1.04E-06 | 2.54E-07 | 7.79E-07 | 7.91E-06 | 1.32E-06 | 5.58E-06 | 1.50E-06 |
| 364 | 175.13342 | C9H19O3+ | 1.76E-07 | 6.43E-07 | 1.24E-07 | 5.15E-07 | 4.33E-07 | 1.61E-06 | 3.64E-07 | 1.54E-06 |
| 365 | 175.14868 | C13H19+ | 2.19E-07 | 7.41E-07 | 1.56E-07 | 6.05E-07 | 4.36E-07 | 1.70E-06 | 3.62E-07 | 1.61E-06 |
| 366 | 175.169256 | C10H23O2+ | 0.00154996 | 0.00068682 | 0.00111173 | 0.00062118 | 5.43E-07 | 1.36E-06 | 5.85E-07 | 1.28E-06 |
| 367 | 177.05518 | C10H9O3+ | 4.82E-06 | 1.05E-05 | 3.01E-05 | 1.07E-05 | 5.88E-07 | 1.73E-06 | 4.32E-07 | 1.75E-06 |
| 368 | 177.07042 | C14H9+ | 2.09E-07 | 6.55E-07 | 1.46E-07 | 5.23E-07 | 5.15E-07 | 1.31E-06 | 3.85E-07 | 1.22E-06 |
| 369 | 177.16432 | C13H21+ | 0.00044968 | 0.00055941 | 0.00035369 | 0.00044811 | 6.68E-05 | 0.00010771 | 0.00028912 | 0.00026551 |
| 370 | 179.07082 | C10H11O3+ | 8.42E-05 | 9.13E-05 | 9.93E-05 | 0.00014035 | 8.41E-06 | 2.92E-05 | 7.19E-06 | 2.04E-05 |
| 371 | 179.14359 | C12H19O+ | 4.13E-06 | 7.53E-06 | 2.54E-06 | 4.30E-06 | 0.00019869 | 3.10E-05 | 0.00017985 | 2.50E-05 |
| 372 | 179.17998 | C13H23+ | 3.00E-07 | 7.82E-07 | 2.28E-07 | 6.85E-07 | 3.71E-07 | 1.36E-06 | 3.13E-07 | 1.33E-06 |
| 373 | 181.05008 | C9H9O4+ | 4.28E-07 | 1.35E-06 | 3.45E-07 | 1.25E-06 | 6.90E-07 | 2.41E-06 | 5.06E-07 | 2.43E-06 |
| 374 | 181.06534 | C13H9O+ | 2.53E-07 | 6.87E-07 | 1.81E-07 | 5.55E-07 | 5.10E-07 | 1.32E-06 | 3.78E-07 | 1.22E-06 |
| 375 | 181.12285 | C11H17O2+ | 0 | 0.00019663 | 0 | 2.91E-05 | 3.88E-06 | 0.0007009 | 7.14E-06 | 0.00048363 |
| 376 | 181.15924 | C12H21O+ | 2.52E-07 | 8.11E-07 | 1.88E-07 | 6.95E-07 | 4.05E-07 | 1.60E-06 | 3.36E-07 | 1.56E-06 |
| 377 | 181.19563 | C13H25+ | 2.38E-07 | 6.64E-07 | 1.75E-07 | 5.37E-07 | 3.80E-07 | 1.27E-06 | 3.63E-07 | 1.30E-06 |
| 378 | 182.030865 | C5H4N5O3+ | 2.48E-07 | 7.08E-07 | 1.68E-07 | 5.63E-07 | 3.76E-07 | 1.24E-06 | 2.86E-07 | 1.16E-06 |
| 379 | 183.08099 | C13H11O+ | 0.00040689 | 0.00050623 | 0.00013942 | 0.00013076 | 2.17E-05 | 3.89E-05 | 6.81E-05 | 0.00012428 |
| 380 | 183.10211 | C10H15O3+ | 2.00E-07 | 6.23E-07 | 1.45E-07 | 4.87E-07 | 7.10E-07 | 1.34E-06 | 5.05E-07 | 1.21E-06 |
| 381 | 183.1385 | C11H19O2+ | 1.52E-07 | 2.57E-06 | 8.70E-08 | 2.61E-06 | 2.59E-07 | 4.84E-05 | 7.95E-07 | 3.42E-05 |
| 382 | 183.1749 | C12H23O+ | 8.36E-07 | 8.02E-07 | 6.28E-07 | 6.72E-07 | 3.81E-07 | 1.22E-06 | 3.15E-07 | 1.19E-06 |
| 383 | 183.21127 | C13H27+ | 2.98E-07 | 6.58E-07 | 2.26E-07 | 5.39E-07 | 3.86E-07 | 1.17E-06 | 3.96E-07 | 1.18E-06 |
| 384 | 185.02386 | C11H5O3+ | 4.29E-05 | 5.34E-05 | 4.27E-05 | 5.13E-05 | 4.47E-05 | 8.93E-05 | 2.94E-05 | 8.86E-05 |
| 385 | 185.11777 | C10H17O3+ | 1.56E-05 | 4.08E-06 | 4.89E-06 | 4.62E-06 | 0.00068894 | 0.00025515 | 0.00050458 | 0.00013388 |
| 386 | 185.13303 | C14H17+ | 2.04E-07 | 8.96E-07 | 1.46E-07 | 7.29E-07 | 3.60E-07 | 1.31E-06 | 2.75E-07 | 1.23E-06 |
| 387 | 185.15416 | C11H21O2+ | 2.03E-07 | 7.14E-07 | 1.44E-07 | 5.75E-07 | 3.62E-07 | 1.47E-06 | 2.82E-07 | 1.41E-06 |
| 388 | 185.19054 | C12H25O+ | 2.99E-06 | 6.60E-07 | 1.93E-06 | 6.86E-07 | 4.31E-07 | 1.21E-06 | 6.17E-07 | 1.23E-06 |
| 389 | 187.01839 | C14H3O+ | 2.22E-07 | 6.70E-07 | 1.60E-07 | 5.44E-07 | 3.88E-07 | 1.29E-06 | 2.86E-07 | 1.21E-06 |
| 390 | 187.03952 | C11H7O3+ | 9.04E-07 | 2.32E-06 | 5.75E-07 | 2.13E-06 | 1.03E-06 | 2.75E-06 | 7.87E-07 | 2.58E-06 |
| 391 | 187.09703 | C9H15O4+ | 5.77E-07 | 7.94E-07 | 4.17E-07 | 7.09E-07 | 3.21E-06 | 1.91E-06 | 2.37E-06 | 1.98E-06 |
| 392 | 187.11229 | C13H15O+ | 2.01E-07 | 6.11E-07 | 1.43E-07 | 4.94E-07 | 3.68E-07 | 1.16E-06 | 2.74E-07 | 1.09E-06 |
| 393 | 187.13342 | C10H19O3+ | 2.33E-07 | 7.05E-07 | 1.65E-07 | 5.71E-07 | 4.53E-07 | 2.29E-06 | 3.67E-07 | 2.03E-06 |
| 394 | 187.14868 | C14H19+ | 1.35E-07 | 4.03E-06 | 9.33E-08 | 2.62E-06 | 3.27E-07 | 4.59E-06 | 2.82E-07 | 3.68E-06 |
| 395 | 187.1698 | C11H23O2+ | 2.47E-07 | 6.81E-07 | 1.78E-07 | 5.66E-07 | 3.84E-07 | 1.26E-06 | 2.98E-07 | 1.20E-06 |
| 396 | 189.05518 | C11H9O3+ | 3.83E-07 | 1.16E-06 | 2.98E-07 | 1.12E-06 | 3.66E-07 | 1.31E-06 | 2.68E-07 | 1.21E-06 |
| 397 | 189.07042 | C15H9+ | 2.72E-07 | 7.43E-07 | 1.97E-07 | 5.94E-07 | 2.11E-06 | 2.47E-06 | 1.71E-06 | 2.19E-06 |
| 398 | 189.16432 | C14H21+ | 4.03E-07 | 4.68E-06 | 2.68E-07 | 3.14E-06 | 1.08E-06 | 1.80E-05 | 7.91E-07 | 1.62E-05 |
| 399 | 189.184906 | C11H25O2+ | 8.67E-07 | 9.37E-07 | 6.03E-07 | 8.34E-07 | 4.08E-07 | 1.26E-06 | 5.49E-07 | 1.19E-06 |
| 400 | 190.98279 | C5H3O8+ | 2.94E-07 | 7.53E-07 | 1.95E-07 | 5.96E-07 | 3.88E-07 | 1.22E-06 | 2.82E-07 | 1.16E-06 |

**Table S1.** Continued.

| | | | 2024 | | | | 2025 | | | |
|---|---|---|---|---|---|---|---|---|---|---|
| | | | Forest-interior (FI) atmospheres | | Canopy-top (CT) atmospheres | | Forest-interior (FI) atmospheres | | Canopy-top (CT) atmospheres | |
| ID | Exact m/z | Ion formula | Daytime-FI | Nighttime-FI | Daytime-CT | Nighttime-CT | Daytime-FI | Nighttime-FI | Daytime-CT | Nighttime-CT |
| 401 | 191.03444 | C10H7O4+ | 3.52E-07 | 9.17E-07 | 2.67E-07 | 7.99E-07 | 6.01E-07 | 1.98E-06 | 4.31E-07 | 1.93E-06 |
| 402 | 191.04968 | C14H7O+ | 1.97E-07 | 6.47E-07 | 1.40E-07 | 5.23E-07 | 3.40E-07 | 1.13E-06 | 2.53E-07 | 1.07E-06 |
| 403 | 191.05556 | C7H11O6+ | 2.00E-07 | 6.62E-07 | 1.42E-07 | 5.37E-07 | 3.36E-07 | 1.14E-06 | 2.50E-07 | 1.06E-06 |
| 404 | 191.07082 | C11H11O3+ | 2.56E-07 | 7.54E-07 | 1.80E-07 | 6.05E-07 | 5.96E-07 | 1.51E-06 | 4.54E-07 | 1.41E-06 |
| 405 | 191.158346 | <unknown0236> | 3.80E-06 | 1.88E-06 | 2.39E-06 | 1.51E-06 | 6.16E-07 | 1.78E-06 | 5.52E-07 | 1.66E-06 |
| 406 | 191.17998 | C14H23+ | 2.84E-07 | 1.66E-06 | 2.02E-07 | 1.55E-06 | 4.55E-07 | 2.06E-06 | 3.95E-07 | 1.89E-06 |
| 407 | 192.138291 | C12H18NO+ | 3.03E-07 | 5.63E-07 | 4.02E-07 | 4.20E-07 | 3.30E-07 | 1.09E-06 | 2.55E-07 | 1.04E-06 |
| 408 | 193.02896 | C13H5O2+ | 2.44E-07 | 7.26E-07 | 1.73E-07 | 5.89E-07 | 3.99E-07 | 1.34E-06 | 3.00E-07 | 1.27E-06 |
| 409 | 193.03482 | C6H9O7+ | 2.08E-07 | 6.80E-07 | 1.47E-07 | 5.47E-07 | 3.64E-07 | 1.22E-06 | 2.69E-07 | 1.15E-06 |
| 410 | 193.05008 | C10H9O4+ | 2.77E-07 | 8.61E-07 | 2.07E-07 | 7.07E-07 | 5.46E-07 | 1.56E-06 | 4.08E-07 | 1.46E-06 |
| 411 | 193.15924 | C13H21O+ | 1.46E-06 | 5.75E-06 | 1.01E-06 | 3.02E-06 | 1.04E-06 | 1.08E-05 | 1.03E-06 | 8.11E-06 |
| 412 | 193.19563 | C14H25+ | 2.69E-07 | 7.53E-07 | 2.08E-07 | 7.66E-07 | 3.84E-07 | 1.38E-06 | 3.25E-07 | 1.33E-06 |
| 413 | 195.05048 | C6H11O7+ | 3.34E-07 | 8.67E-07 | 2.41E-07 | 7.33E-07 | 4.88E-07 | 1.80E-06 | 3.51E-07 | 1.79E-06 |
| 414 | 195.06573 | C10H11O4+ | 1.99E-07 | 6.54E-07 | 1.43E-07 | 5.30E-07 | 3.48E-07 | 1.13E-06 | 2.64E-07 | 1.08E-06 |
| 415 | 195.08099 | C14H11O+ | 1.90E-07 | 6.08E-07 | 1.36E-07 | 4.88E-07 | 3.34E-07 | 1.10E-06 | 2.49E-07 | 1.05E-06 |
| 416 | 195.08687 | C7H15O6+ | 2.97E-07 | 9.06E-07 | 1.87E-07 | 7.08E-07 | 5.67E-07 | 1.60E-06 | 4.58E-07 | 1.42E-06 |
| 417 | 195.1749 | C13H23O+ | 0.00021152 | 4.16E-05 | 0.00014935 | 2.39E-05 | 1.25E-06 | 1.04E-05 | 2.75E-06 | 9.01E-06 |
| 418 | 195.21127 | C14H27+ | 1.84E-07 | 6.31E-07 | 1.30E-07 | 5.12E-07 | 3.52E-07 | 1.18E-06 | 2.85E-07 | 1.16E-06 |
| 419 | 197.00861 | C8H5O6+ | 6.42E-07 | 1.17E-06 | 3.40E-07 | 1.09E-06 | 8.07E-07 | 2.17E-06 | 4.83E-07 | 1.78E-06 |
| 420 | 197.09663 | C14H13O+ | 6.04E-05 | 2.85E-05 | 9.55E-05 | 0.00017439 | 8.42E-06 | 1.29E-05 | 1.05E-05 | 8.49E-06 |
| 421 | 197.19054 | C13H25O+ | 3.83E-06 | 6.17E-06 | 2.61E-06 | 3.44E-06 | 6.22E-07 | 2.78E-06 | 1.42E-06 | 2.97E-06 |
| 422 | 199.03952 | C12H7O3+ | 2.77E-07 | 7.57E-07 | 1.94E-07 | 6.23E-07 | 6.36E-07 | 2.00E-06 | 4.44E-07 | 1.86E-06 |
| 423 | 199.04539 | C5H11O8+ | 1.96E-07 | 6.37E-07 | 1.38E-07 | 5.18E-07 | 3.27E-07 | 1.10E-06 | 2.42E-07 | 1.04E-06 |
| 424 | 199.05478 | C16H7+ | 1.83E-07 | 6.24E-07 | 1.29E-07 | 4.99E-07 | 3.24E-07 | 1.10E-06 | 2.40E-07 | 1.03E-06 |
| 425 | 199.06065 | C9H11O5+ | 1.87E-07 | 5.93E-07 | 1.32E-07 | 4.81E-07 | 3.25E-07 | 1.10E-06 | 2.43E-07 | 1.04E-06 |
| 426 | 199.0759 | C13H11O2+ | 7.28E-07 | 2.05E-06 | 5.71E-07 | 1.49E-06 | 2.72E-06 | 3.14E-06 | 1.90E-06 | 2.67E-06 |
| 427 | 199.20619 | C13H27O+ | 0.00013059 | 0.00015659 | 8.85E-05 | 0.00012938 | 9.05E-07 | 1.91E-06 | 2.79E-06 | 2.21E-06 |
| 428 | 201.01878 | C11H5O4+ | 2.35E-07 | 6.66E-07 | 1.70E-07 | 5.47E-07 | 4.57E-07 | 1.58E-06 | 3.31E-07 | 1.51E-06 |
| 429 | 201.05518 | C12H9O3+ | 2.85E-05 | 9.67E-05 | 2.26E-05 | 8.14E-05 | 5.03E-05 | 9.30E-05 | 3.26E-05 | 6.68E-05 |
| 430 | 201.16432 | C15H21+ | 1.55E-06 | 1.09E-05 | 9.63E-07 | 6.82E-06 | 1.52E-06 | 2.54E-05 | 2.29E-06 | 2.05E-05 |
| 431 | 201.18546 | C12H25O2+ | 2.02E-07 | 6.61E-07 | 1.47E-07 | 5.37E-07 | 3.43E-07 | 1.19E-06 | 2.61E-07 | 1.12E-06 |
| 432 | 203.01331 | C14H3O2+ | 2.59E-07 | 7.22E-07 | 1.78E-07 | 5.99E-07 | 4.14E-07 | 1.34E-06 | 3.00E-07 | 1.23E-06 |
| 433 | 203.03444 | C11H7O4+ | 3.37E-07 | 8.79E-07 | 2.26E-07 | 7.51E-07 | 3.79E-07 | 1.28E-06 | 2.77E-07 | 1.18E-06 |
| 434 | 203.04968 | C15H7O+ | 2.14E-07 | 6.43E-07 | 1.51E-07 | 5.25E-07 | 3.41E-07 | 1.17E-06 | 2.52E-07 | 1.09E-06 |
| 435 | 203.05556 | C8H11O6+ | 2.13E-07 | 6.81E-07 | 1.54E-07 | 5.51E-07 | 4.33E-07 | 1.21E-06 | 3.18E-07 | 1.13E-06 |
| 436 | 203.12834 | C10H19O4+ | 1.80E-07 | 5.75E-07 | 1.27E-07 | 4.62E-07 | 4.01E-07 | 1.02E-06 | 2.97E-07 | 9.53E-07 |
| 437 | 203.17998 | C15H23+ | 0 | 0.0014279 | 0 | 0.00022956 | 0 | 0.00128532 | 0 | 0.000858 |
| 438 | 203.200556 | C12H27O2+ | 3.11E-07 | 8.90E-07 | 2.38E-07 | 7.41E-07 | 3.92E-07 | 1.25E-06 | 3.21E-07 | 1.19E-06 |
| 439 | 204.12104 | <unknown0259> | 2.43E-07 | 5.90E-07 | 1.78E-07 | 4.78E-07 | 3.90E-07 | 1.20E-06 | 2.90E-07 | 1.13E-06 |
| 440 | 204.194124 | <unknown0260> | 2.53E-07 | 2.29E-06 | 1.63E-07 | 1.66E-06 | 4.40E-07 | 3.59E-06 | 3.59E-07 | 3.15E-06 |
| 441 | 205.05008 | C11H9O4+ | 1.24E-06 | 4.95E-07 | 1.06E-06 | 5.02E-07 | 8.43E-07 | 1.19E-06 | 5.20E-07 | 1.13E-06 |
| 442 | 205.14398 | C10H21O4+ | 1.82E-07 | 5.81E-07 | 1.30E-07 | 4.66E-07 | 3.25E-07 | 1.15E-06 | 2.40E-07 | 1.05E-06 |
| 443 | 205.15924 | C14H21O+ | 1.99E-07 | 5.77E-07 | 1.45E-07 | 4.63E-07 | 3.37E-07 | 1.43E-06 | 2.49E-07 | 1.26E-06 |
| 444 | 205.19563 | C15H25+ | 0.00110659 | 0.01510593 | 0.00065211 | 0.01112338 | 0.00128517 | 0.0225739 | 0.00144446 | 0.01974359 |
| 445 | 205.216207 | C12H29O2+ | 1.67E-07 | 7.18E-07 | 1.18E-07 | 5.85E-07 | 3.79E-07 | 1.16E-06 | 2.81E-07 | 1.09E-06 |
| 446 | 207.0141 | C6H7O8+ | 1.21E-06 | 1.78E-06 | 1.21E-06 | 1.74E-06 | 7.25E-07 | 1.95E-06 | 5.17E-07 | 1.75E-06 |
| 447 | 207.21127 | C15H27+ | 7.74E-06 | 1.28E-05 | 3.88E-06 | 1.46E-05 | 1.35E-06 | 7.07E-06 | 1.35E-06 | 6.52E-06 |
| 448 | 209.02386 | C13H5O3+ | 2.35E-07 | 6.81E-07 | 1.65E-07 | 5.51E-07 | 3.62E-07 | 1.18E-06 | 2.70E-07 | 1.12E-06 |
| 449 | 209.02974 | C6H9O8+ | 1.90E-07 | 6.49E-07 | 1.36E-07 | 5.23E-07 | 3.41E-07 | 1.16E-06 | 2.53E-07 | 1.09E-06 |
| 450 | 209.03912 | C17H5+ | 2.05E-07 | 6.30E-07 | 1.45E-07 | 5.13E-07 | 3.45E-07 | 1.15E-06 | 2.56E-07 | 1.07E-06 |

**Table S1.** Continued.

| | | | 2024 | | | | 2025 | | | |
|---|---|---|---|---|---|---|---|---|---|---|
| | | | Forest-interior (FI) atmospheres | | Canopy-top (CT) atmospheres | | Forest-interior (FI) atmospheres | | Canopy-top (CT) atmospheres | |
| ID | Exact m/z | Ion formula | Daytime-FI | Nighttime-FI | Daytime-CT | Nighttime-CT | Daytime-FI | Nighttime-FI | Daytime-CT | Nighttime-CT |
| 451 | 209.045 | C10H9O5+ | 2.53E-07 | 8.53E-07 | 1.82E-07 | 6.81E-07 | 4.36E-07 | 1.37E-06 | 3.21E-07 | 1.26E-06 |
| 452 | 209.19054 | C14H25O+ | 5.76E-06 | 1.73E-05 | 2.53E-06 | 1.24E-05 | 9.62E-07 | 6.26E-06 | 9.62E-07 | 4.99E-06 |
| 453 | 209.22693 | C15H29+ | 1.87E-07 | 6.02E-07 | 1.33E-07 | 4.87E-07 | 3.41E-07 | 1.13E-06 | 2.81E-07 | 1.09E-06 |
| 454 | 211.00313 | C12H3O4+ | 2.00E-07 | 6.44E-07 | 1.44E-07 | 5.19E-07 | 3.51E-07 | 1.17E-06 | 2.58E-07 | 1.09E-06 |
| 455 | 211.009 | C5H7O9+ | 1.95E-07 | 6.09E-07 | 1.39E-07 | 4.93E-07 | 3.36E-07 | 1.14E-06 | 2.52E-07 | 1.06E-06 |
| 456 | 211.01839 | C16H3O+ | 1.82E-07 | 6.08E-07 | 1.29E-07 | 4.91E-07 | 3.36E-07 | 1.12E-06 | 2.48E-07 | 1.06E-06 |
| 457 | 211.02426 | C9H7O6+ | 2.00E-07 | 6.31E-07 | 1.42E-07 | 5.10E-07 | 3.50E-07 | 1.18E-06 | 2.57E-07 | 1.09E-06 |
| 458 | 211.03952 | C13H7O3+ | 2.41E-07 | 7.28E-07 | 1.63E-07 | 5.93E-07 | 3.86E-07 | 1.25E-06 | 2.82E-07 | 1.17E-06 |
| 459 | 211.0759 | C14H11O2+ | 2.08E-07 | 6.70E-07 | 1.45E-07 | 5.41E-07 | 3.59E-07 | 1.19E-06 | 2.62E-07 | 1.11E-06 |
| 460 | 211.08177 | C7H15O7+ | 2.53E-07 | 7.45E-07 | 1.92E-07 | 6.09E-07 | 4.98E-07 | 1.43E-06 | 3.85E-07 | 1.33E-06 |
| 461 | 211.1698 | C13H23O2+ | 5.96E-07 | 2.76E-06 | 4.24E-07 | 2.00E-06 | 8.71E-07 | 8.35E-06 | 9.81E-07 | 6.44E-06 |
| 462 | 211.20619 | C14H27O+ | 3.30E-07 | 6.73E-07 | 2.68E-07 | 5.44E-07 | 3.26E-07 | 1.09E-06 | 2.48E-07 | 1.03E-06 |
| 463 | 211.24257 | C15H31+ | 2.21E-07 | 6.74E-07 | 1.64E-07 | 5.55E-07 | 3.46E-07 | 1.13E-06 | 3.07E-07 | 1.11E-06 |
| 464 | 213.03404 | C16H5O+ | 2.04E-07 | 6.49E-07 | 1.45E-07 | 5.28E-07 | 3.64E-07 | 1.19E-06 | 2.67E-07 | 1.11E-06 |
| 465 | 213.03992 | C9H9O6+ | 2.07E-07 | 6.57E-07 | 1.48E-07 | 5.30E-07 | 3.43E-07 | 1.18E-06 | 2.54E-07 | 1.11E-06 |
| 466 | 213.05518 | C13H9O3+ | 2.81E-07 | 8.54E-07 | 1.87E-07 | 6.78E-07 | 5.73E-07 | 1.54E-06 | 4.04E-07 | 1.45E-06 |
| 467 | 213.16432 | C16H21+ | 2.51E-07 | 9.64E-07 | 1.63E-07 | 7.39E-07 | 6.73E-07 | 2.10E-06 | 5.49E-07 | 1.84E-06 |
| 468 | 213.18546 | C13H25O2+ | 5.38E-07 | 1.02E-06 | 4.76E-07 | 7.89E-07 | 3.46E-07 | 1.20E-06 | 2.66E-07 | 1.14E-06 |
| 469 | 213.22185 | C14H29O+ | 2.63E-07 | 8.29E-07 | 1.78E-07 | 6.88E-07 | 3.26E-07 | 1.07E-06 | 2.56E-07 | 1.00E-06 |
| 470 | 215.01918 | C8H7O7+ | 3.09E-07 | 7.65E-07 | 2.09E-07 | 6.41E-07 | 4.30E-07 | 1.37E-06 | 3.01E-07 | 1.24E-06 |
| 471 | 215.04968 | C16H7O+ | 2.33E-07 | 7.26E-07 | 1.64E-07 | 5.91E-07 | 3.65E-07 | 1.21E-06 | 2.65E-07 | 1.14E-06 |
| 472 | 215.05556 | C9H11O6+ | 2.01E-07 | 6.37E-07 | 1.42E-07 | 5.15E-07 | 3.48E-07 | 1.18E-06 | 2.59E-07 | 1.11E-06 |
| 473 | 215.07082 | C13H11O3+ | 1.98E-07 | 6.47E-07 | 1.41E-07 | 5.18E-07 | 3.96E-07 | 1.28E-06 | 2.94E-07 | 1.21E-06 |
| 474 | 215.08607 | C17H11+ | 2.09E-07 | 6.93E-07 | 1.49E-07 | 5.50E-07 | 4.06E-07 | 1.27E-06 | 2.99E-07 | 1.17E-06 |
| 475 | 215.10721 | C14H15O2+ | 2.97E-07 | 8.96E-07 | 2.04E-07 | 7.23E-07 | 5.51E-07 | 1.66E-06 | 4.26E-07 | 1.50E-06 |
| 476 | 215.17998 | C16H23+ | 5.63E-07 | 1.20E-06 | 3.72E-07 | 1.01E-06 | 4.94E-07 | 1.59E-06 | 3.89E-07 | 1.50E-06 |
| 477 | 215.20111 | C13H27O2+ | 2.05E-07 | 6.64E-07 | 1.51E-07 | 5.36E-07 | 3.40E-07 | 1.15E-06 | 2.60E-07 | 1.08E-06 |
| 478 | 217.0137 | C11H5O5+ | 2.04E-07 | 6.20E-07 | 1.39E-07 | 5.00E-07 | 3.64E-07 | 1.18E-06 | 2.62E-07 | 1.09E-06 |
| 479 | 217.02896 | C15H5O2+ | 2.24E-07 | 6.36E-07 | 1.55E-07 | 5.11E-07 | 3.75E-07 | 1.27E-06 | 2.77E-07 | 1.19E-06 |
| 480 | 217.05008 | C12H9O4+ | 9.16E-07 | 4.67E-06 | 5.71E-07 | 3.91E-06 | 1.42E-06 | 3.26E-06 | 9.70E-07 | 3.36E-06 |
| 481 | 217.19563 | C16H25+ | 1.18E-06 | 3.75E-06 | 7.87E-07 | 3.03E-06 | 7.41E-07 | 3.52E-06 | 7.19E-07 | 3.19E-06 |
| 482 | 217.216207 | C13H29O2+ | 2.35E-07 | 6.70E-07 | 1.63E-07 | 5.41E-07 | 3.59E-07 | 1.13E-06 | 2.82E-07 | 1.08E-06 |
| 483 | 218.211709 | <unknown0278> | 2.40E-07 | 7.09E-07 | 1.76E-07 | 5.66E-07 | 3.61E-07 | 1.22E-06 | 2.76E-07 | 1.15E-06 |
| 484 | 219.02347 | C18H3+ | 1.86E-07 | 5.87E-07 | 1.31E-07 | 4.72E-07 | 3.34E-07 | 1.12E-06 | 2.49E-07 | 1.04E-06 |
| 485 | 219.05048 | C8H11O7+ | 4.89E-05 | 1.04E-05 | 1.94E-05 | 9.26E-06 | 1.01E-05 | 6.06E-05 | 8.01E-06 | 4.81E-05 |
| 486 | 219.08099 | C16H11O+ | 1.75E-07 | 8.86E-07 | 1.23E-07 | 7.22E-07 | 4.19E-07 | 1.40E-06 | 3.22E-07 | 1.33E-06 |
| 487 | 219.08687 | C9H15O6+ | 1.86E-07 | 6.30E-07 | 1.33E-07 | 5.06E-07 | 3.35E-07 | 1.12E-06 | 2.49E-07 | 1.06E-06 |
| 488 | 219.10211 | C13H15O3+ | 1.82E-07 | 5.87E-07 | 1.28E-07 | 4.71E-07 | 3.56E-07 | 1.17E-06 | 2.65E-07 | 1.09E-06 |
| 489 | 219.185021 | <unknown0279> | 3.90E-06 | 7.46E-05 | 3.17E-06 | 1.32E-05 | 8.11E-07 | 0.00042439 | 1.04E-06 | 0.00020493 |
| 490 | 219.21127 | C16H27+ | 2.14E-07 | 6.46E-07 | 1.75E-07 | 5.33E-07 | 3.41E-07 | 1.09E-06 | 2.81E-07 | 1.04E-06 |
| 491 | 220.197715 | <unknown0281> | 4.96E-07 | 2.16E-06 | 3.68E-07 | 1.47E-06 | 4.62E-07 | 3.44E-06 | 4.03E-07 | 2.84E-06 |
| 492 | 221.02974 | C7H9O8+ | 2.42E-07 | 7.02E-07 | 1.71E-07 | 5.78E-07 | 3.88E-07 | 1.24E-06 | 2.82E-07 | 1.16E-06 |
| 493 | 221.03912 | C18H5+ | 2.06E-07 | 6.69E-07 | 1.44E-07 | 5.38E-07 | 3.46E-07 | 1.14E-06 | 2.55E-07 | 1.08E-06 |
| 494 | 221.045 | C11H9O5+ | 2.16E-07 | 6.63E-07 | 1.60E-07 | 5.40E-07 | 3.66E-07 | 1.18E-06 | 2.72E-07 | 1.09E-06 |
| 495 | 221.1389 | C10H21O5+ | 1.79E-07 | 5.77E-07 | 1.26E-07 | 4.62E-07 | 3.16E-07 | 1.04E-06 | 2.36E-07 | 9.76E-07 |
| 496 | 221.15416 | C14H21O2+ | 3.35E-07 | 6.46E-07 | 2.20E-07 | 5.34E-07 | 5.00E-07 | 1.42E-06 | 3.67E-07 | 1.33E-06 |
| 497 | 221.19054 | C15H25O+ | 0 | 0.00080518 | 0 | 9.62E-05 | 3.63E-11 | 0.00152109 | 0 | 0.00104945 |
| 498 | 221.22693 | C16H29+ | 2.27E-07 | 7.61E-07 | 1.71E-07 | 6.37E-07 | 4.05E-07 | 1.32E-06 | 3.30E-07 | 1.26E-06 |
| 499 | 222.98788 | C9H3O7+ | 1.98E-07 | 6.29E-07 | 1.42E-07 | 5.11E-07 | 3.52E-07 | 1.15E-06 | 2.57E-07 | 1.06E-06 |
| 500 | 223.00313 | C13H3O4+ | 1.99E-07 | 6.22E-07 | 1.42E-07 | 5.04E-07 | 3.91E-07 | 1.23E-06 | 2.83E-07 | 1.15E-06 |

**Table S1.** Continued.

| | | | 2024 | | | | 2025 | | | |
|---|---|---|---|---|---|---|---|---|---|---|
| | | | Forest-interior (FI) atmospheres | | Canopy-top (CT) atmospheres | | Forest-interior (FI) atmospheres | | Canopy-top (CT) atmospheres | |
| ID | Exact m/z | Ion formula | Daytime-FI | Nighttime-FI | Daytime-CT | Nighttime-CT | Daytime-FI | Nighttime-FI | Daytime-CT | Nighttime-CT |
| 501 | 223.06065 | C11H11O5+ | 6.32E-07 | 8.61E-07 | 4.77E-07 | 7.62E-07 | 5.58E-07 | 1.68E-06 | 4.15E-07 | 1.58E-06 |
| 502 | 223.08177 | C8H15O7+ | 2.15E-07 | 7.89E-07 | 1.43E-07 | 6.17E-07 | 3.33E-07 | 1.13E-06 | 2.50E-07 | 1.06E-06 |
| 503 | 223.09703 | C12H15O4+ | 1.88E-07 | 6.09E-07 | 1.31E-07 | 4.85E-07 | 3.21E-07 | 1.06E-06 | 2.39E-07 | 9.95E-07 |
| 504 | 223.11229 | C16H15O+ | 1.82E-07 | 5.91E-07 | 1.28E-07 | 4.73E-07 | 3.29E-07 | 1.08E-06 | 2.45E-07 | 1.02E-06 |
| 505 | 223.11816 | C9H19O6+ | 1.84E-07 | 5.99E-07 | 1.30E-07 | 4.77E-07 | 3.29E-07 | 1.09E-06 | 2.47E-07 | 1.02E-06 |
| 506 | 223.13342 | C13H19O3+ | 2.06E-07 | 6.19E-07 | 1.48E-07 | 4.98E-07 | 3.96E-07 | 1.26E-06 | 3.00E-07 | 1.18E-06 |
| 507 | 223.1698 | C14H23O2+ | 3.42E-07 | 2.37E-06 | 3.20E-07 | 1.57E-06 | 7.01E-07 | 6.46E-06 | 6.03E-07 | 4.55E-06 |
| 508 | 223.20619 | C15H27O+ | 2.34E-07 | 1.41E-06 | 1.51E-07 | 1.12E-06 | 4.39E-07 | 2.65E-06 | 3.39E-07 | 2.39E-06 |
| 509 | 223.24257 | C16H31+ | 2.05E-07 | 6.51E-07 | 1.47E-07 | 5.31E-07 | 3.57E-07 | 1.25E-06 | 3.12E-07 | 1.22E-06 |
| 510 | 225.02466 | C6H9O9+ | 2.71E-07 | 7.06E-07 | 1.98E-07 | 5.83E-07 | 5.11E-07 | 1.51E-06 | 3.42E-07 | 1.39E-06 |
| 511 | 225.06104 | C7H13O8+ | 5.86E-07 | 1.79E-06 | 4.21E-07 | 1.47E-06 | 5.67E-07 | 1.71E-06 | 4.42E-07 | 1.52E-06 |
| 512 | 225.25822 | C16H33+ | 1.90E-06 | 1.13E-06 | 1.92E-06 | 7.94E-07 | 3.39E-07 | 1.14E-06 | 3.96E-07 | 1.18E-06 |
| 513 | 227.04968 | C17H7O+ | 2.56E-07 | 7.10E-07 | 1.79E-07 | 5.84E-07 | 3.73E-07 | 1.22E-06 | 2.73E-07 | 1.13E-06 |
| 514 | 227.05556 | C10H11O6+ | 1.94E-07 | 6.48E-07 | 1.38E-07 | 5.20E-07 | 3.42E-07 | 1.16E-06 | 2.54E-07 | 1.09E-06 |
| 515 | 227.07082 | C14H11O3+ | 1.92E-07 | 6.25E-07 | 1.33E-07 | 5.05E-07 | 3.39E-07 | 1.15E-06 | 2.51E-07 | 1.09E-06 |
| 516 | 227.07669 | C7H15O8+ | 2.21E-07 | 7.18E-07 | 1.57E-07 | 5.66E-07 | 4.45E-07 | 1.35E-06 | 3.32E-07 | 1.26E-06 |
| 517 | 227.20111 | C14H27O2+ | 4.85E-07 | 1.42E-06 | 3.90E-07 | 1.06E-06 | 4.33E-07 | 1.56E-06 | 3.71E-07 | 1.45E-06 |
| 518 | 227.23749 | C15H31O+ | 3.14E-07 | 6.92E-07 | 2.32E-07 | 5.67E-07 | 3.15E-07 | 1.05E-06 | 2.39E-07 | 9.87E-07 |
| 519 | 229.0137 | C12H5O5+ | 1.99E-07 | 6.36E-07 | 1.40E-07 | 5.13E-07 | 3.56E-07 | 1.16E-06 | 2.62E-07 | 1.09E-06 |
| 520 | 229.01958 | C5H9O10+ | 1.95E-07 | 6.14E-07 | 1.39E-07 | 4.93E-07 | 3.37E-07 | 1.14E-06 | 2.48E-07 | 1.07E-06 |
| 521 | 229.02896 | C16H5O2+ | 1.85E-07 | 6.05E-07 | 1.30E-07 | 4.86E-07 | 3.32E-07 | 1.12E-06 | 2.46E-07 | 1.06E-06 |
| 522 | 229.03482 | C9H9O7+ | 1.97E-07 | 6.23E-07 | 1.37E-07 | 5.01E-07 | 3.42E-07 | 1.13E-06 | 2.55E-07 | 1.05E-06 |
| 523 | 229.05008 | C13H9O4+ | 2.01E-07 | 6.10E-07 | 1.39E-07 | 4.93E-07 | 3.42E-07 | 1.15E-06 | 2.52E-07 | 1.06E-06 |
| 524 | 229.05595 | C6H13O9+ | 1.89E-07 | 6.36E-07 | 1.32E-07 | 5.09E-07 | 3.26E-07 | 1.10E-06 | 2.41E-07 | 1.03E-06 |
| 525 | 229.06534 | C17H9O+ | 1.91E-07 | 6.00E-07 | 1.34E-07 | 4.84E-07 | 3.38E-07 | 1.11E-06 | 2.48E-07 | 1.04E-06 |
| 526 | 229.07121 | C10H13O6+ | 1.91E-07 | 6.38E-07 | 1.34E-07 | 5.10E-07 | 3.33E-07 | 1.12E-06 | 2.47E-07 | 1.05E-06 |
| 527 | 229.08647 | C14H13O3+ | 2.35E-07 | 7.24E-07 | 1.75E-07 | 5.84E-07 | 5.04E-07 | 1.45E-06 | 3.65E-07 | 1.33E-06 |
| 528 | 229.18037 | C13H25O3+ | 2.73E-07 | 8.55E-07 | 1.98E-07 | 6.83E-07 | 4.47E-07 | 1.44E-06 | 3.57E-07 | 1.34E-06 |
| 529 | 229.19563 | C17H25+ | 2.18E-07 | 6.68E-07 | 1.59E-07 | 5.52E-07 | 3.73E-07 | 1.21E-06 | 2.87E-07 | 1.14E-06 |
| 530 | 229.21675 | C14H29O2+ | 2.65E-07 | 7.09E-07 | 2.02E-07 | 5.80E-07 | 3.47E-07 | 1.13E-06 | 2.63E-07 | 1.08E-06 |
| 531 | 231.06573 | C13H11O4+ | 4.88E-07 | 1.18E-06 | 3.12E-07 | 9.67E-07 | 6.55E-07 | 1.73E-06 | 4.26E-07 | 1.58E-06 |
| 532 | 231.1749 | C16H23O+ | 2.91E-07 | 7.05E-07 | 1.80E-07 | 5.48E-07 | 3.85E-07 | 1.25E-06 | 2.88E-07 | 1.18E-06 |
| 533 | 231.19601 | C13H27O3+ | 1.96E-07 | 6.37E-07 | 1.38E-07 | 5.17E-07 | 3.54E-07 | 1.18E-06 | 2.73E-07 | 1.12E-06 |
| 534 | 231.21127 | C17H27+ | 2.72E-07 | 6.94E-07 | 2.39E-07 | 5.87E-07 | 3.63E-07 | 1.17E-06 | 3.07E-07 | 1.14E-06 |
| 535 | 231.241976 | <unknown0292> | 2.84E-07 | 7.62E-07 | 1.98E-07 | 6.39E-07 | 3.34E-07 | 1.11E-06 | 2.64E-07 | 1.05E-06 |

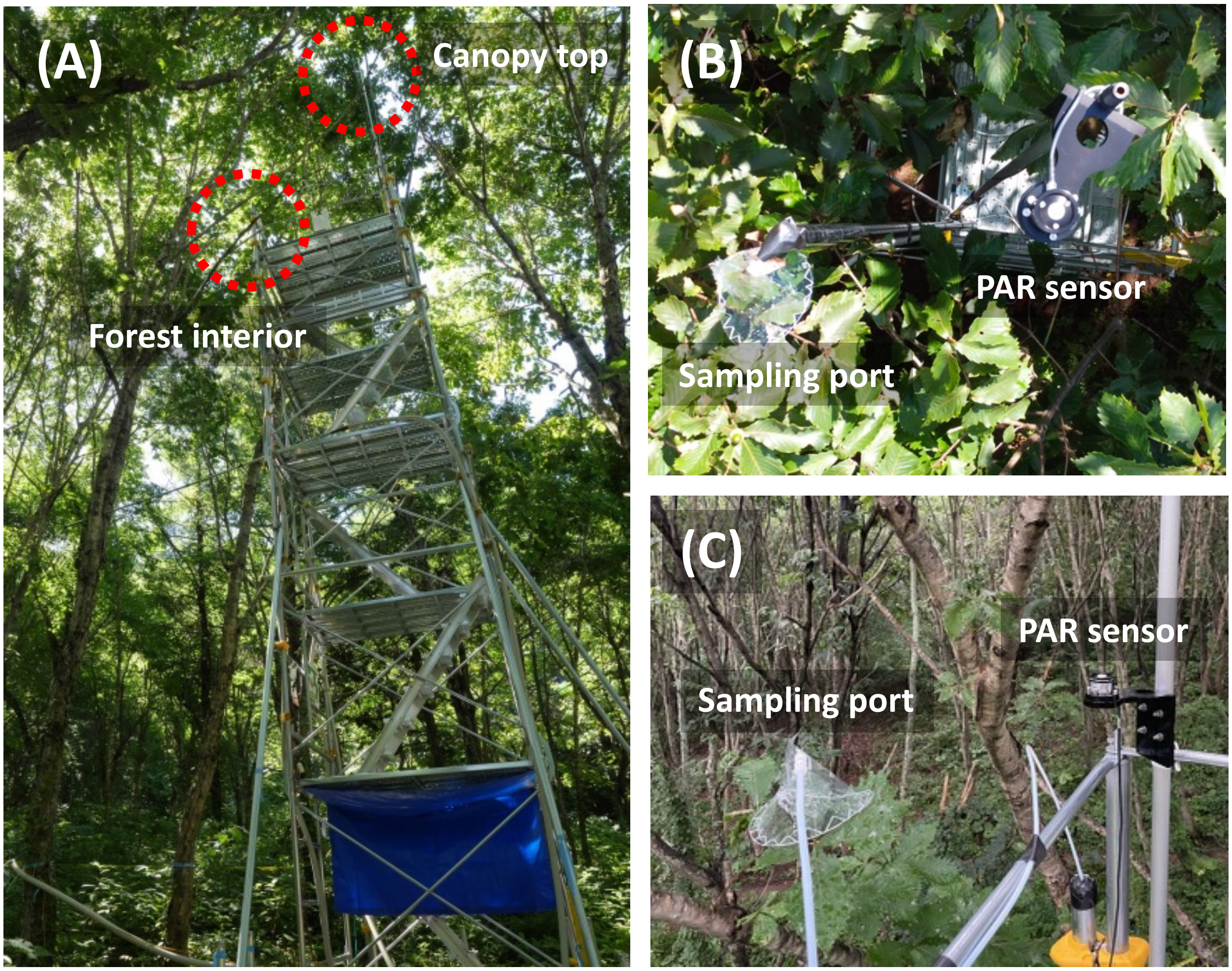


**Figure S1. Study site and sampling locations.** (A) Rolling tower used for air sampling. Locations of air sampling ports and PAR sensors at the (B) canopy top and (C) forest interior.

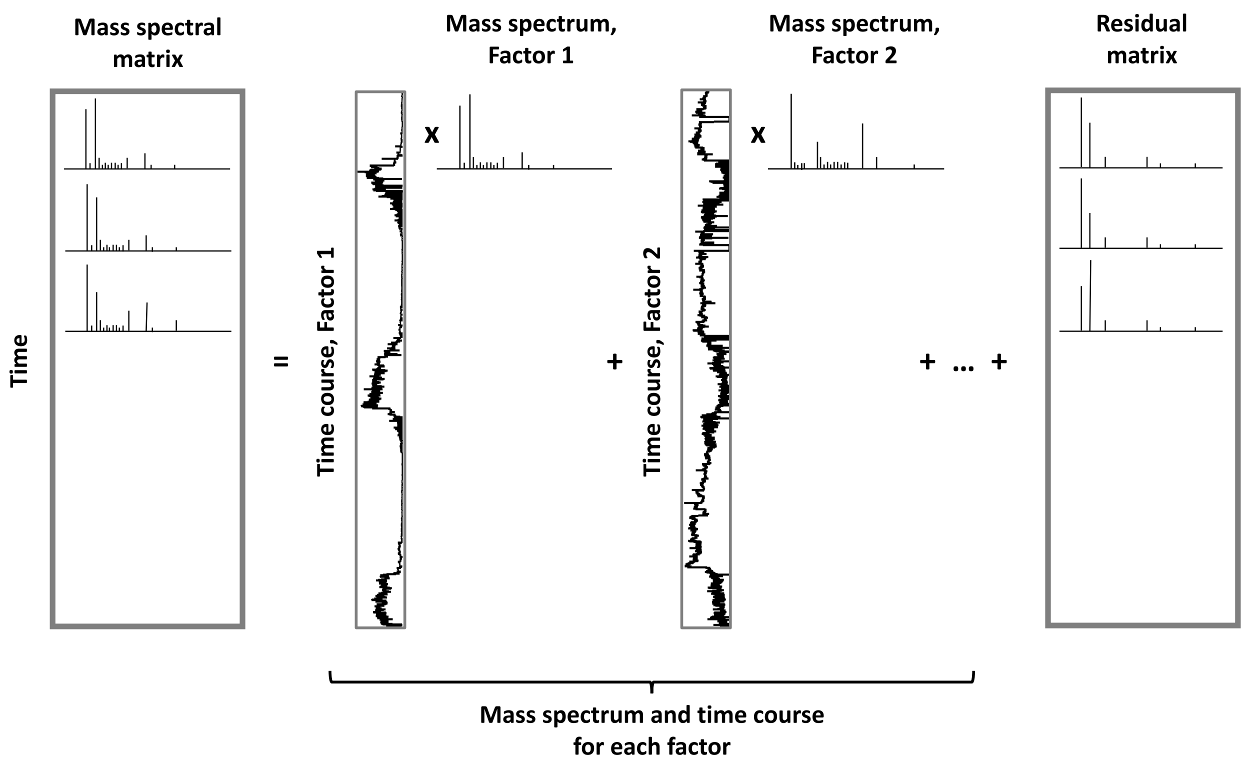


**Figure S2. Schematic representation of PMF factorization.** Factor time courses constitute the matrix ***g***, and factor mass spectra constitute the matrix ***f*** in Eq. (1).

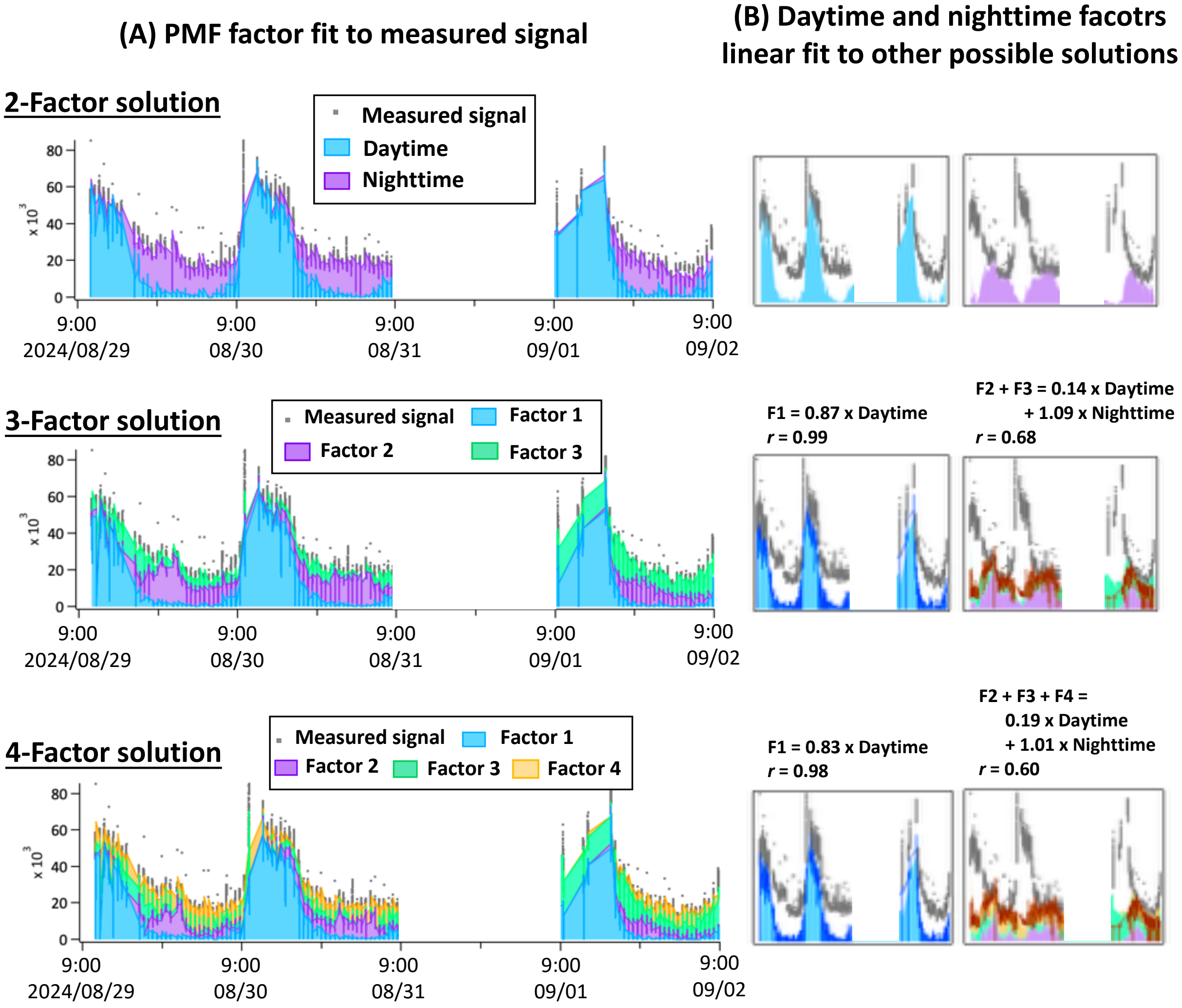


**Figure S3. Comparison of time courses for the forest-interior atmospheres in the 2024 dataset from 2-, 3-, and 4-factor PMF solutions.** The left panels (A) show the stacked contributions of the factors, compared with the measured signal. The right panels (B) show the time courses of individual factors (shaded areas). The daytime and nighttime factors were fitted to corresponding factors in the 3- and 4-factor solutions. Best fits were obtained using the extended time courses. These best-fits are shown as solid lines in the right panels (B), and the best-fit equation and correlation coefficient are also provided.

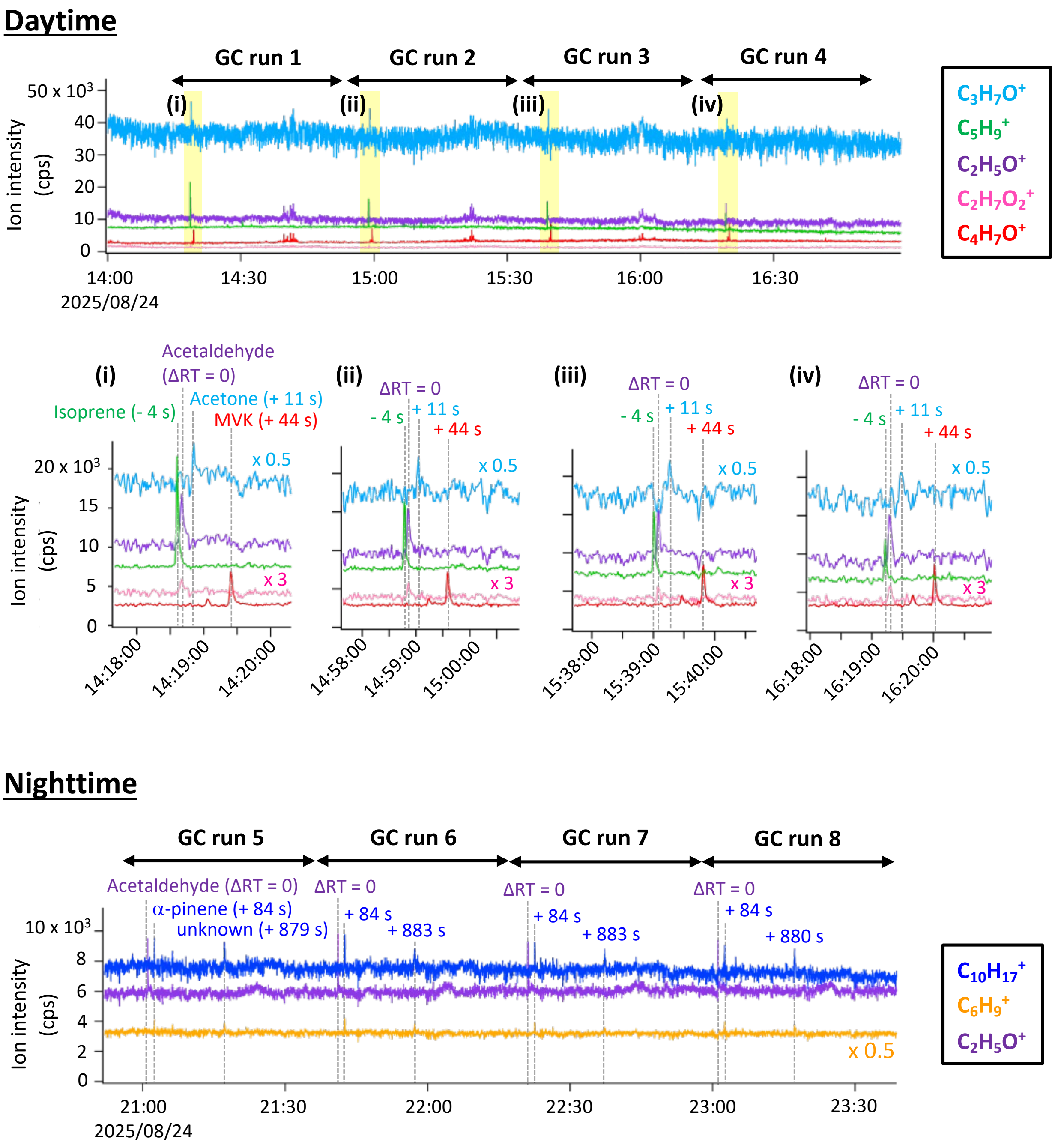


**Figure S4. GC/PTR-ToF-MS results of canopy-top atmospheres during daytime and nighttime on 24th August 2025.** Time courses of selected ions are shown (daytime: $C_3H_7O^+$, $C_5H_9^+$, $C_2H_5O^+$, $C_2H_7O_2^+$, and $C_4H_7O^+$; nighttime: $C_{10}H_{17}^+$, $C_6H_9^+$, and $C_2H_5O^+$). To assess the reproducibility of retention times, quadruplicate GC runs were performed separately during daytime (run 1-4) and nighttime (run 5-8). Panels (i)-(iv) are enlarged views of selected time segments from run 1-4 (highlighted in yellow). ΔRT denotes the difference in retention time for acetaldehyde detected as $C_2H_5O^+$. Based on ΔRT values from the forest air samples and authentic standards, compounds were assigned.

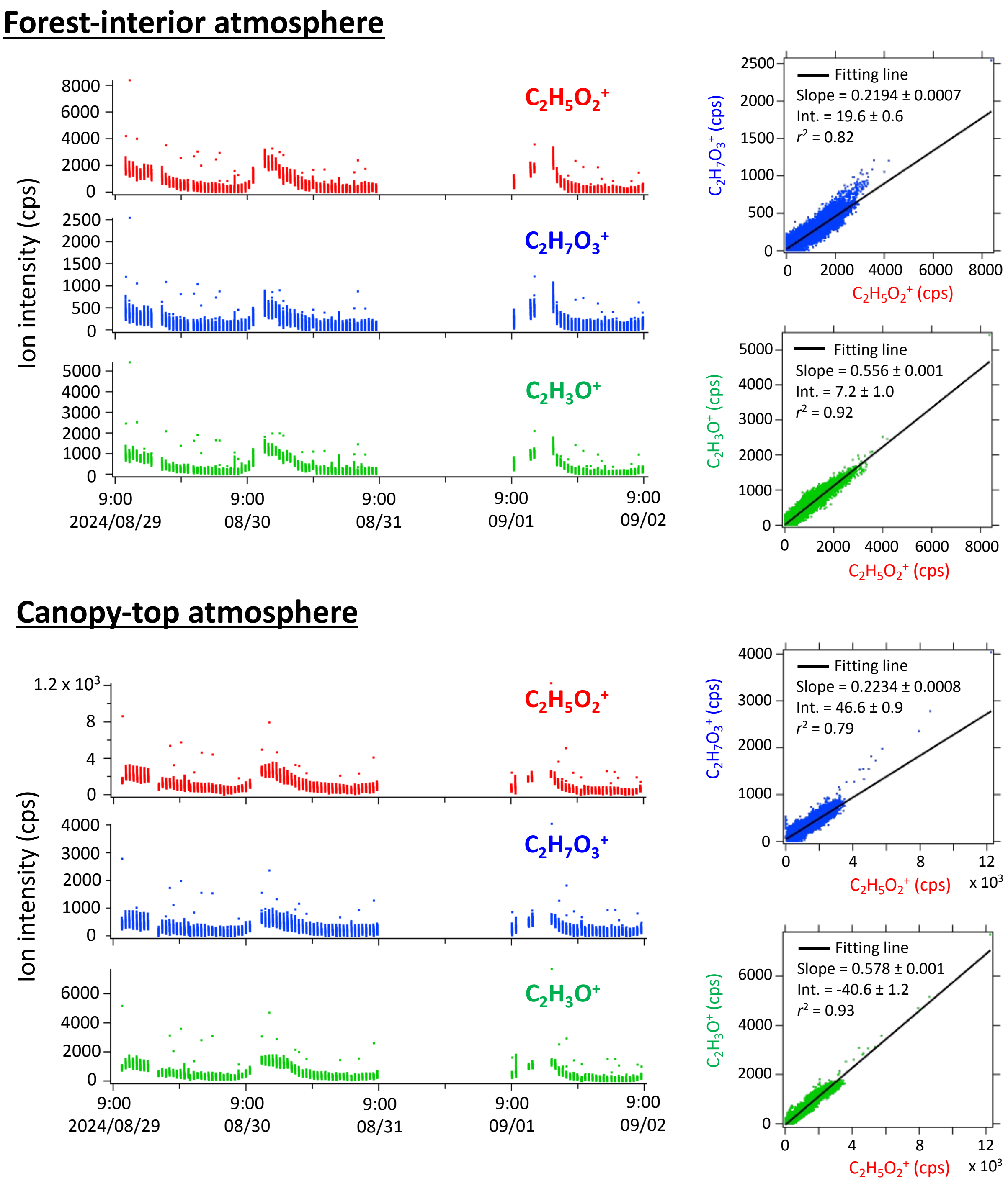


**Figure S5. Correlation among $C_2H_5O_2^+$, $C_2H_7O_3^+$, and $C_2H_3O^+$ in the 2024 dataset.** Time courses of $C_2H_5O_2^+$, $C_2H_7O_3^+$, and $C_2H_3O^+$ (left panels) and scatter plots of $C_2H_5O_2^+$ versus $C_2H_7O_3^+$ and $C_2H_3O^+$ (right panels) for the forest-interior and canopy-top atmospheres. Data points from the time courses (left panels) were used to derive the regression lines and correlation coefficients shown in the scatter plots (right panels).

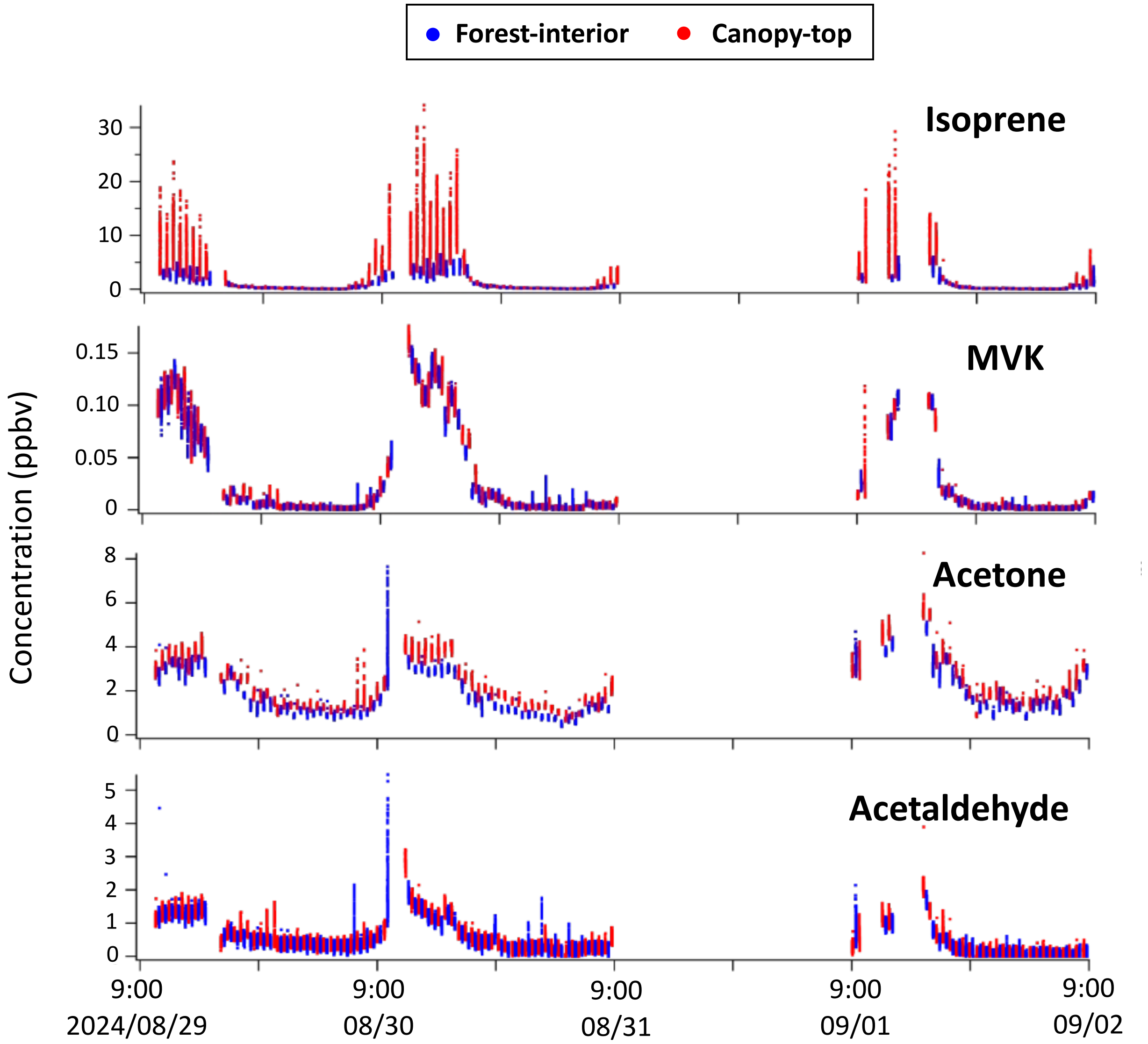


**Figure S6. Time courses of concentrations of eight VOCs in the 2024 dataset.** As noted in the main text, MVK concentrations may reflect not only forest emissions but also artifact formation within the instrument.

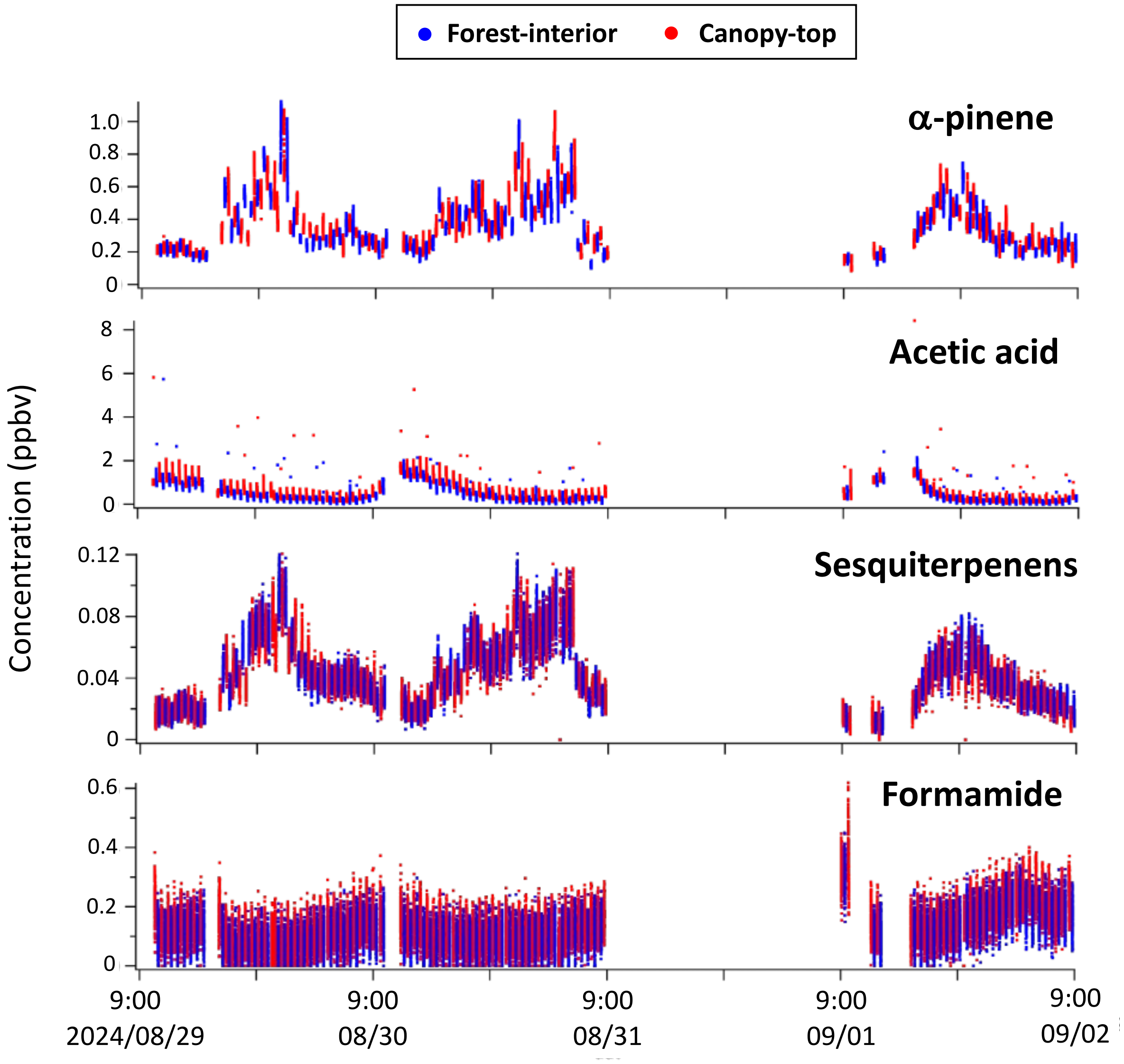


**Figure S6. Continued.**

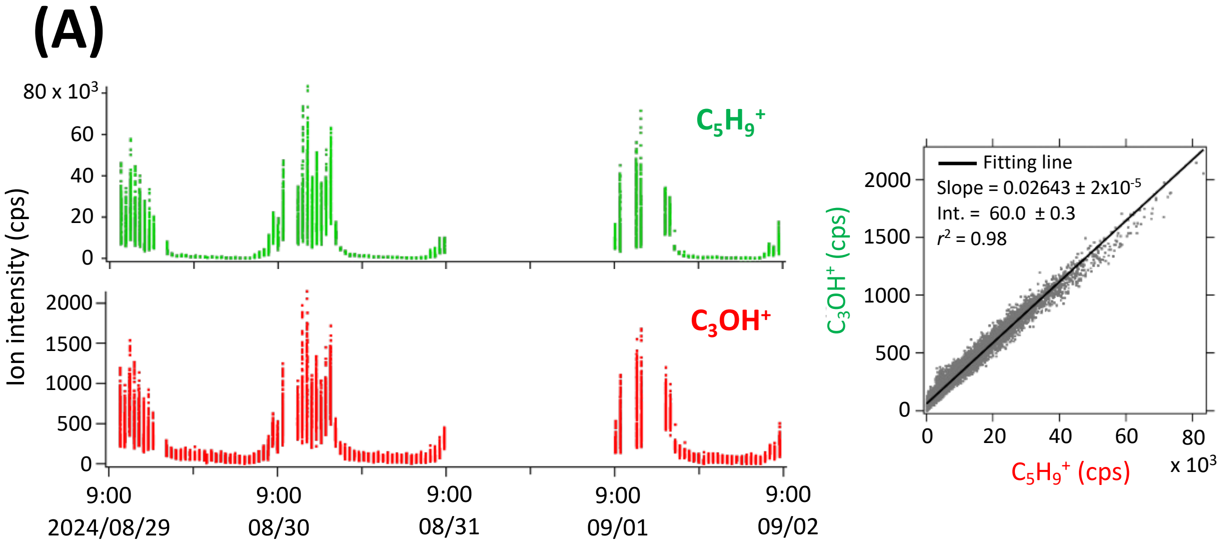


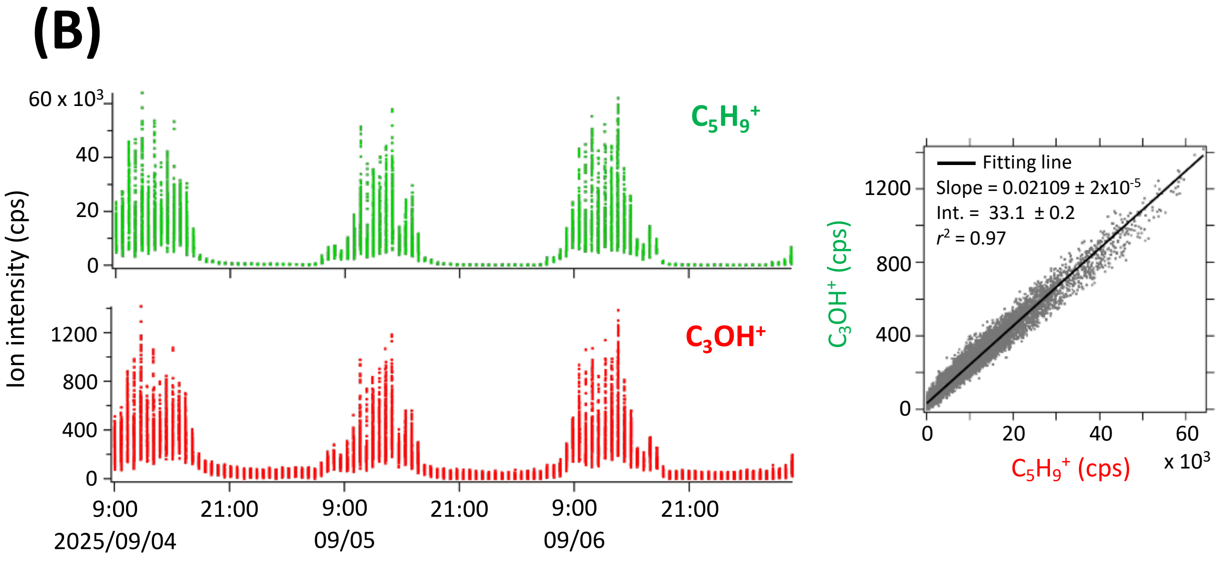


**Figure S7. Correlation between $C_3OH^+$ and $C_5H_9^+$ (isoprene).** Time courses of $C_3OH^+$ and $C_5H_9^+$ (left panels) and scatter plots of $C_3OH^+$ versus $C_5H_9^+$ (right panels) for the (A) 2024 and (B) 2025 datasets. Data points from the time courses (left panels) were used to derive the regression lines and correlation coefficients shown in the scatter plots (right panels).

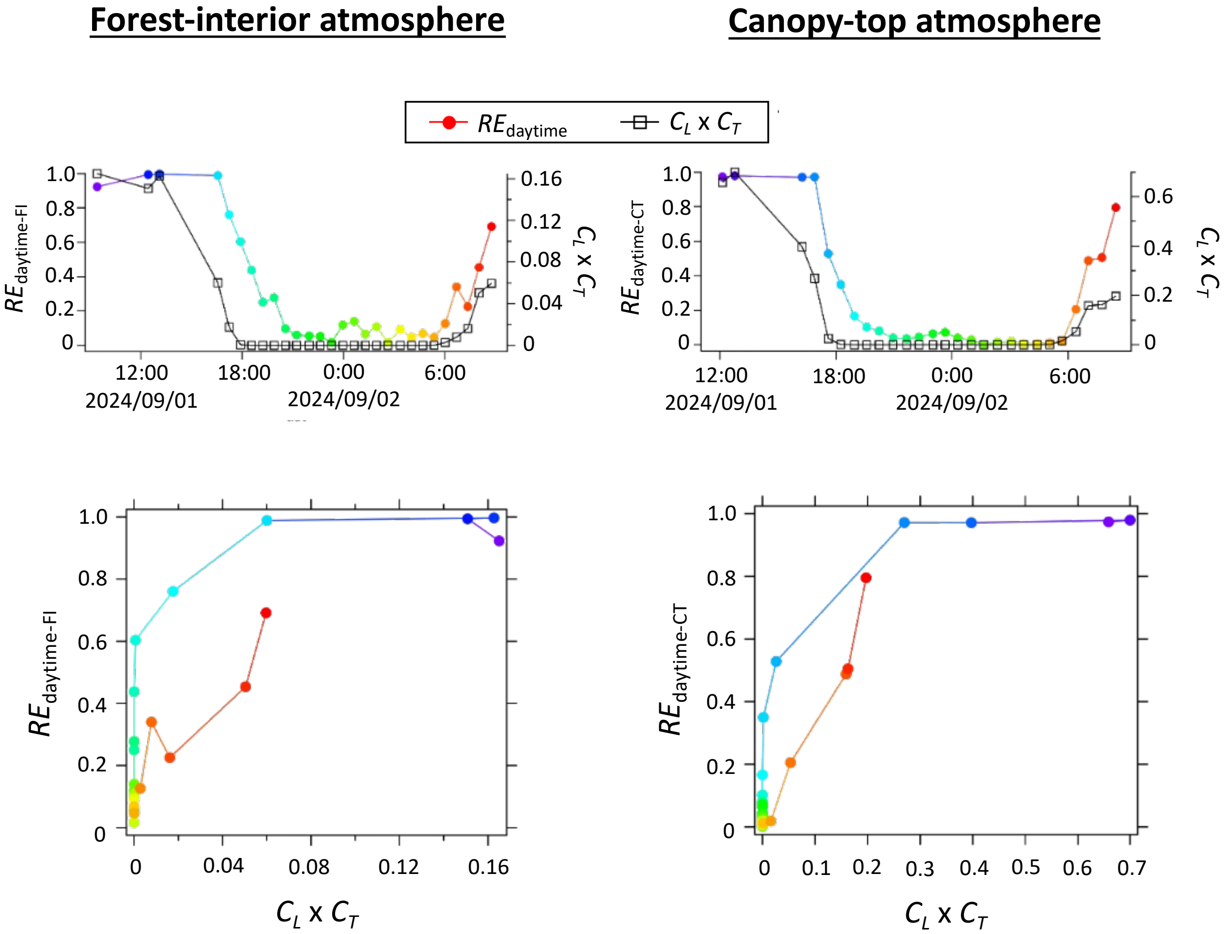


**Figure S8. Relationship between $RE_{daytime\text{-}FI}$, $RE_{daytime\text{-}CT}$, and $C_L \times C_T$ for the period from September 1 to 2, 2024.** Time courses of $RE_{daytime\text{-}FI}$, $RE_{daytime\text{-}CT}$, and $C_L \times C_T$ (upper panels), and scatter plots of $RE_{daytime\text{-}FI}$ or $RE_{daytime\text{-}CT}$ versus $C_L \times C_T$ (lower panels). Color codes in the lower panels indicate time of day and correspond to those used for $RE_{daytime}$ in the upper panels.

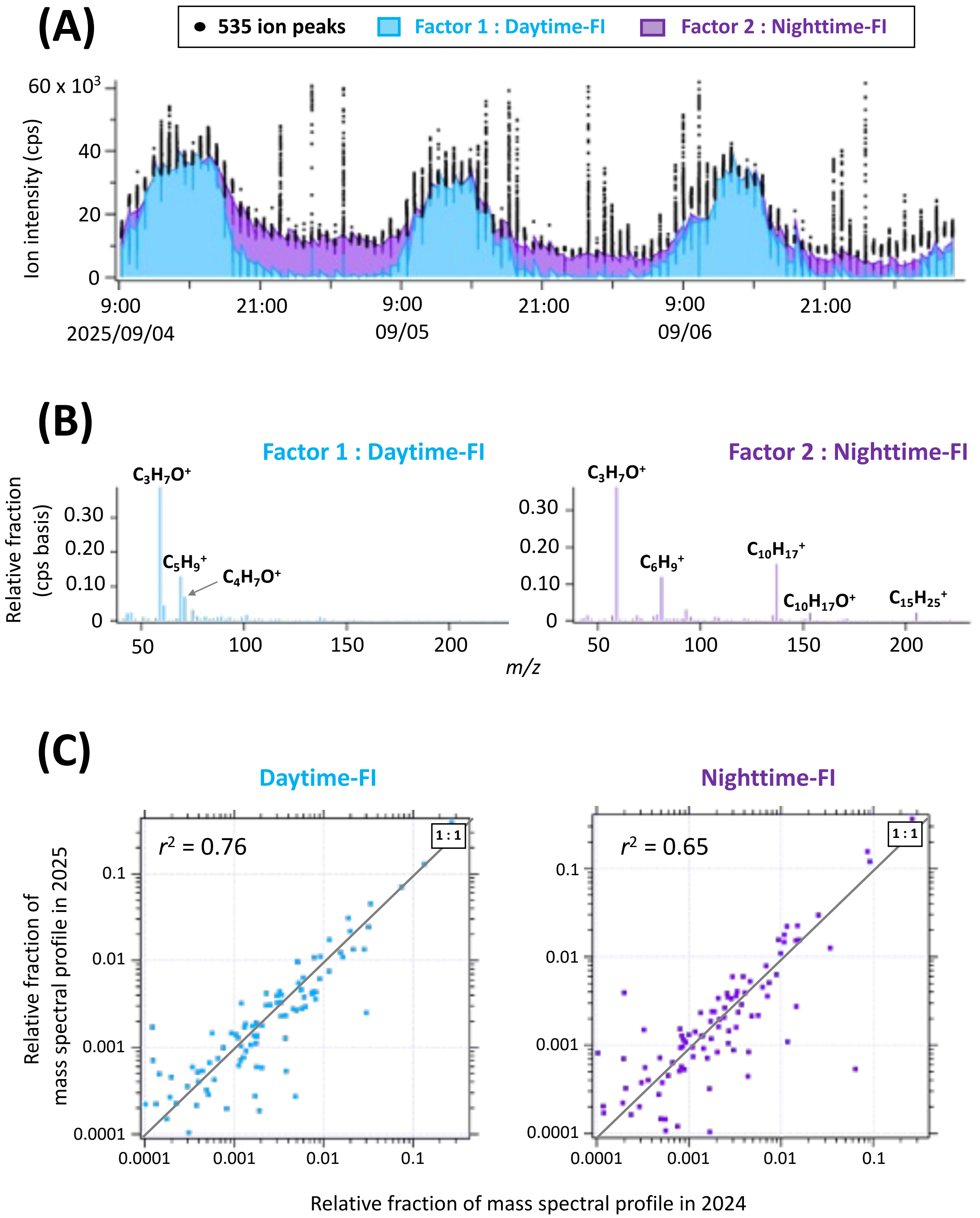


**Figure S9. PMF results for the two-factor solution applied to VOC ion signals measured in forest-interior atmospheres during three rain-free days in the 2025 datasets.** (A) Time courses of 535 measured ion signals and PMF fits. Individual PMF fits (blue and purple) are shown as stacked rather than overlapped. (B) Mass spectral profiles of the daytime-FI and nighttime-FI factors (FI denotes forest-interior). (C) Comparison of mass spectral profiles obtained in 2025 and 2024. Correlation coefficients ($r^2$) were calculated based on relative ion fractions evaluated on a logarithmic scale.

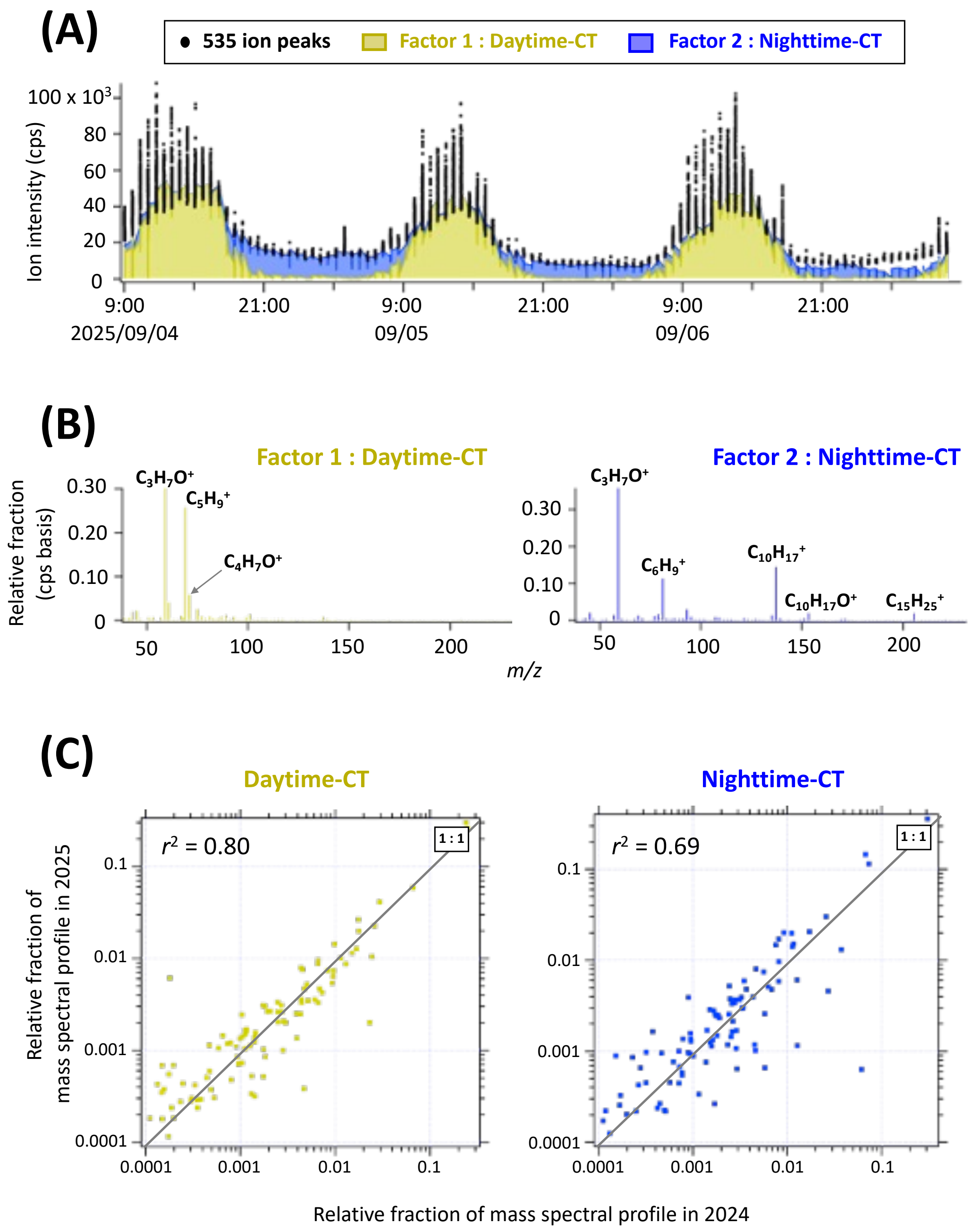


**Figure S10. PMF results for the two-factor solution applied to VOC ion signals measured in canopy-top atmospheres during three rain-free days in the 2025 datasets.** (A) Time-courses of 535 measured ion signals and corresponding PMF fits. Individual PMF fits (yellow and dark blue) are shown as stacked rather than overlapped. (B) Mass spectral profiles of the daytime-CT and nighttime-CT factors (CT denotes canopy-top). (C) Comparison of mass spectral profiles obtained in 2025 and 2024. Correlation coefficients ($r^2$) were calculated based on relative ion fractions evaluated on a logarithmic scale.

### Forest-interior atmosphere

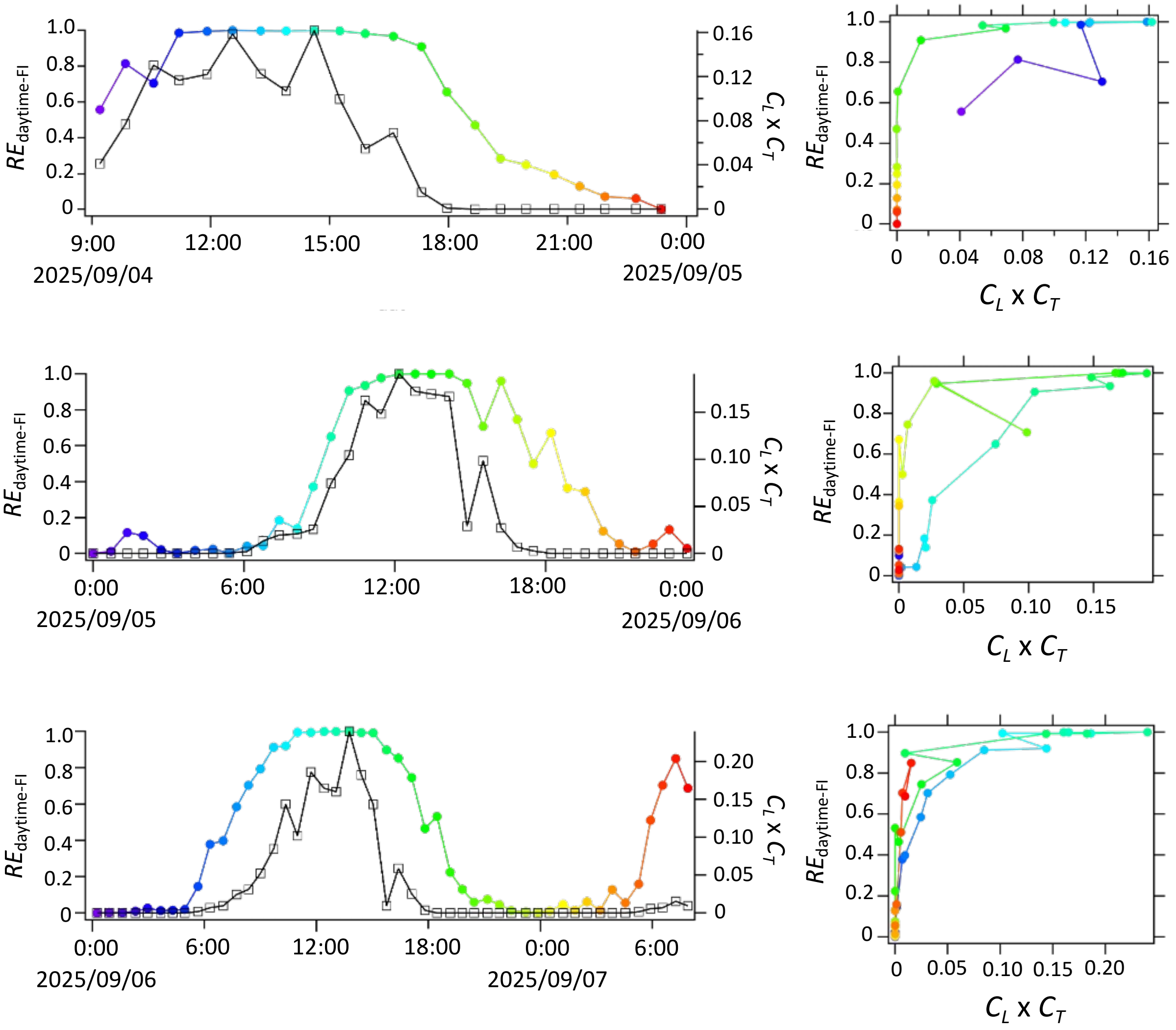


**Figure S11. Relationship between $RE_{\text{daytime-FI}}$, $RE_{\text{daytime-CT}}$, and $C_L \times C_T$ for the period from September 4 to 7, 2024.** Time courses of $RE_{\text{daytime-FI}}$, $RE_{\text{daytime-CT}}$, and $C_L \times C_T$ (left panels), and scatter plots of $RE_{\text{daytime-FI}}$ or $RE_{\text{daytime-CT}}$ versus $C_L \times C_T$ (right panels). Color codes in the right panels indicate time of day and correspond to those used for $RE_{\text{daytime}}$ in the left panels.

### Canopy-top atmosphere

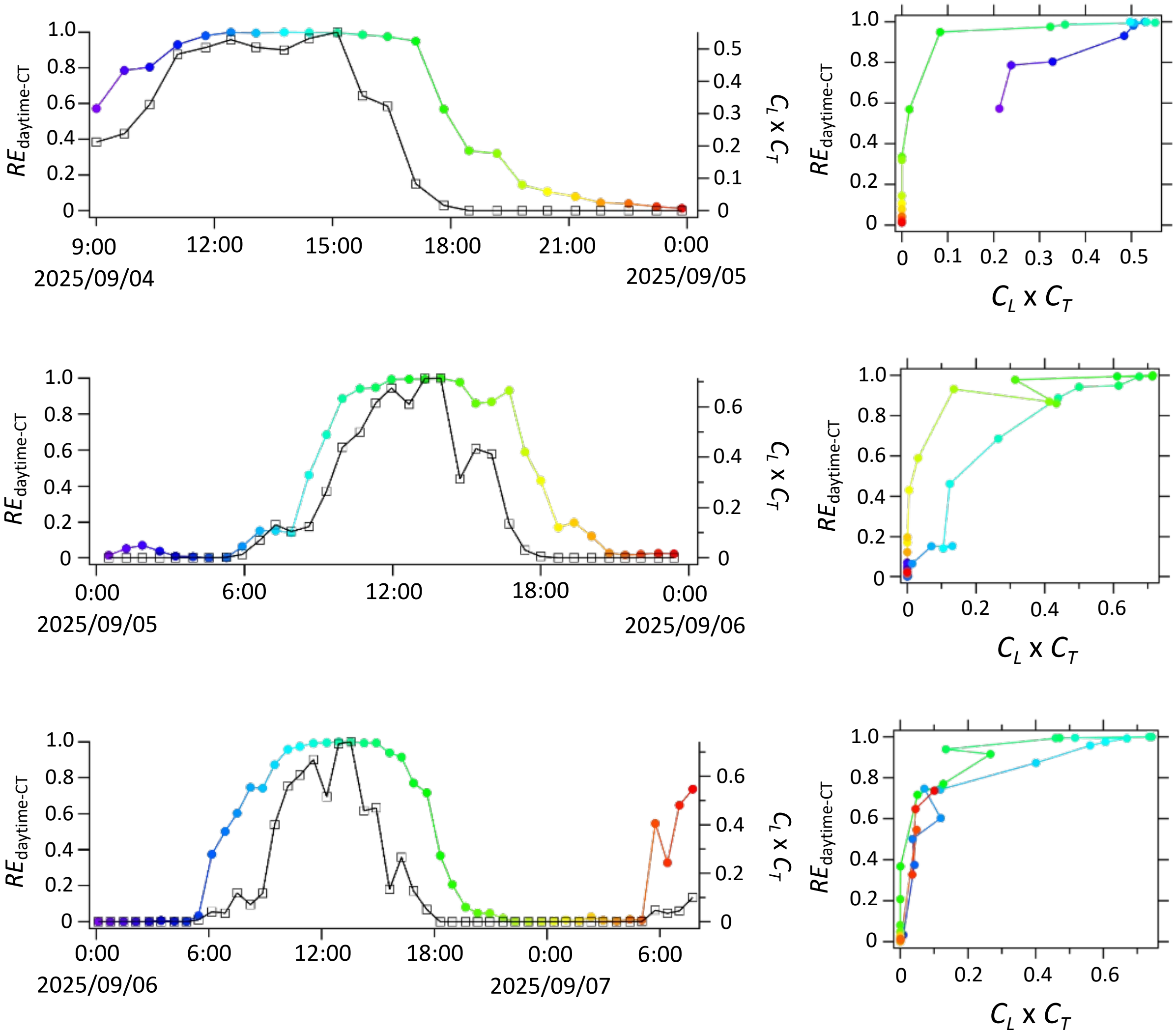


**Figure 11. continued.**
